# Mechanisms of Nuclear Pore Complex disassembly by the mitotic Polo-Like Kinase 1 (PLK-1) in *C. elegans* embryos

**DOI:** 10.1101/2023.02.21.528438

**Authors:** Sylvia Nkombo Nkoula, Griselda Velez-Aguilera, Batool Ossareh-Nazari, Lucie Van Hove, Cristina Ayuso, Véronique Legros, Guillaume Chevreux, Laura Thomas, Géraldine Seydoux, Peter Askjaer, Lionel Pintard

## Abstract

The nuclear envelope, which protects and organizes the interphase genome, is dismantled during mitosis. In the *C. elegans* zygote, nuclear envelope breakdown (NEBD) of the parental pronuclei is spatially and temporally regulated during mitosis to promote the unification of the parental genomes. During NEBD, Nuclear Pore Complex (NPC) disassembly is critical for rupturing the nuclear permeability barrier and removing the NPCs from the membranes near the centrosomes and between the juxtaposed pronuclei. By combining live imaging, biochemistry, and phosphoproteomics, we characterized NPC disassembly and unveiled the exact role of the mitotic kinase PLK-1 in this process. We show that PLK-1 disassembles the NPC by targeting multiple NPC sub-complexes, including the cytoplasmic filaments, the central channel, and the inner ring. Notably, PLK-1 is recruited to and phosphorylates intrinsically disordered regions of several multivalent linker nucleoporins, a mechanism that appears to be an evolutionarily conserved driver of NPC disassembly during mitosis. (149/150 words)

**One-Sentence Summary:** PLK-1 targets intrinsically disordered regions of multiple multivalent nucleoporins to dismantle the nuclear pore complexes in the *C. elegans* zygote.

## Introduction

During each round of cell division eukaryotic cells undergo a profound reorganization: the replicated DNA condenses into individual chromosomes, and the nuclear envelope, which protects and organizes the genome, breaks down during mitosis (*1, 2*). This step is critical in order for microtubules to access kinetochores and form the mitotic spindle. How NEBD is regulated remains incompletely understood, particularly during development.

The nuclear envelope is a double membrane composed of an outer and an inner phospholipid bilayer. Tethered to the inner nuclear membrane are the nuclear lamina and lamina-associated proteins which contribute to the mechanical integrity of the nucleus and organization of the genome during interphase. Transport across the two membranes occurs exclusively through nuclear pore complexes (NPCs), composed of roughly 30 evolutionarily conserved nucleoporins (NUPs), present in 8-32 copies (*3, 4*).

In contrast to vertebrate cells, where NEBD occurs entirely in prophase, NEBD in the semi-open mitosis of the *Caenorhabditis elegans* embryo is regulated both spatially and temporally (*5–9*). The first reported sign of NEBD in the zygote is the deformation of the nuclear envelope in the vicinity of centrosomes and the rupture of the NPC permeability barrier, detectable by the presence of soluble tubulin in the pronuclear space (*10*). NPC disassembly, manifested by loss of some nucleoporins from the nuclear envelope and lamina depolymerization, starts in the vicinity of the centrosomes, and progresses to the juxtaposed pronuclear envelopes located between the parental chromosomes (*5, 6, 11, 12*). However, other nucleoporins and the lamina persist on the remnant envelopes surrounding the mitotic spindle until the metaphase-to-anaphase transition (*5–7, 11, 12*). After lamina depolymerization is completed, the proper mingling of both parental chromosome sets on the metaphase plate requires the formation of a membrane scission event (also called a membrane gap) (*13*).

The mitotic Polo-like kinase PLK-1 kinase plays a prominent role in NEBD in the early embryo (*7, 11, 14, 15*). Consistently, PLK-1 is dynamically recruited to the nuclear envelope in prophase just before NEBD (*7*). PLK1 also localizes to the nuclear envelope in human cells (*16*) where it contributes to NPC disassembly together with NIMA and CyclinB-Cdk1 kinases (*16–18*). In particular, PLK1 is recruited to and phosphorylates NUP53, a structural subunit of the nuclear pore in human cells (*16*). PLK-1 likely promotes NPC disassembly in *C. elegans,* but its critical targets in this process are currently unknown. Beyond NPC disassembly, PLK-1 triggers lamina depolymerization in *C. elegans* by directly phosphorylating the lamina (*11*). In fact, expression of a *lmn-1*^8A^ allele, mutated on eight PLK-1 phosphorylation sites (*19*), is sufficient to prevent lamina depolymerization, membrane gap formation, and the fusion of the parental genomes during mitosis (*10, 11*). Whether stabilization of the lamina affects the disintegration of the NPC is not known. Some nucleoporins and the lamina accumulate at the NE upon *plk-1* inactivation (*15*) suggesting that PLK- 1 might regulate both lamina depolymerization and NPC disassembly. Alternatively, as the lamina interacts physically with nucleoporins (*20–22*), lamina persistence could alter NPC disassembly. The *lmn-1*^8A^ allele represents a unique genetic tool to test these possibilities.

Here, by combining live-imaging of fluorescently tagged nucleoporins with biochemical and phosphoproteomic approaches, we characterize the steps and mechanism of NPC disassembly in *C. elegans* embryos. We show that PLK-1 triggers NPC disassembly independently of lamina depolymerization, and we identify its critical targets in this process. PLK-1 phosphorylates intrinsically disordered regions of multivalent linker nucleoporins notably NPP-14^NUP214^, NPP- 10N^NUP98^ and NPP-19^NUP53^ to dismantle the NPC.

## Results

### PLK-1 triggers the disassembly of soluble nucleoporins independently of lamina depolymerization

Nuclear envelope components are conserved in *C. elegans,* including the lamina, encoded by a single gene *lmn-1* (B-type lamin) (*23*) and most of the nucleoporins, although the percentage of identity between human (NUP) and *C. elegans* (NPP) nucleoporins is not higher than 25% (*9, 24*) (**Fig. 1A**). Within the nuclear pores, nucleoporins assemble into modular subcomplexes organized as a three- ring stacked structure: one inner ring (in orange on **Fig. 1A**) and two outer rings, containing the Y- complexes (green), located on the cytoplasmic and nuclear membrane (*3, 4, 25*). The inner ring and the Y-complexes, which are anchored to the membrane by interacting with transmembrane nucleoporins (brown) and the lipid bilayer, serve as building blocks for the scaffolding of nuclear pores **(Fig. 1A)**. Attached to these rings are nucleoporins of the central channel (light blue) and peripheral cytoplasmic (dark blue) and nucleoplasmic structures (red) responsible for the translocation of substrates through the NPCs (**Fig. 1A**). Nucleoporins with Phenylalanine-Glycine (FG) repeats line the central channel and serve a dual role: they form the permeability barrier by restricting the passive diffusion of macromolecules larger than ∼40 kDa, and they provide binding sites for nuclear transport receptors (*26, 27*).

**Fig. 1:**
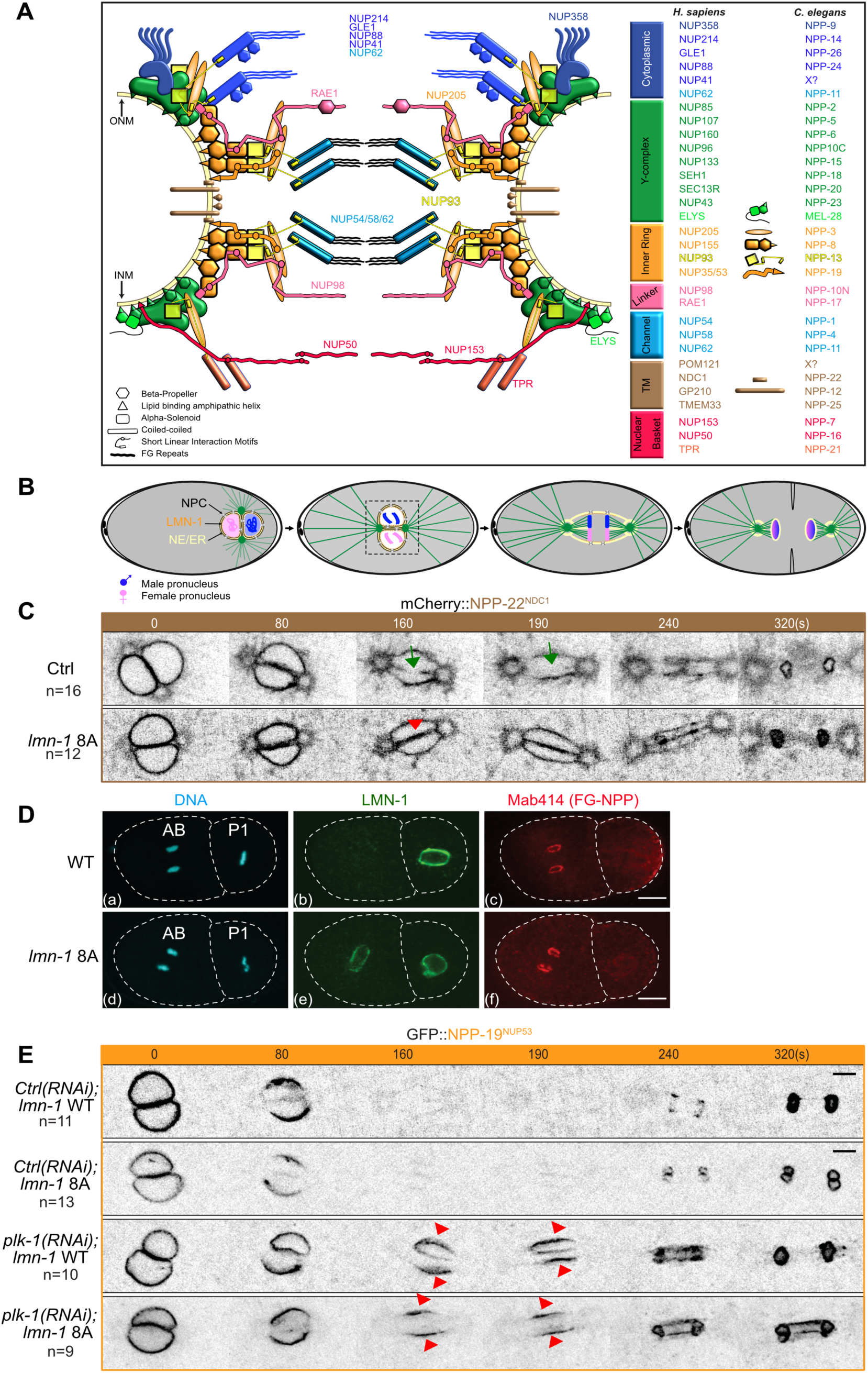
PLK-1 triggers nuclear pore complexes disassembly independently of lamina depolymerization. **A-** Schematic representation of the nuclear pore complex architecture and nomenclature of the nucleoporins in human cells (NUP) and in *C. elegans* (NPP). **B-** Schematics illustrate the division of the one-cell *C. elegans* embryo. After meeting of the female (pink) and male (blue) pronuclei at the posterior pole, the nucleo-centrosomal complex undergoes a 90° rotation to align along the anteroposterior axis of the embryo. Parental chromosomes congress on the metaphase plate before the initiation of anaphase. In telophase, the nuclear envelope reforms around decondensing parental chromosomes that are now mixed in a single nucleus. During mitosis, remnant nuclear envelope (yellow) remains around the segregating chromosomes. This remnant NE contains integral membrane proteins. NPC: Nuclear Pore Complexes, NE/ER: nuclear envelope/endoplasmic reticulum. **C-** Spinning disk confocal micrographs of wild-type or *lmn-1* 8A mutant embryos expressing endogenously tagged GFP::NPP-22^NDC1^ in mitosis. The green arrow marks the presence of a membrane gap in control wild-type as opposed to *lmn-1* 8A embryos (red arrow). Timings in seconds are relative to pronuclei juxtaposition (0s). All panels are at the same magnification. Scale Bar, 10 μm. n= number of embryos analyzed in this and other Figures. **D-** Confocal images of fixed wild-type and *lmn-1* 8A mutant two-cell embryos stained with LMN-1 (green) and Mab414 (red) antibodies and counterstained with DAPI (magenta). All panels are at the same magnification. Scale Bar, 10 μm. FG-NPP: Nucleoporins with FG repeats detected by the Mab414 antibody. **E-** Spinning disk confocal micrographs of live wild-type or *lmn-1* 8A embryos expressing endogenously tagged GFP::NPP-19^NUP53^ in mitosis exposed to control, or *plk-1*(*RNAi*). The red arrowheads point to GFP::NPP-19^NUP53^ persisting on the nuclear envelope during mitosis. Timings in seconds are relative to pronuclei juxtaposition (0s). All panels are at the same magnification. Scale Bar, 10μm.

To investigate whether lamina stabilization impacts NPC disassembly, we analyzed the dynamics of the transmembrane nucleoporin NPP-22^NDC1^, endogenously tagged with mCherry, in wild-type versus *lmn-1* 8A mutant one-cell embryos during mitosis using spinning disk confocal microscopy (**Fig. 1B, 1C**). mCherry::NPP-22^NDC1^ localized exclusively to the nuclear envelope of the male and female pronuclei during their migration in wild-type embryos (**Fig. 1C**) (*10, 28*). Then, as the embryos progressed through mitosis, it was progressively excluded from the region between the juxtaposed pronuclei, starting where the characteristic pronuclear membrane scission (gap) event occurs (**Fig. 1C, green arrows**) (*13*), and accumulated on the endoplasmic reticulum surrounding the centrosomes, recently named the “centriculum” (*29*). By contrast, in *lmn-1* 8A mutant embryos, where the lamina is not dismantled during mitosis, mCherry::NPP-22^NDC1^ also persisted on the juxtaposed pronuclear envelopes (**Fig. 1C, red arrowhead**). Thus, stabilization of the lamina can influence the dynamics of the transmembrane nucleoporin NPP-22^NDC1^ during mitosis.

Next, we monitored nucleoporins with glycosylated FG repeats (*30*) using the Mab414 antibodies in indirect immunofluorescence experiments. The asynchronous mitoses of the two-cell stage embryo (**Fig. 1D, a, d**) allow a direct comparison of nuclear envelope components behavior in metaphase (the P1 cell, right) and telophase (the AB cell, left). In wild type embryos, FG nucleoporins were undetectable in metaphase (P1 blastomere) (**Fig. 1D, c**), while the lamina still localized around the DNA (**Fig. 1D, b**) indicating that nucleoporins disassemble before lamina depolymerization. During nuclear envelope reformation, the FG nucleoporin signal reaccumulated around the decondensing DNA in telophase in the AB blastomere (**Fig. 1D, c**), while the lamina had not yet polymerized (**Fig. 1D, b**). In *lmn-1 8A* embryos, where lamina persist during mitosis (**Fig1D, e**), FG nucleoporins presented an identical localization as in wild-type embryos (**Fig. 1D, f**), indicating that the persistence of the lamina at the nuclear envelope does not affect their disassembly or reassembly during mitosis. To corroborate these observations, we filmed one-cell embryos expressing endogenously tagged GFP::NPP-19^NUP53^, a core subunit of the inner ring complex (*22, 31–36*). GFP::NPP-19^NUP53^ disappeared from the nuclear envelope with the same kinetics in wild type and *lmn-1* 8A mitotic embryos confirming that the persistence of the lamina does not affect the disassembly of soluble nucleoporins (**Fig. 1E**). However, GFP::NPP-19^NUP53^ readily persisted on the nuclear envelope throughout mitosis upon *plk-1* inactivation, both in wild-type and in *lmn-1* 8A embryos (**Fig. 1E,** red arrowheads).

Taken together, these observations indicate that PLK-1 promotes disassembly of some nucleoporins independently of lamina depolymerization, although forcing maintenance of the lamina is sufficient to alter the dynamics of the transmembrane nucleoporin NPP-22^NDC1^ (**Fig. 1C**).

### PLK-1 targets the cytoplasmic filaments, the inner ring complex and the central channel nucleoporins

To identify the nuclear pore subcomplexes targeted by PLK-1, we used spinning disk confocal microscopy to monitor the dynamics of fluorescently tagged nucleoporins from each NPC subcomplex during mitosis in control embryos, or upon *plk-1* inactivation. Since PLK-1 regulates several processes in the early embryo (*7, 15, 37–43*), we used partial RNAi to exclude indirect effects, reasoning that direct PLK-1 targets should be the most sensitive to mild *plk-1* inactivation.

We filmed embryos expressing fluorescently tagged nucleoporins of the cytoplasmic filaments (GFP::NPP-9^NUP358^, GFP::NPP-24^NUP88^), the Y-complex (GFP::NPP-5^NUP107^, GFP::MEL-28^ELYS^), the inner ring (superfolder (s)GFP::NPP-13^NUP93^, mCherry::NPP-8^NUP155^), the central channel (NPP- 1^NUP54^::GFP and GFP::NPP-11^NUP62^), and the nuclear basket (sGFP::NPP-7^NUP153^ and NPP- 21^TPR^::GFP). We also monitored the dynamics of NPP-10N^NUP98^ (tagged with NeonGreen), the *C. elegans* counterpart of NUP98, which belongs to multiple subcomplexes in human cells (*4, 44*), and possibly also in *C. elegans*. Some nucleoporins were endogenously tagged with fluorescent markers using CRISPR/Cas9 whereas others were expressed from single-copy transgenes (Material and Methods).

At time 0, defined as when the two pronuclei are juxtaposed and centered in the middle of the embryos, all the nucleoporins we monitored localized to the nuclear envelope (**Fig. 2 C, D, E, F**), except nucleoporins of the nuclear basket and of the Y-complex (**Fig. 2, A, B**). The basket nucleoporins GFP::NPP-7^NUP153^ and GFP::NPP-21^TPR^ localized to the nucleoplasm (**Fig. 2A**) while the Y-complex subunits GFP::NPP-5^NUP107^ and GFP::MEL-28^ELYS^ had begun to accumulate at the kinetochores (**Fig. 2B**). Partial *plk-1* inactivation did not noticeably affect the dynamics of these nucleoporins during mitosis. Both GFP::NPP-5^NUP107^ and GFP::MEL-28^ELYS^ localized to the kinetochores as in control embryos (**Fig. 2A and 2B**).

**Fig. 2:**
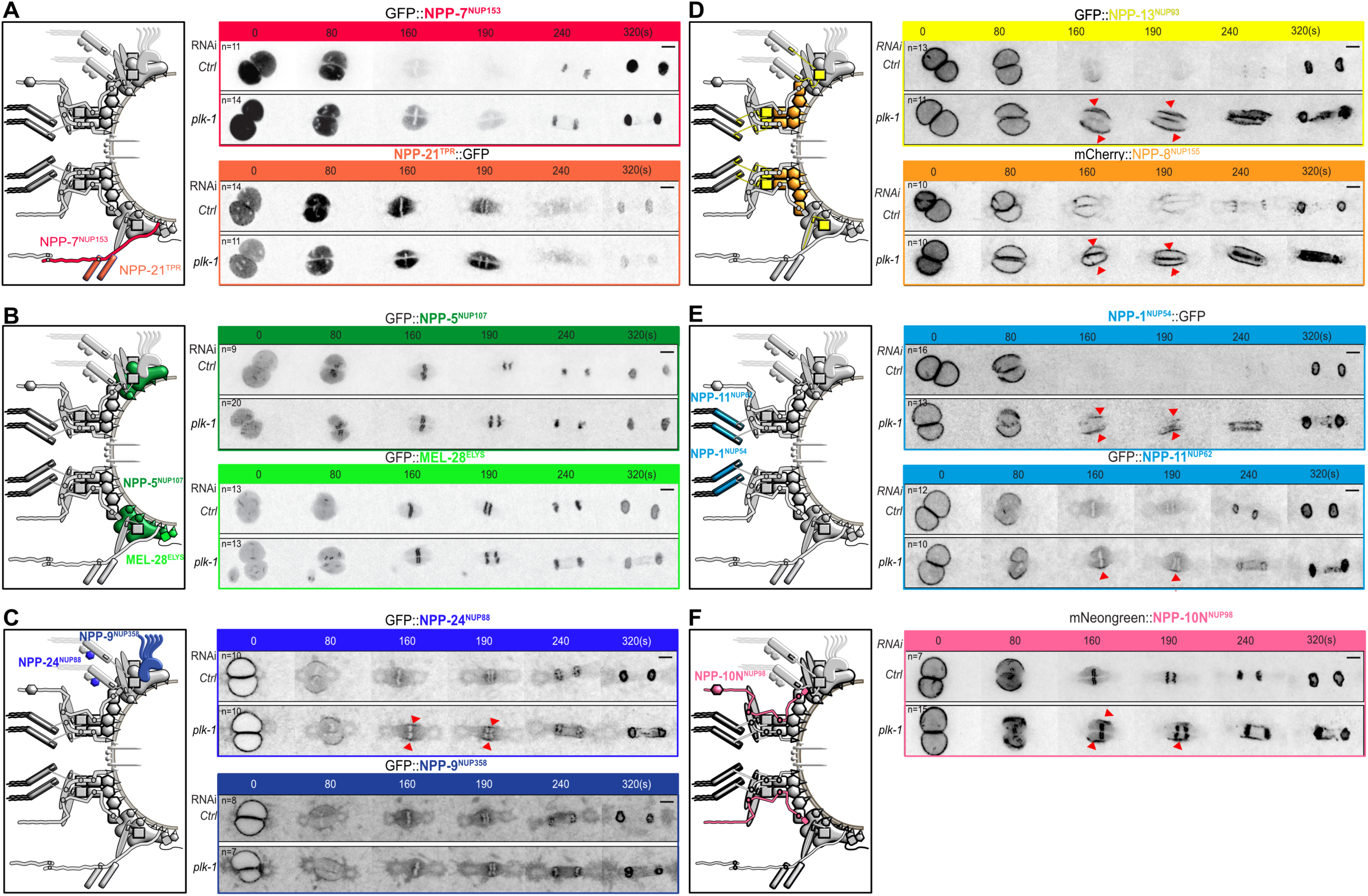
Nucleoporins of the cytoplasmic filaments, the central channel and the inner ring accumulate at the nuclear envelope upon *plk-1* inactivation A-F: Spinning disk confocal micrographs of embryos expressing the indicated tagged nucleoporins exposed to control, or *plk-1(RNAi)* starting when the pronuclei are juxtaposed, centered and aligned along the anteroposterior axis of the embryo (time 0). The red arrowheads show the persisting nucleoporins on the nuclear envelope during mitosis. All panels are at the same magnification. Scale Bar, 10μm.

On the cytoplasmic face of the NPC, NUP88 forms a complex with NUP214 (*44*). As shown **Fig. 2C**, *plk-1* inactivation affected the dynamics of GFP::NPP-24^NUP88^. While this nucleoporin relocalized to the centriculum and the mitotic spindle area during mitosis in control and *plk-1(RNAi)* embryos, a fraction persisted on the remnant envelope surrounding the mitotic spindle during the metaphase-to- anaphase transition upon reduction of *plk-1* function (**Fig. 2C**). OLLAS::NPP-14^NUP214^ similarly persisted on the nuclear envelope remnants upon partial inactivation of *plk-1* during mitosis (**Fig. S1**), indicating that PLK-1 controls the disassembly of the cytoplasmic filaments. GFP::NPP-9^NUP358^, which also localizes to the cytoplasmic face the NPC, was however unaffected upon *plk-1* inactivation (**Fig. 2C**).

Next, we monitored the dynamics of the nucleoporins of the inner ring complex. Within this complex, in other systems, the linker nucleoporin NUP53 has a scaffolding role and binds NUP155, NUP93 and NUP205 (*22, 32–36*). Previous immunofluorescence studies in early *C. elegans* embryos showed that NPP-3^NUP205^ and NPP-13^NUP93^ localize to the nuclear envelope before NEBD but then disappear first in the vicinity of the centrosomes (*6*). We confirmed these observations using endogenously tagged GFP::NPP-13^NUP93^. Before NEBD, when the two pronuclei are juxtaposed (time 0), GFP::NPP-13^NUP93^ localized to the nuclear envelope but after 160s, it was barely detectable at the nuclear envelope (**Fig. 2D**). However, upon *plk-1* partial inactivation, GFP::NPP-13^NUP93^ persisted on the nuclear envelope throughout mitosis (**Fig. 2D,** red arrowheads), similar to GFP::NPP-19^NUP53^ (**Fig. 1E**). mCherry::NPP-8^NUP155^ also persisted on the nuclear envelope throughout mitosis upon *plk- 1* inactivation, and failed to relocalize to the centriculum (**Fig. 2D**). Thus, three components of the inner ring complex: NPP-8^NUP155^, NPP-13^NUP93^ and NPP-19^NUP53^ persist on the nuclear envelope upon partial *plk-1* inactivation.

Within the inner ring complex, NPP-13^NUP93^ recruits the FG nucleoporins NPP-1^NUP54^, NPP-4^NUP58^ and NPP-11^NUP62^, which form a trimeric complex located in the central channel of the NPC (*7, 45*) (**Fig. 1A**). In control conditions, NPP-1^NUP54^::GFP localized to the nuclear envelope and was still detected on the remnant envelopes 80s after juxtaposition of the pronuclei. As the embryo progressed through mitosis, it became undetectable and reaccumulated on the nuclear envelope only in telophase (**Fig. 2E**). However, GFP::NPP-1^NUP54^ persisted on the nuclear envelope during mitosis upon partial depletion of PLK-1, consistent with previous observations (*15*). GFP::NPP-11^NUP62^, also persisted to the nuclear envelope upon *plk-1* inactivation (**Fig. 2E**). Thus, *plk-1* depletion also affects the disassembly of the central channel nucleoporins from the NPCs.

Finally, we analyzed the dynamics of NeonGreen::NPP-10N^NUP98^. In wild-type conditions, NeonGreen::NPP-10N^NUP98^ localized to the NE but relocalized to the kinetochores during mitosis. Upon PLK-1 depletion, NeonGreen::NPP-10N^NUP98^ also relocalized to the kinetochores but a significant fraction persisted on the nuclear envelope during mitosis (**Fig. 2F, red arrows**).

Taken together, these observations indicate that PLK-1 inactivation affects the disassembly of the cytoplasmic filaments (NPP-14^NUP214^, NPP-24^NUP88^), the inner ring complex (NPP-8^NUP155^, NPP- 13^NUP93^, NPP-19^NUP53,^ NPP-10N^NUP98^), and the central channel (NPP-1^NUP54^, NPP-11^NUP62^) during mitosis.

### PLK-1 recruitment to the nuclear envelope precedes the disassembly of filament, central channel and inner ring subcomplexes

The order and precise timing of NPC subcomplex disassembly has not yet been characterized in *C. elegans*. In human tissue culture cells, NUP98 is the first nucleoporin to leave the NPC during mitosis, causing the rupture of the nuclear permeability barrier (*17, 46*). Intriguingly, our observations suggest that nucleoporins of the Y complex and the nuclear basket leave the NPC well before NPP-10N^NUP98^, already during pronuclear migration and meeting (**Fig. 2**), well before NPP-10N^NUP98^ is removed, and possibly even before PLK-1::sGFP is recruited to the NE and the nuclear permeability barrier is ruptured.

To precisely characterize the order of NPC disassembly relative to PLK-1 recruitment to the NE, we used live-spinning disk confocal microscopy at high temporal resolution (1 image/2s). We quantified the levels of nucleoporins of the different subcomplexes and PLK-1::sGFP at the NE, relative to the rupture of the nuclear permeability barrier and anaphase onset (time 0). To determine when the NE becomes permeable, we quantified the amount of free mCherry::Histone, which is higher in the nucleoplasm than in the cytoplasm before NEBD but equilibrates between the two compartments once the NE becomes permeable (Material and Methods).

This analysis confirmed that nucleoporins of the basket and the Y-complex leave the nuclear envelope well before it becomes permeable (**Fig. 3A**). GFP::NPP-7^NUP153^ localized to the NE during pronuclear migration (-410s before anaphase, **Fig. S2A**, arrow) but had already relocalized to the nucleoplasm before pronuclear meeting. Likewise, NPP-21^TPR^::GFP had almost entirely relocalized to the nucleoplasm at the time of pronuclear meeting (**Fig. 3B, S2B**). The Y complex subunit GFP::MEL- 28^ELYS^ similarly disappeared from the nuclear envelope by pronuclei meeting and relocalized to the kinetochores, concomitantly with chromosome condensation (**Fig. 3B, S2C**). The other Y complex subunits GFP::NPP-5^NUP107^, GFP::NPP-6^NUP160^ and GFP::NPP-18^SEH1^ departed the NE slightly after GFP::MEL-28^ELYS^ but all behaved similarly (**Fig. 3B, S2D, E, F**), suggesting that the entire Y complex leaves the NE early in prophase to relocalize to the kinetochores. Quantifications of the signal at the pronuclear envelope revealed that GFP::NPP-7^NUP153^ is the first nucleoporin to leave the NE followed by GFP::MEL-28^ELYS^, NPP-21^TPR^::GFP and the other Y complex subunits (**Fig. 3B, S2G**). All these subunits leave the NE before the recruitment of PLK-1::sGFP, which starts to accumulate at the NE 250s before anaphase onset (**Fig. 3A**). PLK-1::sGFP accumulation at the NE precedes the departure of NPP-10N^NUP98^, NPP-24^NUP54^, NPP-1^NUP54^, and NPP-19^NUP53^. As showed in the **Fig. 3**, NPP-10N^NUP98^ leaves the NE slightly before the cytoplasmic filament nucleoporin NPP- 24^NUP88^ and the central channel nucleoporin NPP-1^NUP54^, which disappears from the NE before NPP- 19^NUP53^ (**Fig. 3**, **Fig. S3**). The NE becomes permeable only when NPP-10N^NUP98^ is removed from the NPC (**Fig. 3A**). More specifically, we observed a significant loss of permeability barrier when almost 60 % of NPP-10N was removed from the NE (**Fig. 3A**)

**Fig. 3:**
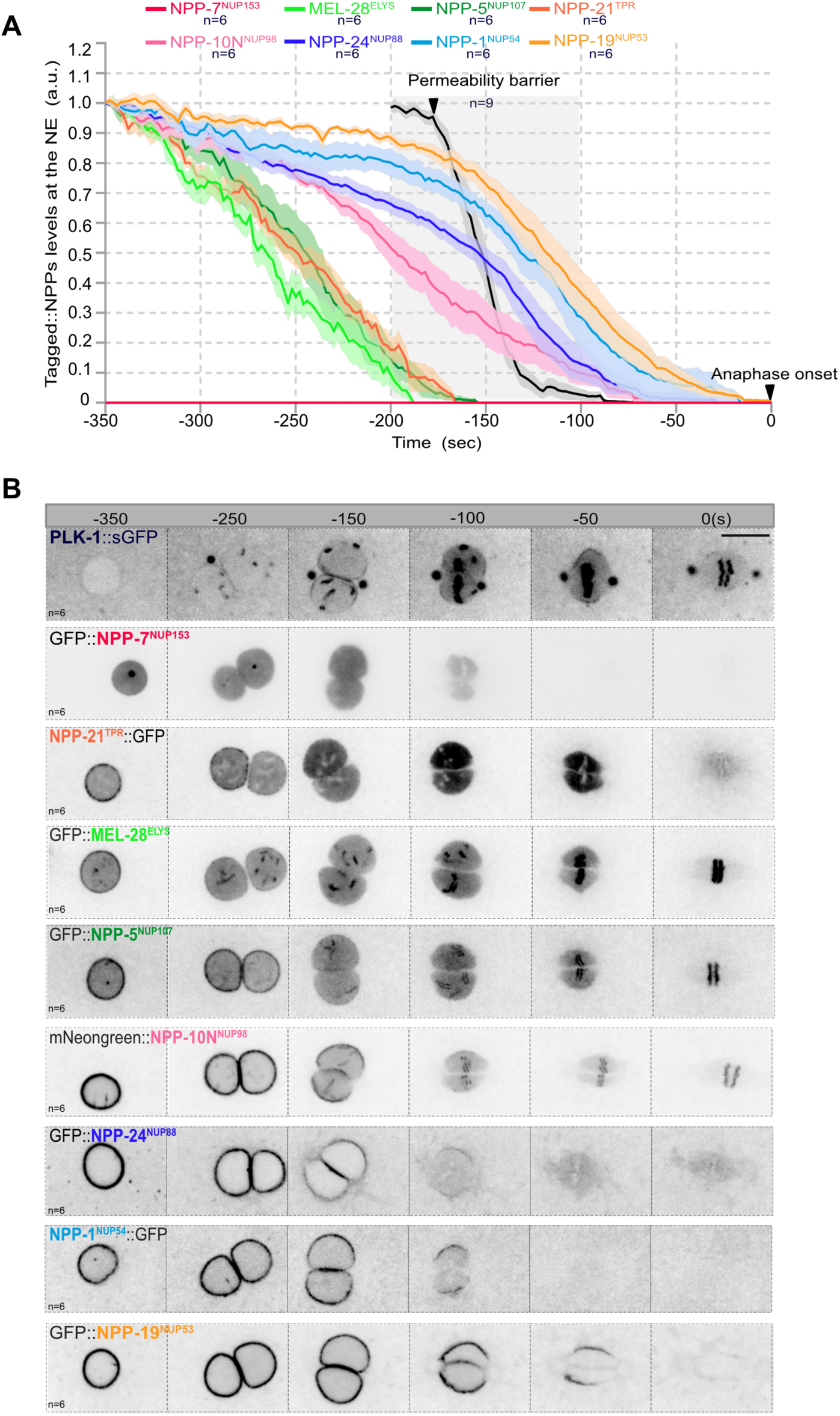
Timing of NPC subcomplexes disassembly in the *C. elegans* zygote relative to the rupture of the nuclear permeability barrier. **A-** Graph presenting the quantification of GFP::NPP signal intensity at the NE in the *C. elegans* zygote from pronuclear nuclear migration to anaphase onset (time 0). The average signal intensity of the fluorescently tagged NPPs at the NE 350s before anaphase was arbitrarily defined as 1. The mean +/- SEM is presented for n=6 embryos. Data were collected from three independent experiments. The dark line presents the quantification of the nuclear permeability barrier starting 200s before anaphase onset. **B-** Spinning disk confocal micrographs of one-cell embryos expressing PLK-1::sGFP or the indicated tagged nucleoporins from pronuclear migration and meeting to anaphase onset (time 0). All panels are at the same magnification. Scale Bar, 10μm.

Taken together, these data indicate that the Y complex and the nuclear basket nucleoporins leave the NE before PLK-1 recruitment, which precedes the departure of NPP-10N^NUP98^ and the rupture of the nuclear permeability barrier (**Fig. 3**).

### A biochemical screen for PLK-1 PBD-interacting phospho-nucleoporins

We next searched for the direct PLK-1 substrates at the nuclear pore complexes. PLK-1 uses its conserved C-terminal Polo-box domain (PBD) to bind its substrates phosphorylated on specific sequence motifs (aka Polo-docking sites [S-pS/pT-P/X]) that are created by other priming kinases (non-self-priming) or are self-primed by PLK-1 itself, thus providing an efficient mechanism to regulate PLK-1 subcellular localization and substrate selectivity in space and time (*47–51*). We previously showed that PLK-1 uses its PBD domain to localize to the nuclear envelope in the early *C. elegans* embryo where it binds nucleoporins of the central channel NPP-1^NUP54^, NPP-4^NUP58^ and NPP-11^NUP62^ phosphorylated on Polo-docking sites by PLK-1 and Cyclin B-Cdk1 (*7*).

To determine which nucleoporins are phosphorylated on Polo-docking sites *in vivo*, we analyzed the *C. elegans* embryos phosphoproteome using FeNTA affinity chromatography and liquid- chromatography-tandem mass spectrometry (LC-MS/MS) (**Fig. 4A**). While previous phosphoproteomics studies identified a total of 62 sites on nucleoporins (Phosida) (*52*), we mapped 133 phosphosites on 21 out of the 28 nucleoporins characterized in worms (**Fig. 4B**, **Data S1, Table S1**). 41% (54/133) of these sites match the minimal consensus for Cyclin-Cdk-dependent phosphorylation (pS/pT-P), among which 26% (14 out of 54) are part of Polo-docking sites suggesting that Cyclin-Cdk primes PLK-1 binding to multiple nucleoporins (**Fig. 4B**). Indeed, we identified 14 phosphorylated Polo-docking sites matching the consensus for non-self-priming (S- pS/pT-P) in the nucleoporins of the cytoplasmic filaments NPP-14^NUP214^ (5 Polo-docking sites), the central channel NPP-1^NUP54^ (1), in the transmembrane nucleoporins NPP-12^NUP210^ (1) and NPP- 22^NDC1^ (1), in the subunits of the inner ring complex NPP-8^NUP155^ (1) and NPP-19^NUP53^ (1), but also in NPP-10N^NUP98^ (2), NPP-16^NUP50^ (1), and MEL-28^ELYS^ (1) (**Data S1, Table S1**). In addition, we identified one phosphorylated Polo-docking sites matching the consensus for self-priming (S-pS/pT-X) in NPP-10N^NUP98^ (**Data S1, Table S1**).

**Fig. 4:**
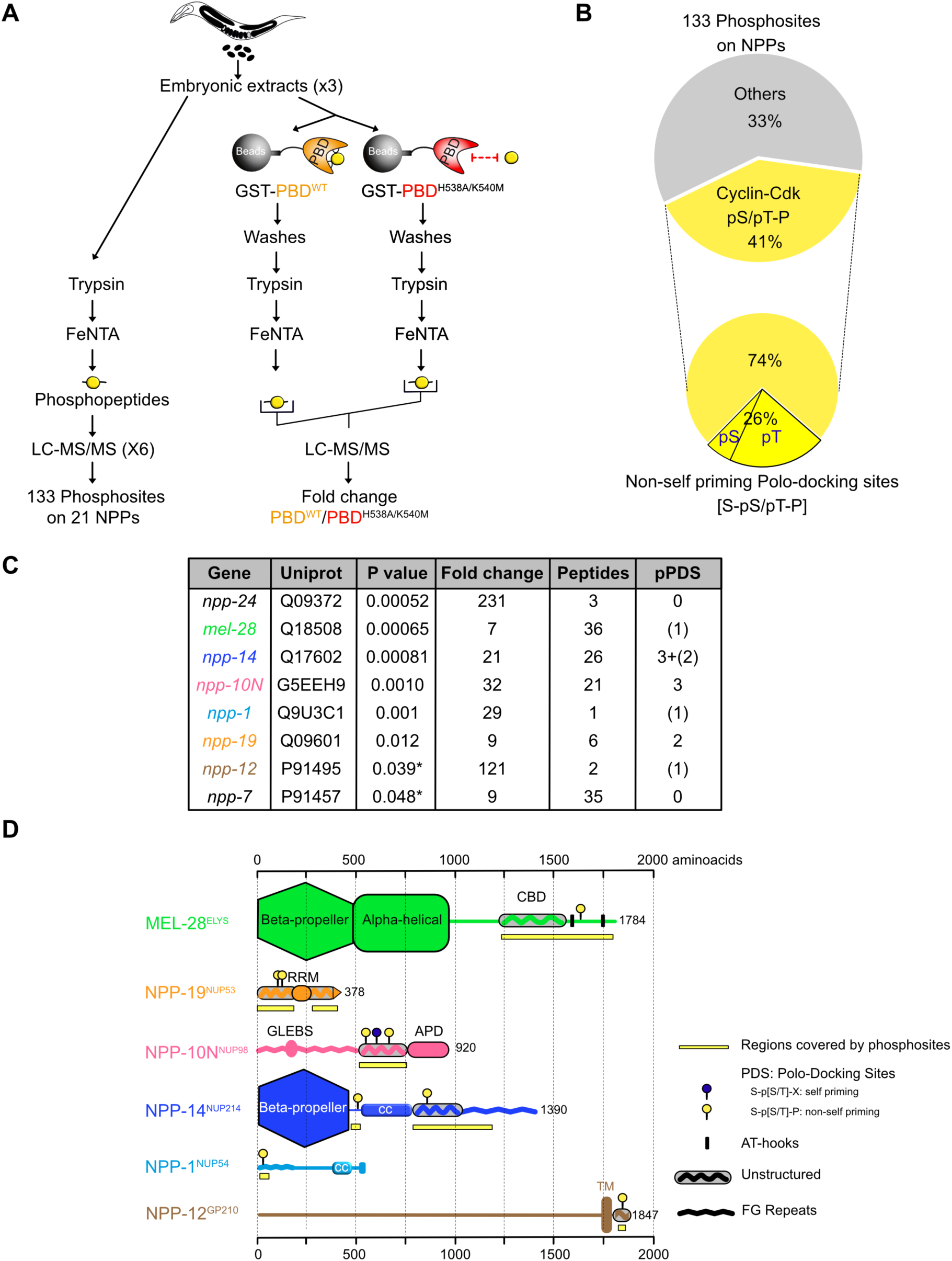
A biochemical and phosphoproteomic screen for PLK-1 targets at the nuclear pore complexes. **A-** Flow-chart of the approach used to map the nucleoporins phosphoproteome and identify nucleoporins specifically binding the Plk1 PBD, along with the phosphorylation sites responsible for the interaction. Embryonic extracts, prepared from young adults were digested with trypsin and phosphopeptides were affinity purified using FeNTA Immobilized metal ion affinity chromatography (IMAC) columns before identification by tandem mass spectrometry. A fraction of the embryonic extracts was incubated on an affinity matrix consisting of the GST-PBD wild-type or mutated on the phosphopincers. After several washing steps, retained proteins were digested with trypsin and phosphopeptides were affinity purified on FeNTA before identification by liquid chromatography and tandem mass spectrometry (LC-MS/MS). **B-** Circle chart showing the proportion of phosphorylation sites identified on nucleoporins presenting the consensus for phosphorylation by proline-directed kinases ([pS/pT-P], Cyclin-Cdk). The circle chart below presents the percentage of the phosphosites targeted by proline-directed kinases that are also part of Polo-docking sites matching the consensus for non-self priming phosphorylation [S- pS/pT-P]. **C-** Table summarizing the label-free quantitative mass spectrometry analysis of the GST-PBD wild- type or mutated on the phosphopincers pull-downs from three independent *C. elegans* embryonic extracts. The P-value (<0.05) indicates statistical significance for the enrichment of the nucleoporins in the GST-PBD wild-type pull-down. pPDS (phosphorylated Polo-docking sites) indicate the number of Polo-docking sites identified in the different nucleoporins. The value in parentheses correspond to cases where a phosphorylated peptide contains a Polo-docking site but the exact position of the phosphorylation is ambiguous. **D-** Domain organization and distribution of the phosphorylation sites on the nucleoporins specifically retained on the GST-PLK-1 PBD affinity matrix: MEL-28^ELYS^, NPP-19^NUP53^, NPP-10N^NUP98^ and NPP-14^NUP214^, NPP-1^NUP54^ and NPP-12^gp210^. CBD: Chromatin-binding domain, RRM: RNA Recognition Motif, APD: Auto-Proteolytic Domain, CC: coiled-coiled.

To identify *bona-fide* Plk1 PBD-interacting phospho-nucleoporins, we combined affinity purification with phosphoproteomics (**Fig. 4A**). We loaded embryonic extracts on an affinity matrix containing either the GST-Plk1 PBD wild-type or a control GST-Plk1 PBD^H538A/K540M^, carrying mutations on the “phosphopincers” (*47–49*), and thus defective in binding phosphopeptides (**Fig. 4A**). The retained proteins were digested with trypsin, and phosphopeptides purified on FeNTA affinity chromatography (FeNTA) before identification by LC-MS/MS. Here we specifically concentrated our analysis on nucleoporins; full details of this analysis will be published elsewhere.

Through this approach, we recovered phosphopeptides corresponding to NPP-1^NUP54^, MEL-28^ELYS^, NPP-10N^NUP98^, NPP-14^NUP214^, NPP-19^NUP53^, NPP-7^NUP153^, NPP-12^NUP210^ and NPP-24^NUP88^ (p value<0.05, n=3 independent experiments using different embryo extracts, **Table S2**) but we unambiguously detected phosphorylated Polo-docking sites only in NPP-10N^NUP98^ (3 sites), NPP- 14^NUP214^ (2 sites), and NPP-19^NUP53^ (2 sites) (**Fig. 4C, data S2, table S3**). Several MEL-28^ELYS^ phosphopeptides were recovered, including one encompassing a Polo-docking site, but the exact position of the phosphorylation could not be unambiguously assigned on this peptide. We note that this specific Polo-docking site was already detected phosphorylated in embryo extracts (**Table S1, data S1**). Likewise, we recovered in the PBD pull-downs phosphopeptides covering one Polo-docking site in NPP-1^NUP54^ and NPP-12^NUP210^. As for MEL-28^ELYS^, both Polo-docking sites were unambiguously phosphorylated in the embryonic extracts strongly suggesting that MEL-28^ELYS^, NPP-1^NUP54^ and NPP-12^NUP210^ were specifically recovered in the PLK-1 PBD pull-downs via these phosphorylated Polo-docking sites.

Next, we analyzed the distribution of the phosphorylation sites on the various functional domains of these nucleoporins. MEL-28^ELYS^, the first nucleoporin to associate with chromatin at the end of mitosis (*53, 54*), is a large protein composed of multiple functional domains: a N-terminal β-propeller domain, a central α-helical domain, and an unstructured C-terminal region that includes two AT hooks (**Fig. 4D**). The N-terminal domains are required for NPC targeting and kinetochore localization, whereas the C-terminal domain is required for chromatin binding (*55*). Most of the phosphorylation sites identified on MEL-28^ELYS^, including the phosphorylated Polo-docking site, localized on the chromatin-binding region, suggesting that phosphorylation could modulate this activity (**Fig. 4D**).

Phosphorylation sites were also restricted to a defined region in NPP-10N^NUP98^, which contains GLFG domains and a Rae1 binding site in the N-terminal region, similar to its human counterpart NUP98 (**Fig. 4D**). The C-terminal part, located upstream of the conserved autoproteolytic domain is however poorly conserved (**Fig. S4A**) (*36*). This region is predicted to be unstructured in both human NUP98 and *C. elegans* NPP-10N^NUP98^. Most of the phosphorylation sites localized to this intrinsically disordered region of NPP-10N^NUP98^, embedded in motifs matching the consensus for PLK-1 phosphorylation or PLK-1 docking (**Fig. 4D, Fig. S4A**).

Likewise, NPP-14^NUP214^, besides containing an evolutionarily conserved β-propeller domain, is mainly unstructured with FG repeats in the C-terminal part of the protein. Here again most of the phosphosites localized to the non-conserved and unstructured region (**Fig. 4B**). Phosphorylated Polo- docking sites were also identified in the unstructured regions of NPP-19^NUP53^, which contains an RNA Recognition Motif (RRM) dimerization domain and a C-terminal alpha-helix required for membrane binding (*56*). Finally, the unique phosphorylated Polo-docking sites (SpTP) found in NPP-1^NUP54^ and NPP-12^gp210^ also localized in the unstructured N-terminal FG repeat of NPP-1^NUP54^ and C-terminal cytoplasmic tail of NPP-12^gp210^ **(Figure 4D)**.

In summary, most of the identified phosphorylation sites are distributed in unstructured, intrinsically disordered regions of the nucleoporins, specifically interacting with the PLK-1 PBD. Most importantly, the identification of NPP-1^NUP54^, NPP-14^NUP214^, NPP-19^NUP53^ and NPP-10N^NUP98^ as specific phospho-binding partners of the Plk1 PBD is consistent with our findings showing that PLK- 1 depletion affects the disassembly of these nucleoporins, and their direct binding partners, during mitosis (**Fig. 2**).

### NPP-10N^NUP98^ contributes to PLK-1 recruitment to the nuclear envelope, possibly through direct phospho-dependent binding to the PBD

We have previously reported that nucleoporins of the central channel recruit PLK-1 to the NE (*7*). We thus investigated whether NPP-14^NUP214^, NPP-10N^NUP98^, and NPP-19^NUP53^, identified as specific phosphorylated binding partners of the PBD, also contribute to the recruitment of PLK-1 to the nuclear envelope in prophase. To this end, we used spinning disk confocal microscopy to monitor the recruitment of endogenously tagged PLK-1::sGFP to the nuclear envelope. RNAi-mediated depletion of NPP-14^NUP214^ did not affect PLK-1::sGFP recruitment to the nuclear envelope **(Fig. 5A)**. However, partial depletion of NPP-10N^NUP98^ or NPP-19^NUP53^ drastically reduced PLK-1::sGFP signal at the nuclear envelope (**Fig. 5A**). Knocking down the central channel nucleoporin NPP-1^NUP54^ resulted in a similar phenotype (**Fig. 5A**), consistent with our previous observations (*7*).

**Fig. 5:**
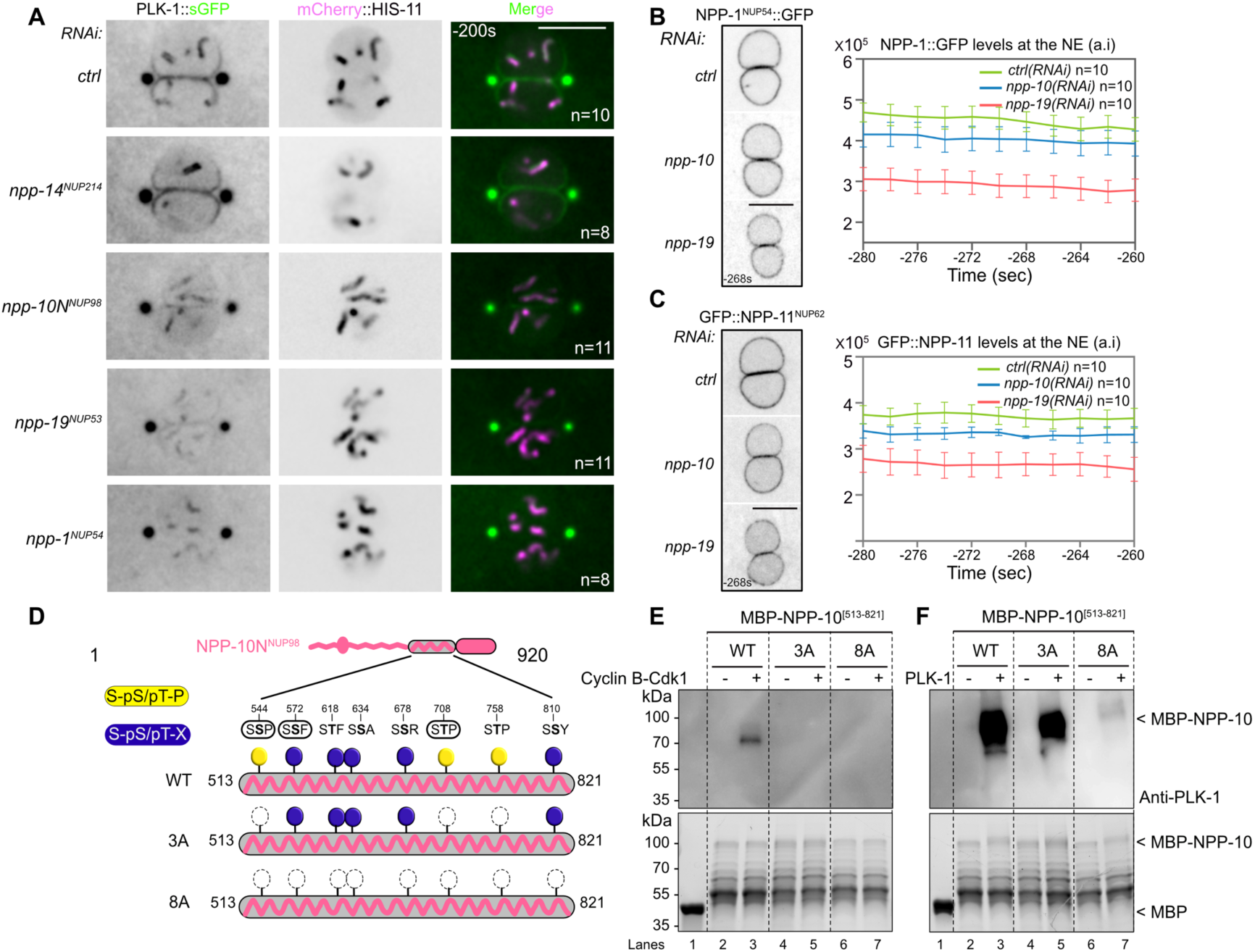
Phosphorylated NPP-10N^NUP98^ binds the PLK-1 PBD and contributes to PLK-1 recruitment to the nuclear envelope. **A-** Spinning disk confocal micrographs of embryo expressing PLK-1::sGFP (shown alone, and in green in the merged images) and mCherry::HIS-11 (magenta, in the merged image) exposed to control *(ctrl)*, *npp-14^NUP214^*, *npp-10N^NUP98^*, *npp-19^NUP53^* or *npp-1^NUP54^* RNAi. At this stage the juxtaposed male and female pronuclei are in prophase. All panels are at the same magnification. Scale Bar, 10μm. **B- C-** Spinning disk confocal micrographs of embryos expressing GFP::NPP-1^NUP54^ (**B**) or GFP::NPP-11^NUP62^ (**C**) exposed to control *(ctrl)*, *npp-10N ^NUP98^*, or *npp-19^NUP53^* RNAi. The graphs on the right present the quantification of GFP::NPP-1^NUP54^ or GFP::NPP-11^NUP62^ average intensity at the nuclear envelope at six time-points relative to anaphase onset. (Time is in sec relative to anaphase onset). **D-** Schematic and domain organization of NPP-10N^NUP98^. The C-terminal disordered NPP-10N^NUP98^ [513-821] fragment contains three polo-docking sites matching the consensus for non-self-priming (yellow, phosphorylatable by Cyclin-Cdk1), and five sites matching the consensus for self-priming (blue, phosphorylatable by PLK-1). The sites that are circled have been identified *in vivo*. Besides the WT, two additional fragments containing the three non-self-priming polo-docking sites (3A), or all the polo-docking sites substituted with Alanine are presented. **E-F** *In vitro* kinase assays were performed with CyclinB-Cdk1 or PLK-1 kinases and the NPP- 10N^NUP98^ [513-821] fragments WT, 3A or 8A tagged with the maltose-binding protein (MBP) as substrates. The samples were subjected to SDS-PAGE, followed by a Far-Western ligand-binding assay using the Polo-box domain fused to GST (upper panel). The bottom panel shows the Stain-Free Blot (Chemidoc, Bio-Rad) of the same membrane.

Central channel nucleoporins are anchored to the NPC via interaction with NPP-13^NUP93^, which is itself connected through NPP-19^NUP53^ binding (**Fig. 1A**). Thus, inhibition of *npp-19^NUP53^* might be expected to cause a reduction of the central channel nucleoporins at the nuclear envelope, and consequently of PLK-1::sGFP. Accordingly, *npp-19^NUP53^* inhibition did reduce GFP::NPP-1^NUP54^ and GFP::NPP-11^NUP62^ signals at the nuclear envelope (**Fig. 5B and 5C**). However, *npp-10N^NUP98^* depletion only mildly reduced central channel nucleoporin levels at the nuclear envelope (**Fig. 5B and 5C**), while profoundly reducing PLK-1::sGFP levels (**Fig. 5A**). This result suggested that NPP- 10N^NUP98^ may directly contribute to PLK-1::sGFP recruitment, presumably by interacting with the PLK-1 PBD, which would be consistent with the NPP-10N^NUP98^ phosphorylated consensus Polo- docking sites we identified in the PBD pull-downs (**Fig. 4**). We tested this hypothesis using a Far- Western ligand-binding assay. We pre-phosphorylated the NPP-10N^NUP98^ C-terminal fragment [513- 821] using CyclinB-Cdk1 or PLK-1 kinases, and then tested its ability to interact with the PLK-1 PBD. NPP-10N^NUP98^ [513-821] pre-phosphorylated by either kinase readily interacted with the PLK- 1-PBD (**Fig. 5E and 5F lane 3**) but not with a version of the PBD carrying mutated phosphopincers (**Fig. S4B and S4C lane 3**), demonstrating that this interaction is phospho-dependent. To test if the Polo-docking sites accounted for PBD binding, we then analyzed NPP-10N^NUP98^ [513-821] fragments containing various alanine substitutions in the predicted Polo-docking sites.

The NPP-10N^NUP98^ fragment harbors three putative Polo-docking sites matching non-self-priming and binding (1 SSP and 2 STP sites) (**Fig. 5D, S4A**), and we recovered phosphopeptides specifically phosphorylated on these three sites in the PLK-1 PBD pull-downs (**Fig. 4, table S3**). In the far- western binding assay, Alanine substitutions of these sites abrogated NPP-10N^NUP98^ binding to the PLK-1 PBD, primed by CyclinB-Cdk1 (**Fig. 5E, lane 5**). However, these Alanine substitutions did not disrupt binding when NPP-10N^NUP98^ was primed by PLK-1 (**Fig. 5F, lane 5**), indicating that PLK- 1 phosphorylates additional Polo-docking sites. Consistently, sequence analysis identified at least five other putative Polo-docking sites matching the self-priming and binding consensus (SSX/STX). Four of these five sites perfectly match PLK-1 phosphorylation sites (see Eukaryotic Linear Motif resource for Functional Sites in Proteins http://elm.eu.org/) (**Fig. S4A**), among which one was recovered phosphorylated in the PBD pull-downs (**Fig. 4**). Alanine substitutions of the eight Polo-docking sites fully abolished the binding of NPP-10N^NUP98^ to the PLK-1 PBD primed by PLK-1 (**Fig. 5F lane 7**).

We conclude that CyclinB-Cdk1 and PLK-1 both phosphorylate NPP-10N^NUP98^ on multiple Polo-docking sites in its unstructured region, thereby promoting its binding to the PLK-1 PBD. This interaction contributes to the recruitment of PLK-1 to the NPC, where it docks on NPP-10N^NUP98^. PLK-1 docking might be required for subsequent phosphorylations or it might disrupt NPP-10N^NUP98^ binding to its partners at the NPC, both of which would promote NPP-10N^NUP98^ dissociation from the NPC.

### Embryos expressing NPP-19^NUP53^ 10A defective in PLK-1 binding are deficient in inner ring complex disassembly

We then investigated the functional importance of the PLK-1 - NPP-19^NUP53^ interaction for the disassembly of the inner ring complex (**Figs. 1E, 2D, 3**). To this end, we searched for all the NPP- 19^NUP53^ Polo-docking sites responsible for interacting with the PLK-1 PBD to engineer an NPP- 19^NUP53^ variant specifically defective in PLK-1 binding and evaluate its impact on NPC disassembly *in vivo*.

NPP-19^NUP53^ contains ten potential Polo-docking sites relatively well conserved in nematode species (**Fig. S5**); seven sites match the consensus for self-priming (S-S/T-X), whereas three sites match the consensus for non-self-priming (S-S/T-P) **(Fig. 6A)**, one of which was identified in the PLK-1 PBD pull-downs (**Fig. 4**, table 1). Alanine substitution of the three non-self-priming sites (S-A-P) abolished NPP-19^NUP53^ binding to the Plk1 PBD primed by Cyclin B-Cdk1 in a Far-Western binding assay (**Fig. 6B left, lane 4**). However, these three alanine substitutions did not significantly affect the interaction when PLK-1 was the priming kinase (**Fig. 6B right, lane 4**). Alanine substitutions of all ten Polo-docking sites were required to eliminate the ability of PLK-1 to trigger the interaction between NPP- 19^10A^ and the Plk1 PBD (**Fig. 6B, lane 6**). We obtained similar results using the shorter and more soluble NPP-19^NUP53^ [1-301] fragment tagged with 6xHis (**Fig. S6C, S6D**). These studies indicate that Cyclin B-Cdk1 and PLK-1 both prime NPP-19^NUP53^ binding to the Plk1 PBD via the phosphorylation of multiple polo-docking sites, similar to NPP-10N^NUP98^.

**Fig. 6:**
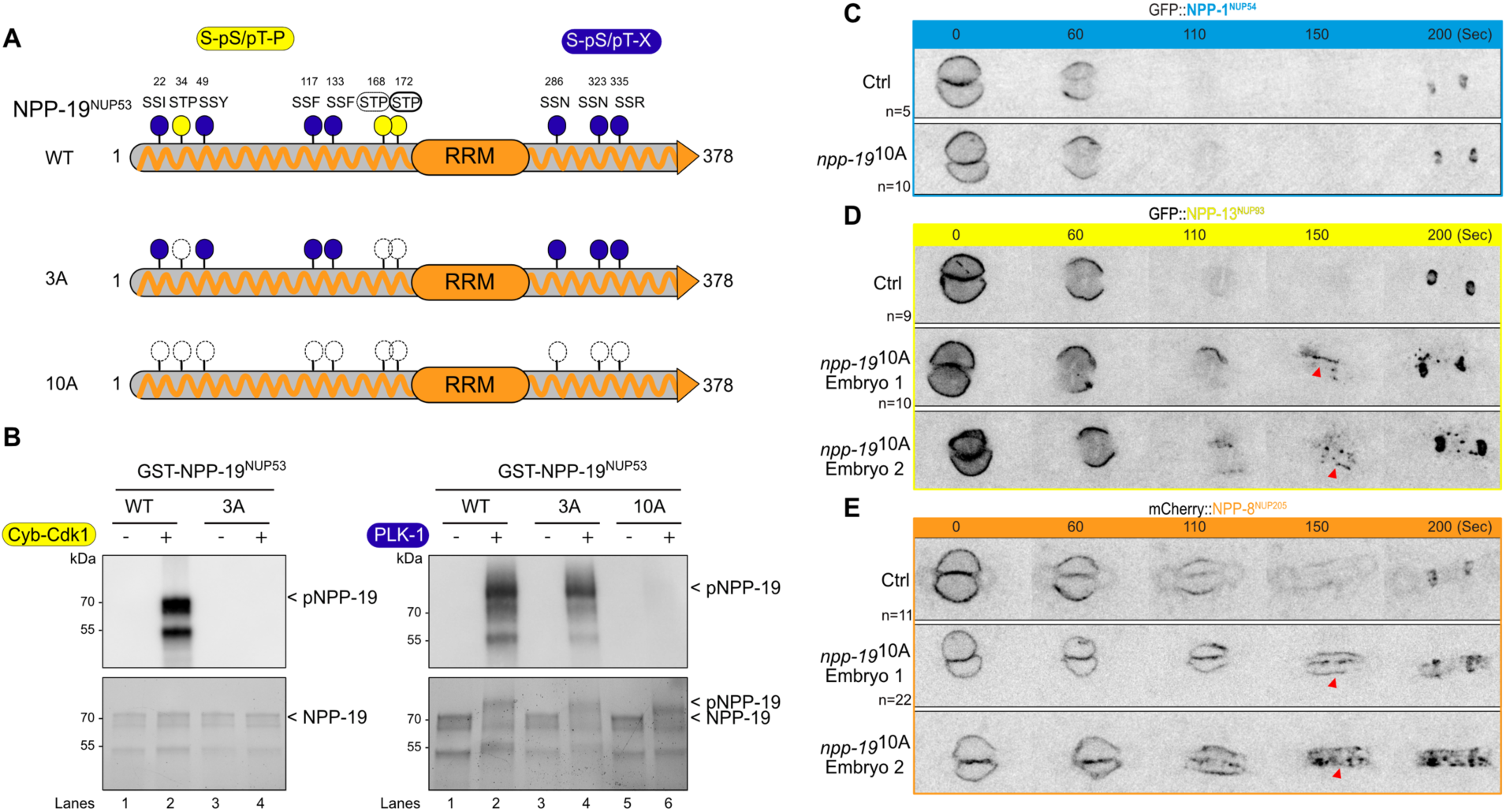
Multisite phosphorylation of NPP-19^NUP53^ is required for the disassembly of the inner ring complex. **A-** NPP-19^NUP53^ contains three polo-docking sites matching the consensus for non-self-priming (yellow), and seven sites matching the consensus for self-priming (blue). Besides the WT, two additional NPP-19^NUP53^ variants containing the three non-self-priming polo-docking sites (3A), or all the polo-docking sites (10A) substituted with Alanine are presented. **B-** *In vitro* kinase assays were performed with CyclinB-Cdk1 or PLK-1 kinases and the NPP-19^NUP53^ WT, 3A or 10A tagged with the Glutathion-S-transferase (GST) as substrates. The samples were subjected to SDS-PAGE, followed by a Far-Western ligand-binding assay using the Polo-box domain fused to GST (upper panel). The bottom panel shows the Stain-Free Blot (Chemidoc, Bio-Rad) of the same membrane. **C-D-E** Spinning disk confocal micrographs of control (upper panels) or *npp-19^NUP53^* 10A mutant embryos (lower panels) expressing GFP::NPP-1^NUP54^ (**C**), GFP::NPP-13^NUP93^ (**D**) or GFP::NPP- 8^NUP155^ (**E**) during mitosis. Timings in second (sec) are relative to pronuclei juxtaposition (0). All panels are at the same magnification. Scale Bar, 10μm.

Next, we used CRISPR/Cas9-based mutagenesis to engineer an *npp-19^NUP53^* allele carrying alanine substitutions in the ten Polo-docking sites to investigate the functional consequences of a defect in PLK-1 binding to NPP-19^NUP53^ for NPC disassembly *in vivo*. These ten alanine substitutions did not alter NPP-19^NUP53^ stability nor its ability to localize to the NPCs, as revealed by the normal levels of expression and the proper localization of NPP-19^NUP53^ 10A to the nuclear envelope (**Fig. S7A, B**). Furthermore, *npp-19*^NUP53^ 10A mutant embryos were viable and did not present the major cell cycle defects previously observed in *npp-19^NUP53^* loss-of-function mutant embryos (*31*) (**Fig. S7C**). PLK- 1::sGFP localized to the nuclear envelope in prophase in these *npp-19^NUP53^* 10A mutant embryos (**Fig. S7D**), indicating that the defect in PLK-1::sGFP recruitment to the NE previously observed in *npp- 19(RNAi)* embryos (**Fig. 5A**) is due to the loss of NPP-19-associated PLK-1 recruiters (see Discussion). The dynamics of the central channel nucleoporin GFP::NPP-1^NUP54^ was also largely unaffected in *npp-19^NUP53^* 10A mutant embryos during mitosis (**Fig. 6C**). However, disassembly of the inner ring complex nucleoporins GFP::NPP-13^NUP93^ and mCherry::NPP-8^NUP205^, which are direct binding partners of NPP-19^NUP53^, was compromised. GFP::NPP-13^NUP93^ persisted at the nuclear envelope during mitosis in *npp-19^NUP53^* 10A mutant embryos (**Fig. 6D,** red arrowheads). Likewise, mCherry::NPP-8^NUP205^ remained associated with the nuclear envelope throughout mitosis in *npp-19^NUP53^* 10A mutant embryos and failed to relocalize to the centriculum, as observed in control embryos (**Fig. 6E**).

These observations indicate that phospho-dependent recruitment of PLK-1 to NPP-19^NUP53^, at the core of the nuclear pore complexes, is required for efficient inner ring complex disassembly.

## Discussion

The early *C. elegans* embryo provides a rich developmental context for dissecting the mechanisms regulating nuclear envelope breakdown (*8, 9*). In the zygote, spatio-temporal coordination of nuclear envelope breakdown is required for the unification of the two parental genomes into a single nucleus after the first mitosis, and PLK-1 is critical to this process (*15*). We previously reported that PLK-1 is dynamically recruited to the nuclear envelope in prophase to trigger lamina depolymerization (*7, 10, 11*) but its exact contribution for NPC disassembly was not clear.

Using a combination of live-imaging, biochemistry and phosphoproteomics, we show unequivocally that PLK-1 promotes NPC disassembly independently of lamina depolymerization. We characterized NPC disassembly in the early embryo and identified the critical PLK-1 targets in this process. Our findings (summarized in **Fig. 7**) indicate that PLK-1 is dynamically recruited, in a phospho-dependent manner, to intrinsically disordered regions of multivalent linker nucleoporins to trigger the disassembly of NPC subcomplexes, including the cytoplasmic filaments, the inner ring complex and the central channel. We also reveal a distinct order of NPC disassembly in *C. elegans* embryos compared to that of human cells (**Fig. 7**).

**Fig. 7:**
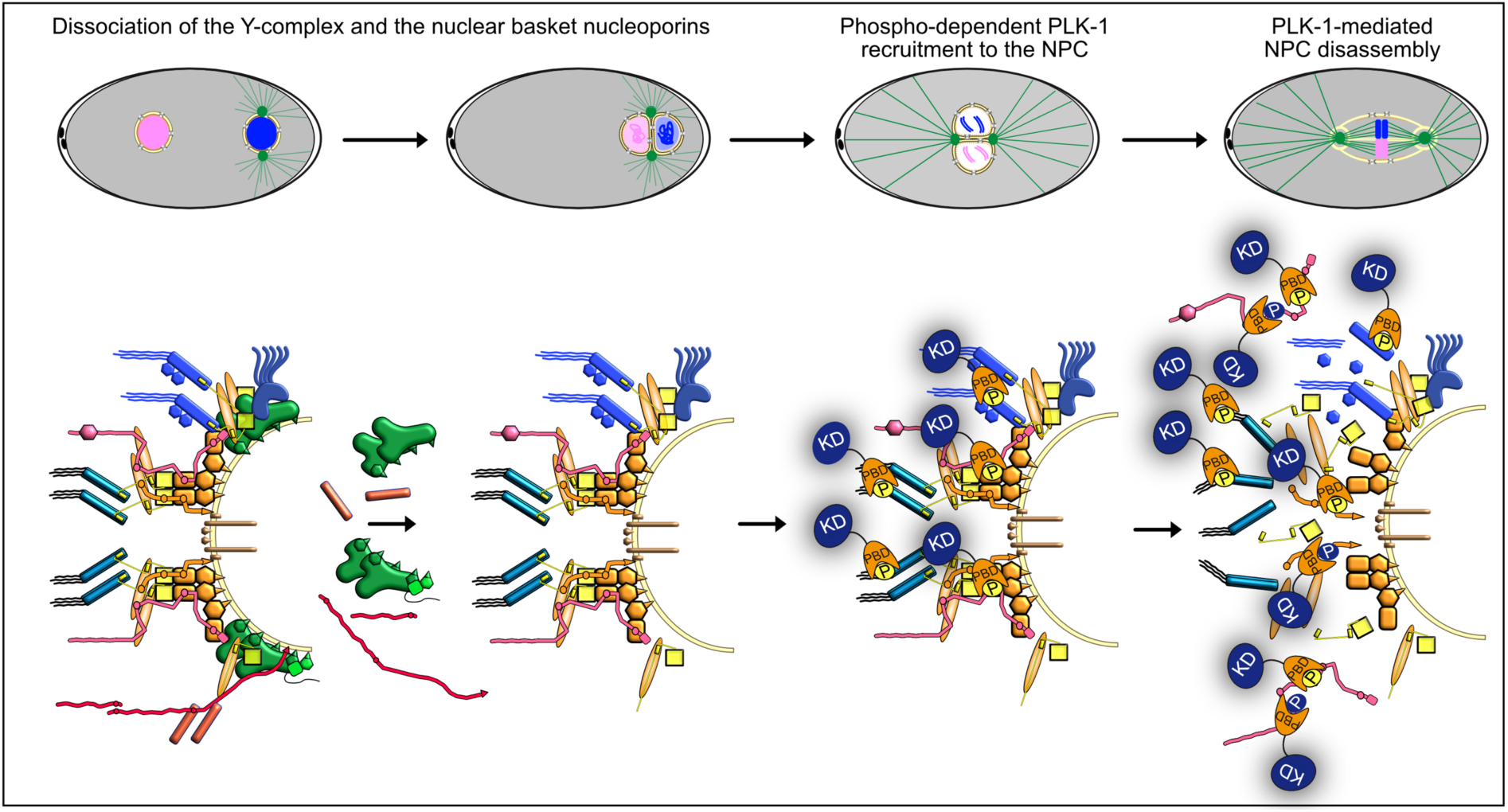
PLK-1 is recruited to multiple multivalent nucleoporins to dismantle the nuclear pore complexes. Dynamics of nuclear pore complexes during the first mitotic division of the *C. elegans* embryo. During pronuclear migration and meeting, the Y-complex and nuclear basket nucleoporins leave the NPC. PLK-1 is then recruited to the NPC, 250s before anaphase onset, via its PBD domain in a phospho-dependent manner just before NEBD when the two pronuclei are juxtaposed. PLK-1 is recruited via the multivalent nucleoporins NPP-10N^NUP98^, NPP-19^NUP53^, the central channel nucleoporins and by NPP-14^NUP214^ phosphorylated at Polo-docking sites by CyclinB-Cdk1 and PLK- 1 itself. PLK-1 recruitment and docking on nucleoporins promotes NPC disassembly.

### Characterization of NPC disassembly in early *C. elegans* embryo

NPCs are massive structures assembled from various nucleoporin subcomplexes that dynamically assemble and disassemble during the cell cycle (*25*). While the mechanisms driving NPC assembly are beginning to emerge (*57, 58*), how NPCs are disassembled in various organisms is not well understood. In particular, the order of component disassembly is still elusive. This question has been mainly addressed using cultured human cells (*18*). In this context, the nucleoporin NUP98, the gatekeeper of the nuclear pore, is the first nucleoporin to leave the NPC (*17, 46, 59*) followed by the rapid and nearly synchronous disassembly of the other nucleoporins (*46*). NUP98 maintains the permeability barrier of the NPC by virtue of the self-cohesive interactions of its N-terminal GLFG domain (*26*) and thus its departure from the NPCs facilitates nuclear permeabilization at the onset of mitosis (*17, 46*).

In the early *C. elegans* embryo, the order of nucleoporin disassembly had yet to be investigated. We found here that the nuclear basket nucleoporins NPP-7^NUP153^, NPP21^TPR^, and the Y complex subunits MEL-28^ELYS^, NPP-5^NUP107^, NPP-6^NUP160^, NPP-18^SEH-1^ all leave the NPC early in prophase while the two pronuclei are still migrating or are about to meet at the posterior pole of the embryo, well before their nuclear envelope becomes permeable. It is only later, when NPP-10N^NUP98^, the central channel and the inner ring complex nucleoporins leave the NPC that the permeability barrier is breached. Thus, the order of NPC disassembly in the *C. elegans* embryo appears to be different from that of human cells, with some nucleoporin subcomplexes leaving the NPC earlier without affecting the permeability barrier. This echoes one study in human cells in which Auxin-mediated depletion of the entire Y-complex had little impact on the stability of the inner ring complex and the central channel nucleoporins (*60*). The functional relevance of the earlier departure of the nuclear basket and the Y complex subunits from the NPC in *C. elegans* as compared to human cells is presently unclear. A portion of the Y complex relocalizes to the kinetochore during mitosis in human cells (*61*) and in *C. elegans* (*62*) where it contributes to kinetochore assembly and chromosome segregation (*62, 63*). Perhaps the holocentric nature of kinetochores of *C. elegans* requires an earlier recruitment of the Y subunits for their proper assembly.

### PLK-1 is independently recruited to multiple nucleoporins

While PLK-1 is not required for the removal of the nuclear basket and the Y complex subunits from the NPC, it is critically required for the disassembly of the central channel, the cytoplasmic filaments and the inner ring complex. After activation in the cytoplasm via phosphorylation of its activation segment (*64, 65*), PLK-1 is recruited to the nuclear envelope in prophase (*7, 16*), before they are permeable. Our results suggest that after its delivery to the NPC, PLK-1 uses its Polo-box domain to dock on the intrinsically disordered regions of multiple nucleoporins, phosphorylated on Polo- docking sites by Cyclin-Cdk1 and PLK-1 itself. While in human cells several mitotic kinases are involved in this process (*16–18*), PLK-1 has a prominent role in NPC disassembly in *C. elegans*. PLK-1 not only docks on NPP-19^NUP53^ as in human cells (*16*) but also on NPP-10N^NUP98^, NPP- 14^NUP214^, and on the central channel nucleoporins phosphorylated on multiple Polo-docking sites. Accordingly, alanine substitution of the ten Polo-docking sites in NPP-19^NUP53^, which abrogates its binding to the PLK-1 PBD *in vitro,* is not sufficient to prevent PLK-1 recruitment to the NPC, clearly indicating the presence of additional PLK-1 recruiters at the NPC in *C. elegans*. NPP-1/4/11 and NPP-10N^NUP98,^ which line the central channel, are the most accessible nucleoporins and thus likely act as primary recruiters of PLK-1 at the NPC.

These nucleoporins all possess phosphorylated Polo-docking sites matching both non-self-and self- priming and binding motifs, suggesting that CyclinB-Cdk1 might dictate the initial binding of the first PLK-1 molecules before PLK-1 itself promotes its own docking and further accumulation required for NPC disassembly.

### How does PLK-1 trigger NPC disassembly?

*In vitro* binding experiments and structural studies have shown that NUP98 and NUP53 interact with several nucleoporins and thereby serve as multivalent linkers within the NPC scaffold (*22, 32–36, 66*). NUP98 binds through its C-terminal NPC targeting domain subunits of the Y-complex and members of the inner ring complex notably NUP205 and NUP155 (*35, 36*). Likewise, NUP53 uses multiple regions to interact with distinct subunits of the inner ring complex including NUP93, NUP155 and NUP205 but also with the transmembrane protein NDC1 (*22, 32–36, 56, 66, 67*). Previous studies in human cells showed that NUP98 and NUP53 are targeted by multiple kinases including Cyclin-Cdk1, NIMA and Plk1 (*16, 17*). Cyclin-Cdk1 phosphorylates NUP98 on 8 sites and NUP53 on 16 sites (*16, 17*) (**Fig. S4 and S5**). Notably, introduction in human NUP98 of two phosphomimetic residues is sufficient to abrogate its binding to NUP155 C-terminal domain (*17, 35*). Structural analysis of the NUP98-NUP155 interface using *C. thermophilium* proteins has revealed a possible mechanism that could explain how the phosphorylation of these two residues affects this interaction. NUP98 phosphorylation induces structural rearrangements destroying the hydrophobic interface required for NUP155 binding (*17, 35*). This type of mechanism is ideally suited for the unzipping of NUP-NUP interactions because the phosphorylatable residues are directly accessible to the mitotic kinases. Another (non-exclusive) possibility is that the docking of PLK-1 on nucleoporins competes with and displaces interaction between these nucleoporins and their partners. In support of this hypothesis, we have noticed that several Polo-docking sites in NPP-19^NUP53^ localize within or near conserved regions of interaction among NUP53 and its binding partners (**Fig. S5**). For instance, NUP53 binds NUP93 via a linear ten-residue motif [SGAPPVRSIY], which recognizes a hydrophobic surface on the NUP93 solenoid (*36*). This motif, which is evolutionarily conserved between NUP53 orthologues, is immediately flanked by Polo-docking sites, both in *C. elegans* and in human NUP53 (**Fig. S5**). Furthermore, PLK-1 can dimerize on substrates containing several Polo- docking sites (*68, 69*), which would increase the steric hindrance at the NPC. PLK-1 docking on nucleoporins might also destabilize allosterically NPC subcomplexes, as reported in the case of Kap121 binding to Nup53 (*70*). We thus speculate that PLK-1 docking on NPP-19^NUP53^ by itself might contribute to displace NPP-19^NUP53^-NPP-13^NUP93^ interactions. Similarly, PLK-1 docking might displace the interactions between NPP-19^NUP53^ and NPP-8^NUP155^ or NPP-3^NUP205^ because several Polo-docking sites are also present in the regions of interactions (**Fig. S5**). Finally, PLK-1 docking on NPP-10N^NUP98^ could similarly disrupts its interaction with partners (**Fig. S4**).

Beyond NPP-19^NUP53^ and NPP-10N^NUP98^, PLK-1 appears to have multiple independent targets at the NPC in *C. elegans* including NPP-14^NUP214^ and the central channel nucleoporins. This is also supported by our finding that worms expressing the NPP-19^NUP53^ 10A variant specifically defective in PLK-1 docking displays problems in inner ring disassembly but not in central channel nucleoporin disassembly.

Comprehensive identification of all the sites phosphorylated by PLK-1 on nucleoporins and the functional dissection of their roles in NPC disassembly will be a major future challenge. Obtaining or modelling the structure of the *C. elegans* NPC would be helpful in this endeavor.

Overall, despite some noticeable differences between the mechanisms of NPC disassembly in human cells versus *C. elegans*, NPP-10N^NUP98^ and NPP-19^NUP53^ are major phosphorylation targets in both species. Remarkably, the phosphorylated sites on NPP-10N^NUP98^ and NPP-19^NUP53^ are poorly conserved between human and *C. elegans* proteins, and targeted by different kinases, yet they are concentrated in the exact same regions. Phosphorylation of intrinsically disordered regions of multivalent linker nucleoporins thus appears to be evolutionarily conserved mechanism driving nuclear pore complex disassembly during mitosis.

### Limitations of the study

Monitoring the live dynamics of endogenous nucleoporins requires their tagging with large fluorescent markers, NeonGreen, mCherry or GFP. While all the lines used in this study expressing endogenously tagged nucleoporins are viable, some of them are slightly compromised in function. In particular, lines expressing mCherry::NPP-8^NUP155^ or NeonGreen::NPP-10N^NUP98^. For instance NeonGreen::NPP-10N^NUP98^ was synthetic lethal with *npp-14^NUP214^* null (L. Thomas and G. Seydoux). We also noticed some cumulative defects when trying to generate lines co-expressing two tagged nucleoporins, in particular combining GFP::NPP-13^NUP93^ and mCherry::NPP-8^NUP205^ that belong to the same complex. This precluded the simultaneous visualization of multiple tagged nucleoporins in the same embryo. Although we have not tested all combinations, care should be taken when analyzing the dynamics of nucleoporins in strains co-expressing multiple tagged nucleoporins.

Unfortunately, we could not look at NPP-19 10A during mitotic progression, we do not have an endogenously tagged fluorescent version, and it is technically challenging to visualize persisting NPP-19 foci during mitosis by immunostaining using our antibody.

Finally, it is worth mentioning that the *C. elegans* phosphoproteome has been generated from total embryonic extracts that are not synchronized and thus the reported phosphosites on nucleoporins might be present in interphase or mitosis.

## Materials and Methods

### Nematode strains and RNAi

*C. elegans* strains were cultured and maintained using standard procedures (*71*). RNAi was performed by the feeding method using HT115 bacteria essentially as described (*72*) except that 2 mM of IPTG was added to the NGM plates and in the bacterial culture just prior seeding the bacteria. As control, animals were exposed to HT115 bacteria harboring the empty feeding vector L4440 (mock RNAi). RNAi clones were obtained from the Arhinger library (Open Source BioScience). For partial *npp-1* and *npp-14* inactivation, L4 animals were fed with bacteria for 24 h at 20°C. For partial *npp-19* and *npp-10* inactivation, L4 animals were fed with bacteria for 7-9 h at 20°C. For partial *plk- 1* inactivation, L4 animals were fed with bacteria for 7-9 h at 23°C.

### CRISPR/Cas9 genome engineering

To endogenously tag *npp-21*, a double-stranded PCR product encoding GFP and flanked by 33-35nt sequences with homology to *npp-21* was synthesized in two steps using first primers B767 and B783 followed by primers B769 and B784. Plasmid pBN221 *npp-21* sgRNA was generated by whole-plasmid PCR amplification using pBN180 (*55*) as template and primers B635 and B782. Strain BN424 *dpy-10(cn64) npp-21::gfp* was generated by microinjection of the GFP PCR fragment together with plasmids pBN221 *npp-21* sgRNA, pBN207 *dpy-10* sgRNA (*55*), #1286 *eft-3p::Cas9* (*73*) and *dpy-10* single-stranded oligodeoxynucleotide (ssODN) B725, into the gonads of N2 young adult hermaphrodites (*74*). Next, *npp-21::gfp* was segregated from *dpy-10(cn64)* by successive crosses with BN189 and N2 to generate BN1062 *npp-21::gfp lmn-1p::mCh::his-58*.

Targeting plasmids pBN317 *g>f>p::npp-19* and pBN394 *g>f>p::npp-24* (">" denotes an FRT site) for SapTrap CRISPR/Cas9 gene modification were obtained as described (*75*)using primers B969- B974 and B1085-B1090, respectively, as well as pMLS256, pMLS288 and pBN312 as donor and receptor plasmids (*75, 76*)). Plasmid pBN404 *g>f>p::npp-19* was derived from pBN317 by whole-plasmid PCR using primers B1070+B1071 to eliminate the unc-119(+) gene. Strain BN739 *npp- 24(bq15[g>f>p::npp-24])* II was generated by microinjection of targeting plasmid pBN394 into the gonads of HT1593 *unc-119(ed3)* III young adult hermaphrodites (*75*). Targeting plasmid was injected at 65 ng/µl together with plasmids #1286 *eft-3p::Cas9* (*73*), 25 ng/µl), pBN1 (10 ng/µl), pCFJ90 (2.5 ng/µl) and pCFJ104 (5 ng/µl). Successful modification of the target loci was confirmed by detection of fluorescent fusion proteins in the nuclear envelope throughout the body of wild-type moving animals without ectopic mCherry expression from the co-injection markers. The *unc-119(+)* selection marker was next excised by microinjection with Cre expression plasmid pMLS328 (*75*). Finally, uncoordinated hermaphrodites were outcrossed to N2 males to remove the *unc-119(ed3)* allele.

Endogenous tagging of *npp-19* was performed based on protocols for nested CRISPR (*77*) and “hybrid” partially single-stranded DNA donors (*78*). First, we inserted a ssODN containing gfp 5’ and 3’ sequences immediately after the *npp-19* ATG by microinjection of ssODNs B1205 (1 µM) and B1304 (4 µM), crRNAs B1206 (2.5 µM) and B1069 (10 µM), trRNA (12.5 µM) and Cas9 (3 µM) into the gonads of HT1593 *unc-119(ed3)* III young adult hermaphrodites. Wildtype-moving F1s (heterozygous for unc-119) were isolated, left to lay eggs and analyzed by PCR with primers B1305 and B1306. A sequence verified line was established (*npp-19(bq27[ssODN B1304]) II; unc-119(ed3) III*) and microinjected with ssODN B1205 (1 µM), crRNAs B1206 (2.5 µM) and B1311 (10 µM), trRNA (12.5 µM), Cas9 (3 µM) and a hybrid PCR product (∼0.1 µg/µl) generated by mixing PCR A (amplified from pBN404 with primers B1266 and B1267) and PCR B (amplified from pBN404 with primers B1268 and B1269). Successful candidates were verified by expression of GFP::NPP-19 in the nuclear envelope in all cell types and crossed with BN189 to generate BN1018 *g>f>p::npp-19 lmn-1p::mCherry::his-58*.

### Generation of *npp-19^NUP53^* 10A allele

The worm strain expressing *npp-1* 10A was generated via CRISPR/Cas9 insertion of a recoded gDNA of *npp-19* harboring the mutation of the residues S22, T34, S48, S117, S133, T168, T172, S286, S323 and S335 into alanines (SunyBiotech). The recoded part of *npp-19* gDNA consisted of the nucleotides 271 to 936 which are targeted by the RNAi against *npp-19* from the Arhinger library. Wild-type and RNAi resistant sequences are as follows.

### Wild-type sequence (nucleotides 271-936)

TAGCCCGCTCAACACTGCATCGGCACCATGTTCGGATATTTTCGCTGTTTCTGCACCGGCAGTGCCGCAGCATTTGAAGGATACACCGGGCTCTAAATCAGTTCATTGGTCTCCATCTTTGGTGCAATCTGGTGAAAAGTCGGCGGCACAAACACAGAATACACCTGCCAACTTGTCTTTCGGAGGAAATTCATCATTTTCGGCGCCGACAAAGCCAGCTCCTCAGTCGATTCAGACATCATCTTTCGGTGGTCAAGCGATGCATGgttagaatttacattttttcgcgaatatttaaaataaaattgattttcagCACCACCTCTTCGATCTCTTCGCGACAAAGTTGAACCAGCGAAAAAGATCTCCAGACGAAATACATTCACTGCCAGATCAACACCACTTTCCACTCCAATCACTCAACGAGTCACATCCAGGTTGGCTGAAGCAGAAGAACAACCAATGGAAGAAGAAGCCGACGCAGCTGATACCTGGGTCACTGTTTTTGGATTTCAGCCAAGCCAAGTGTCGATTTTGTTGAATTTATTCTCGCGGCACGGGGAAGTAGTTAGTCATCAGACTCCATCAAAAGGAAACTTCATACATATGCGCTATTCGTGTGTCACACACGCTCAACAAGCTATTTCTCGAAACGGAACTCTCCTC

### Recoded sequence (nucleotides 271-936)

TTCACCTCTCAACACTGCGAGCGCGCCATGCAGCGATATTTTCGCCGTTTCTGCGCCTGCCGTACCTCAGCATTTGAAGGATACGCCTGGATCTAAAAGCGTCCATTGGTCTCCATCTCTGGTTCAATCTGGCGAAAAGAGCGCGGCACAAACGCAAAATACGCCGGCTAACCTGTCCTTCGGAGGAAATAGCGCCTTCTCGGCGCCTACGAAGCCAGCCCCGCAGTCCATTCAGACGAGCGCTTTCGGCGGTCAAGCGATGCATGgttagaatttacattttttcgcgaatatttaaaataaaattgattttcagCACCACCGCTTAGGTCTCTCAGAGATAAAGTCGAACCAGCAAAAAAGATCTCTAGAAGGAATACGTTCACTGCCCGCAGCgCCCCACTCTCCgCACCAATCACTCAAAGGGTCACGTCCCGGTTGGCCGAGGCGGAGGAGCAACCAATGGAGGAAGAAGCTGACGCGGCCGACACCTGGGTTACAGTTTTTGGATTTCAACCATCACAAGTAAGCATTTTGTTGAATCTATTCAGCAGGCACGGCGAAGTGGTTTCGCATCAGACACCAAGCAAAGGAAACTTCATTCACATGCGCTATAGCTGTGTCACGCACGCCCAACAAGCCATTTCTAGGAACGGAACTCTTC TC

### Molecular biology

The plasmids and oligonucleotides used in this study are listed in the Key resource table. Gateway cloning was performed according to the manufacturer’s instructions (Invitrogen). All the constructs were verified by DNA sequencing (Eurofins Genomics).

### Biochemical assays

Western blot analysis were performed using standard procedures.

### Protein production and purification

#### GST-NPP-19

The expression of NPP-19 full length N-terminally fused to GST was induced by the addition of 1 mM of isopropyl β-D-thiogalactopyranoside (IPTG) to 1 L Luria Broth cultures of exponentially growing *Escherichia coli* BL21 DE3 pLysS (Invitrogen) strain (OD = 0.6), before incubation for 3 hr at 25°C. After pelleting by centrifugation, the bacteria were resuspended in lysis buffer (0.5 M NaCl, 5% glycerol, 50 mM Tris/HCl pH 8, 1X protease inhibitors (cOmplete, Protease Inhibitor Cocktail, Tablets, Roche)), before lysis by sonication. The soluble portion of the lysate was loaded on 500 μL of GSH beads (slurry) and incubated for 1 hour. The beads were washed with at least 100 mL of lysis buffer and the proteins eluted with 20mM Glutathione solution in 50mM Tris/HCl pH 8. Proteins were aliquoted and flash-frozen in liquid nitrogen and stored at −80°C.

#### 6x(His)-NPP-19NUP53 [1-301]

The expression of NPP-19 [1-301] fragment N-terminally fused to a 6xHis tag was induced by the addition of 1 mM of isopropyl β-D-thiogalactopyranoside (IPTG) to 1 L Luria Broth cultures of exponentially growing *Escherichia coli* BL21 DE3 pLysS (Invitrogen) strain (OD = 0.6), before incubation for 3 hr at 25°C. After pelleting by centrifugation, the bacteria were resuspended in lysis buffer (0.5 M NaCl, 5% glycerol, 50 mM Tris/HCl pH 8, 1X protease inhibitors), before lysis by sonication. The soluble portion of the lysate was loaded on a 1 ml Hi-Trap Column (GE, healthcare) previously loaded with a 1M NiCl_2_ solution. The column was washed with ten volumes of lysis buffer, and bound proteins were eluted in lysis buffer containing 600 mM Imidazole pH 8. After dialysis with lysis buffer to remove imidazole, proteins were concentrated, aliquoted, flash-frozen in liquid nitrogen and stored at −80°C.

#### MBP-NPP-10N^NUP53^ [513-821]

The expression of NPP-10N N-terminally fused to MBP was induced by the addition of 1 mM of isopropyl β-D-thiogalactopyranoside (IPTG) to 1 L Luria Broth cultures of exponentially growing *Escherichia coli* BL21 DE3 pLysS (Invitrogen) strain (OD = 0.6), before incubation for 3 hr at 25°C. After pelleting by centrifugation, the bacteria were resuspended in lysis buffer (0.5 M NaCl, 5% glycerol, 50 mM Tris/HCl pH 8, 1X protease inhibitors), before lysis by sonication. The soluble portion of the lysate was loaded on a 1 ml MBP-Trap Column (GE, healthcare). The column was washed with ten volumes of lysis buffer, and bound proteins were eluted in lysis buffer containing 20 mM Maltose pH 8. Proteins were aliquoted and flash-frozen in liquid nitrogen and stored at −80°C.

#### 6×(His)-PLK-1

To produce *C.e.* 6×(His)-PLK-1, insect Sf9 cells were infected with appropriate baculovirus and then lysed in lysis buffer (PBS, pH 7.2, 250 mM NaCl, 30 mM imidazole, and protease and phosphatase inhibitors [Roche]), passing the cell suspension 30 times through a 21- gauge syringe needle. The lysate was clarified by centrifugation for 10 min at 16,000 g, and the supernatant was injected on HiTrap Chelating HP column loaded with nickel sulfate (GE Healthcare). Proteins were eluted by an imidazole gradient using a fast protein LC Äkta System (GE, Healthcare). Most purified elution fractions were pooled, diluted volume to volume in the lysis buffer without imidazole and containing 50% glycerol, concentrated on a centrifugal concentrator (Vivaspin VS15RH12; Vivaproducts), flash-frozen in liquid nitrogen, and stored at −80°C.

#### GST-Plk1 PBD (H. s) WT and GST-Plk1 PBD H538A/K540M

The human GST-Plk1 PBD WT or phosphate pincer (GST-Plk1 PBD H538A/K540M) mutant fusion proteins were induced by the addition of 1 mM of isopropyl β-D-thiogalactopyranoside (IPTG) to 1L cultures of exponentially growing *E. coli* BL21 DE3 pLysS (Invitrogen) strain (OD = 0.6), before incubation for 3 hr at 25°C. After pelleting by centrifugation, the bacteria were resuspended in lysis buffer (10 mM Tris pH 8, 150 mM NaCl, 1 mM EDTA, 5 mM DTT, 0.05% NP40, 1 mM PMSF, 1X protease inhibitors), before lysis by sonication. The soluble portion of the lysate was loaded on a 1 ml GST-Trap Column (GE, healthcare). After extensive washes, the bound proteins were eluted in lysis buffer containing 20 mM Glutathione pH 8. Proteins were aliquoted and flash-frozen in liquid nitrogen and stored at −80°C.

### Far-Western ligand-binding assay

GST-NPP-19 full-length, GST-NPP-19 [1-301] or MBP-NPP-10N [513-821] fragments, phosphorylated *in vitro* by PLK-1 or CyclinB-Cdk1 as described (*7*) were separated on stain Free SDS-PAGE 10% gel (Biorad). The gel was imaged and then transferred to a PVDF membrane 0.45 µm during 1h30 at 90V. After saturation overnight at 4°C in blocking solution (4% milk in TBS- Tween 0.1%), the membranes were incubated with 2 µg of GST-PBD WT or the GST-PBD H538A/K540M mutant of Plk1 (version of the PBD unable to bind phosphopeptides, negative control) during 5 hr at 4°C. After extensive washing steps (every 15 min for at least 3 hr) with the blocking solution at 4°C, the membrane was incubated overnight at 4°C with human Plk1 antibody (1/1000) in a typical Western blot experiments to reveal the GST-PBD immobilized on the membrane.

### *C. elegans* embryonic extracts and GST-PBD pull-downs

#### Preparation of embryonic extracts

Liquid cultures of N2 worms synchronized at L1 stage were grown on S medium using *E. coli* HB101 bacteria as a food source (*79, 80*). Worms were harvested at the young adult stage by filtration using nylon mesh (Sefar Nitex 35 mm mesh). After several washing steps, embryos were isolated by bleaching, washed several times with M9 buffer, and resuspended in one volume of NaCl 100mM, frozen as beads in liquid nitrogen following cryo-lysis by cryogenic grinding (RETSCH MM400), and kept at-80°C. Three cryolysates were prepared from three independent liquid cultures (A, B, C). For pull-down experiments, 500mg of each embryonic cryolysed were resuspended in 1ml of lysis buffer ((Tris 25mM pH7.5, NaCl 100mM, MgCl2 2mM, DTT 1mM, phosphatase inhibitors (PhosSTOP EASYpack, Roche) and protease inhibitors (cOmplete, Protease Inhibitor Cocktail tablets, Roche) and incubated with 1600U of Benzonase Nuclease (Sigma) for 30min at 4°C. The lysates were clarified by centrifugation (twice 13000 g, 20min at 4C°) to obtain embryonic extracts (roughly 3mg/ml).

#### Preparation of the GST-PBD affinity matrix

GST-PBD Wild-type or GST-PBD H538A/K540M mutant (GST-PBDmut) affinity matrix were obtained by incubating 165 µg of GST-PBDwt or GST- PBDmut proteins with 150µl Glutathione beads in binding buffer (Tris 50mM pH8, NaCl 500mM). After three washing steps the beads were equilibrated in lysis buffer.

#### Loading on GST-PBD beads

500µl of each embryonic extract were applied to GST-PBDwt or GST- PBDmut affinity matrix for 4h at 4°C. After washing the beads with lysis buffer, the proteins were digested on-beads by trypsin, the phospho-peptides were purified by Fe-NTA IMAC and subjected to Mass spectrometry analysis. 50µl of the same embryonic extracts (input) were digested by trypsin, the phospho-peptides were purified by Fe-NTA and subjected to Mass spectrometry analysis.

### Mass spectrometry analysis

#### Material

MS grade Acetonitrile (ACN), MS grade H_2_O, MS grade formic acid (FA) and High- Select™ Fe-NTA Phosphopeptide Enrichment Kit were from ThermoFisher Scientific (Waltham, MA, USA). Sequencing-grade trypsin was from Promega (Madison, WI, USA). Trifluoroacetic acid (TFA), dithiothreitol (DTT), iodoactetamide (IAA) and ammonium bicarbonate (NH_4_HCO_3_) were from Sigma-Aldrich (Saint-Louis, MO, USA). Sep-Pak classic C18 cartridges were from Waters (Milford, MA, USA).

#### Samples preparation prior to LC-MS/MS analysis

150 µg of protein total extracts were precipitated using a six times volume of cold acetone (−20°C). Vortexed tubes were incubated overnight at −20°C then centrifuged for 10 min at 11,000 rpm, 4°C. The protein pellets were dissolved in a 8M urea - 25mM NH_4_HCO_3_ buffer. Samples were then reduced with DTT 10 mM final and alkylated with 20 mM IAA then diluted below 1M urea before trypsin digestion overnight at 37°C (enzyme/substrate ratio of 1/50). Beads from pulldown experiments were incubated overnight at 37°C with 100 µL of 25 mM NH_4_HCO_3_ buffer containing 4 µg of sequencing-grade trypsin.

The digested peptides were desalted on Sep-Pak classic C18 cartridges. Cartridges were sequentially washed with 2 mL methanol, 2 mL of a 70% (v/v) aqueous ACN containing 1% (v/v) TFA and equilibrated with 2 mL of 1% (v/v) aqueous TFA. The digested peptides were acidified with 1% (v/v) aqueous TFA, applied to the column and washed with 2 x 1 mL of a 1% (v/v) aqueous FA. Peptides were eluted by applying 2 x 500 µL of a 70% (v/v) aqueous ACN containing 0.1% (v/v) FA. Eluates were vacuum-dried. Phosphopeptide enrichment was then performed according to manufacturer’s procedure starting from the lyophilized peptide samples. Eluted phosphopeptides were loaded and desalted on evotips provided by Evosep (Odense, Denmark) according to manufacturer’s procedure before LC-MS/MS analysis.

#### LC-MS/MS acquisition

Samples were analyzed on a timsTOF Pro 2 mass spectrometer (Bruker Daltonics, Bremen, Germany) coupled to an Evosep one system (Evosep, Odense, Denmark) operating with the 30SPD method developed by the manufacturer. Briefly, the method is based on a 44-min gradient and a total cycle time of 48 min with a C18 analytical column (0.15 x 150 mm, 1.9µm beads, ref EV-1106) equilibrated at 40°C and operated at a flow rate of 500 nL/min. H_2_O/0.1 % FA was used as solvent A and ACN/ 0.1 % FA as solvent B.

The timsTOF Pro 2 was operated in PASEF mode (*81*) over a 1.3 sec cycle time. Mass spectra for MS and MS/MS scans were recorded between 100 and 1700 *m/z*. Ion mobility was set to 0.75-1.25 V·s/cm^2^ over a ramp time of 180 ms. Data-dependent acquisition was performed using 6 PASEF MS/MS scans per cycle with a near 100% duty cycle. Low *m/z* and singly charged ions were excluded from PASEF precursor selection by applying a filter in the *m/z* and ion mobility space. The dynamic exclusion was activated and set to 0.8 min, a target value of 16000 was specified with an intensity threshold of 1000. Collisional energy was ramped stepwise as a function of ion mobility.

#### Data analysis

MS raw files were processed using PEAKS Online X (build 1.6, Bioinformatics Solutions Inc.). Data were searched against the *Caenorhabditis elegans* Wormpep release 2021_07 database (total entry 28411). Parent mass tolerance was set to 20 ppm, with fragment mass tolerance at 0.05 Da. Specific tryptic cleavage was selected and a maximum of 2 missed cleavages was authorized. For identification, the following post-translational modifications were included: acetyl (Protein N-term), oxidation (M), deamidation (NQ) and phosphorylation (STY) as variables and half of a disulfide bridge (C) (pulldown) or carbamidomethylation (C) (total lysates) as fixed. Identifications were filtered based on a 1% FDR (False Discovery Rate) threshold at both peptide and protein group levels. Label free quantification was performed using the PEAKS Online X quantification module, allowing a mass tolerance of 20 ppm, a CCS error tolerance of 0.05 and 0.5 min of retention time shift tolerance for match between runs. Protein abundance was inferred using the top N peptide method and TIC was used for normalization. Multivariate statistics on protein or peptide measurements were performed using Qlucore Omics Explorer 3.8 (Qlucore AB, Lund, *SWEDEN*). A positive threshold value of 1 was specified to enable a log2 transformation of abundance data for normalization *i.e.* all abundance data values below the threshold will be replaced by 1 before transformation. The transformed data were finally used for statistical analysis *i.e.* evaluation of differentially present proteins or peptides between two groups using a Student’s bilateral t-test and assuming equal variance between groups. A p-value better than 0.01 was used to filter differential candidates.

The confidence of modification sites is estimated by an Ascore, which calculates an ambiguity score as -10×log10(p). The p-value indicates the likelihood that the peptide is matched by chance (Ascore = 13 for a p-value of 0.05).

#### Immunofluorescence and microscopy

Fixation and indirect immunofluorescence of *C. elegans* embryos was performed essentially as described on subbing solution-coated slides (*82*). After Freeze-crack and fixation with cold dehydrated methanol, slides were washed 3 × 5 min, blocked for 1 hr in PBS 2% BSA and incubated overnight at 4°C with primary antibodies diluted in PBS 2% BSA. Working dilutions for the primary antibodies were 1/500 for rabbit LMN-1 and mouse Mab414 antibodies, 1/500 for rabbit NPP-19^NUP53^ antibodies (*31*). Slides were later incubated for 1 hr at room temperature with secondary antibodies, anti-Rabbit (1/1000) and anti-Mouse (1/1000) coupled to the Alexa 568 and 488 fluorophores respectively. Next, embryos were mounted in Vectashield Mounting Medium with DAPI (Vector). Fixed embryos were imaged using a spinning disk confocal microscope with 63 × N/A 1.4 objectives. Captured images were processed using ImageJ and Adobe Photoshop.

For cell cycle timing analysis in live specimens by differential interference contrast (DIC) microscopy, embryos were obtained by cutting open gravid hermaphrodites using two 21-gauge needles. Embryos were handled individually and mounted on a coverslip in 3 µl of M9 buffer. The coverslip was placed on a 3% agarose pad. DIC images were acquired by an Axiocam Hamamatsu ICc one camera (Hamamatsu Photonics, Bridgewater, NJ) mounted on a Zeiss AxioImager A1 microscope equipped with a Plan Neofluar 100×/1.3 NA objective (Carl Zeiss AG, Jena, Germany), and the acquisition system was controlled by Axiovision software (Carl Zeiss AG, Jena, Germany). Images were acquired at 10 s intervals.

Live imaging was performed at 23°C using a spinning disc confocal head (CSU-X1; Yokogawa Corporation of America) mounted on an Axio Observer.Z1 inverted microscope (Zeiss) equipped with 491- and 561 nm lasers (OXXIUS 488 nm 150 mW, OXXIUS Laser 561 nm 150 mW) and sCMOS PRIME 95 camera (Photometrics). Acquisition parameters were controlled by MetaMorph software (Molecular Devices). In all cases a 63×, Plan-Apochromat 63×/1.4 Oil (Zeiss) lens was used, and approximately five z-sections were collected at 1 µm and 10 s intervals. Captured images were processed using ImageJ and Photoshop.

### Quantification and statistical analysis

Tagged NPPs levels at the nuclear envelope were measured in two steps. First, we used the pixel classification workflow from Ilastik software (*83*), which assigns labels to pixels based on pixel features and user annotations. We classified pixels into two labels, one corresponding to the pixels at the NE and the other to the background. For every pixel of the image, Ilastik software estimated the probability that each pixel belongs to each label. The resulting probability maps for each image were then used for quantitative analysis. Once we had generated the pixel classification and the probability maps of each image, we then used the Fiji Software to obtain the fluorescence intensity at the nuclear envelope by using the macro described in Supplementary Materials. The macro consisted of thresholding, making a binary mask, creating a selection of the binary mask by making the region of interest (ROI) of each frame, and measuring the integral density at the nuclear envelope. The average signal intensity of Tagged::NPPs at the NE 350s before anaphase was arbitrarily defined as 1. The raw value were plotted only for the levels of GFP::NPP-1^NUP54^ (**Fig. 5B**) and GFP::NPP-11^NUP62^ (**Fig. 5C**) at the nuclear envelope.

To determine when the NE becomes permeable (**Fig. 3A**), we quantified the fluorescence intensity of mCherry::Histone in the nucleoplasm starting -200 sec before anaphase onset (0 sec) in the different strains analyzed **Fig. 3B**. We performed two separate steps of pixel classification. In the first step, we classified pixels corresponding to the chromosomes in one label and the pixels from the nucleoplasm and the cytoplasm to another label. In the second step, we classified the pixels of total mCherry::Histone signal of both pronuclei to one label and the pixels corresponding to the background to another label. The resulting probability maps of each image were exported and used for quantitative analysis. We then used the Fiji Software and applied the macro described in the Supplementary Materials to obtain the integral density of mCherry::Histone at the chromosomes (value A) and in the nucleus of both pronuclei (value B). We then obtained the integral density mCherry::Histone at the nucleoplasm (value C) by applying the formula (B – A = C) for each time point. The average signal intensity of mCherry::Histone at the nucleoplasm 200s before anaphase was arbitrarily defined as 1. The results are presented as means ± SEM.

### List of strains used in this study

| Name/alias | Description | Genotype | Source | Additional informations |
| --- | --- | --- | --- | --- |
| N2 |  | *C. elegans* N2 Bristol | CGC |  |
| HT1593 |  | *unc-119(ed3)III* | CGC |  |
| OCF3/WLP584 | NPP-1::GFP | *unc-119(ed3) III; ltIs37 IV; jjIs1092. jjIs1092 [(pNUT1) npp-1::GFP + unc-119(+)]. ltIs37 [pie-1p::mCherry::his-58 + unc-119(+)] IV.* | Cohen-fix lab |  |
| BN69 | GFP::NPP-5 | *npp-5(tm3039)/mln1 [dpy-10(e128)] II; bqIs51/ IV.* | Askjaer lab |  |
| PHX1469/WLP799 | sGFP::NPP-7 | *npp-7(syb1469)I* | This study, SunyBiotech |  |
| PHX1475/WLP800 | mcherry::NPP-8 | *npp-8(syb1475)IV* | This study, SunyBiotech |  |
| JH2184 | GFP::NPP-9 | *unc-119(ed3) III; axIs1595. axIs1595 [pie-1p::GFP::npp-9(orf)::npp-9 3'UTR + unc-119(+)]* | (84) |  |
| JH3867 | Neongreen::NPP-10, mCherry::Histone | mNeonGreen::npp-10 (ax4538) III; lmn-1p::mCherry::his-58::pie-1utr(bqSi189) II | Seydoux lab |  |
| JH3872 | NPP-14<sup>OLLAS</sup> | *npp-14::OLLAS (ax4548)I* | Seydoux lab |  |
| TH239/WLP613 | GFP::NPP-11 | *unc-119(ed3)III::ddIs77[npp-11::TY1::EGFP:: 3xFLAG + unc-119(+)]* | (85) |  |
| PHX1526/WLP805 | GFP::NPP-13 | *npp-13(syb1526)I* | This study, SunyBiotech |  |
| BN189 | mcherry::Histone | bqSi189[pBN13(unc-119(+)) lmn-1p::mCherry::his-58] II | (55) |  |
| BN424 | NPP-21::GFP | dpy-10(cn64) npp-21(bq1[npp-21::gfp]) II | This study | Made by injection of N2 |
| BN678 | NPP-21::GFP ; mcherry::Histone | dpy-10(cn64) npp-21(bq1[npp-21::gfp]) bqSi189[pBN13(unc-119(+)) lmn-1p::mCherry::his-58] II | This study | Made by crossing BN189 and BN424 |
| BN1062 | NPP-21::GFP ; mcherry::Histone | npp-21(bq1[npp-21::gfp]) bqSi189[pBN13(unc-119(+)) lmn-1p::mCherry::his-58] II | This study | Made by crossing N2 and BN678 |
| BN1015/WLP824 | GFP::NPP-19 | *npp-19(bq29[GFP::npp-19])II* | This study |  |
| BN1018/WLP894 | GFP::NPP-19, mCherry::Histone | npp-19(bq29[G>F>P::npp-19]) bqSi189[pBN13(unc-119(+)) lmn-1p::mCherry::his-58] | This study | Made by injection of HT1593 |
| PHX1474/WLP801 | mCherry::NPP-22 | *npp-22(syb1474)V* | This study | SunyBiotech |
| BN738 | GFP::NPP-24 | npp-24(bq14[GFP-Frt-KO>unc-119(+)>npp-24]) II; unc-119(ed3) III | This study | Made by injection of HT1593 |
| BN739 | GFP::NPP-24 | npp-24(bq15[G>F>P::npp-24]) II; unc-119(ed3) III | This study | Made by injection of BN738 with Cre expression plasmid |
| BN740 |  | npp-24(bq15[G>F>P::npp-24]) bqSi189[pBN13(unc-119(+)) lmn-1p::mCherry::his-58] II | This study | Made by crossing BN189 and BN739 |
| BN452 |  | *bqSi189 [lmn-1p::mCherry::his-58 + unc-119(+)] II. bq5[GFP::mel-28] III.* | (55) |  |
| WLP820 |  | *npp-7(syb1469)I; mcherry:his58 II* | This study |  |
| UV2059/ WLP758 | LMN-1 8A | *lmn-1(jf140[S21,22,24,32,397,398,403,405A]) I/hT2 [bli-4(e937) let-?(q782) qIs48] (I;III)* | (11) |  |
| WLP893 | LMN-1 8A, GFP::NPP-19 | *lmn-1(jf140[S21,22,24,32,397,398,403,405A]) I/hT2 [bli-4(e937) let-?(q782) qIs48] (I;III); npp-19(bq29[G>F>P::npp-19])II* | This study | Made by crossing WLP758 and WLP894 |
| WLP922 | LMN-1 8A, GFP::NPP-19, mCherry::Histone | *lmn-1(jf140[S21,22,24,32,397,398,403,405A]) I/hT2 [bli-4(e937) let-?(q782) qIs48] (I;III); npp-19(bq29[G>F>P::npp-19]) bqSi189[pBN13(unc-119(+)) lmn-1p::mCherry::his-58]* | This study | Made by crossing WLP758 and WLP894 |
| WLP926 | LMN-1 8A, mCherry::NPP-22 | *lmn-1(jf140[S21,22,24,32,397,398,403,405A]) I/hT2 [bli-4(e937) let-?(q782) qIs48] (I;III) npp-22(syb1474)V* | This study | Made by crossing WLP758 and WLP801 |
| PHX1534 | NPP-19 10A | *npp-19 10A(syb1524)II* | This study | SunyBiotech |
| WLP806 | NPP-19 10A | *npp-19 10A(syb1524)II* | This study | Made by crossing PHX1534 with N2 |
| WLP809 | NPP-19 10A | *npp-19 10A(syb1524)II* | This study | Made by crossing PHX1534 with N2 |
| WLP844 | NPP-19 10A;<br>sGFP::NPP-13 | *npp-13(syb1526)I; npp-19 10A(syb1524)II* | This study | Made by crossing WLP805 and WLP821 |
| WLP821 | NPP-19 10A;<br>mcherry::NPP-8 | *npp-19 10A(syb1524)II; npp-8(syb1475)IV* | This study | Made by crossing WLP800 and WLP809 |
| WLP1107 | NPP-19 10A;<br>GFP::NPP-11 | *npp-19 10A(syb1524)II; unc-119(ed3)III::dds77[npp-11::TY1::EGFP:: 3xFLAG + unc-119(+)]* | This study | Made by crossing WLP584 and WLP809 |
| WLP1106 | NPP-19 10A;<br>NPP-1::GFP | *npp-19 10A(syb1524)II; unc-119(ed3) III; ltIs37 IV; jjIs1092. jjIs1092 [(pNUT1) npp-1::GFP + unc-119(+)]. ltIs37 [pie-1p::mCherry::his-58 + unc-119(+)] IV.* | This study | Made by crossing WLP613 and WLP809 |
| OD2425 | PLK-1::sGFP | *plk-1(lt18) ([plk-1::sgfp] loxp)III* | (7) |  |
| WLP552 | PLK-1::sGFP,<br>mCherry::Histone | *plk-1(lt18) ([plk-1::sgfp] loxp)III, Pmex5 - mcherry-his-11- 3'UTR tbb-2 II* | (7) |  |
| WLP1160 | *npp-19 10A*,<br>PLK-1::sGFP | *npp-19 10A(syb1524)II; plk-1(lt18) ([plk-1::sgfp] loxp)III* | This study | WLP552 and WLP1006 |
| OD999 | GFP::NPP-18,<br>mCherry::Histone | *unc- 119(ed3)III; ltSi245[pNH42;Pnpp-18::GFP- npp- 18; cb- unc-119(+)]II; ltIs37[pAA64; pie-1/mCherry::his- 58; unc-119 (+)] IV* | (86) |  |
| OD1496 | GFP::NPP-6,<br>mCherry::Histone | *unc- 119(ed3)III; ltSi464[pNH103;Pmex-5::npp- 6::GFP::tbb- 2:3'UTR; cbunc- 119(+)]I; ltIs37[pAA64;pie-1/mCherry::his- 58; unc- 119 (+)] IV* | (86) |  |

| Reagent type (species) or resource | Designation | Source or reference | Identifiers |
| --- | --- | --- | --- |
| Antibody | Plk1 (human)<br>(Mouse monoclonal) | Merck Millipore | Cat#05-844<br>RRID:AB_310836 |
| Antibody | GST<br>(Rabbit polyclonal) | This study | L. Pintard Lab<br>(Home made) |
| Antibody | Anti-Mouse IgG (Fab specific)<br>-Peroxidase antibody<br>(produced in goat) | Sigma | Cat#A9917<br>RRID:AB_258476 |
| Antibody | Anti-Rabbit IgG (whole molecule)<br>-Peroxidase antibody<br>(Produced in goat) | Sigma | Cat#A0545<br>RRID:AB_257896 |
| Antibody | Anti-Rabbit IgG (H+L) Cross-Adsorbed Secondary Antibody, Alexa Fluor 568<br>(Produced in goat)<br>Fluor 568 | Invitrogen | Cat#A-11011<br>RRID:AB_143157 |
| Antibody | Anti-Mouse IgG (H+L) Cross-Adsorbed Secondary Antibody, Alexa Fluor 488<br>(Produced in goat) | Invitrogen | Cat#A-11001<br>RRID:AB_2534069 |
| Antibody | OLLAS (Rat monoclonal) | Novus | Cat#NBP1-067113 |
| Chemical compound, drug | Coomassie R250 | Sigma | Cat#B014925G |
| Chemical compound, drug | Ponceau Red | Sigma | Cat#A1405 |
| Chemical compound, drug | VECTASHIELD Mounting Medium with DAPI | Eurobio | Cat#H-1200 |
| Chemical compound, drug | IPTG | Euromedex | Cat#EU0008-B |
| Chemical compound, drug | GLUCOSE | Sigma | Cat#G8270 |
| Chemical compound, drug | Adenosine TriPhosphate (ATP) | Sigma | Cat#A2383 |
| Chemical compound, drug | Imidazole | Sigma | Cat#I202 |
| Chemical compound, drug | Glutathione | Sigma | Cat#G4251 |
|  | Maltose |  |  |
| Chemical compound, drug | HiTrap Chelating HP 5 × 1 mL | GE Healthcare | Cat#17-0408-01 |
|  | MBPTrap HP 5 x 1mL | GE Healthcare | Cat#28-9187-78 |
| Chemical compound, drug | Glutathion Sepharose 4B Fast Flow | GE Healthcare | Cat#17-0756-01 |
| Chemical compound, drug | Pfu | Promega | Cat#M7741 |
| Chemical compound, drug | DpnI | Biolabs | Cat#R0176S |
| Chemical compound, drug | NdeI | ThermoScientific | Cat# 10349709 |
| Commercial assay or kit | ECL reagent | Millipore | Cat#WBKLS0500 |
| Commercial assay or kit | BP Clonase II Enzyme Mix (Gateway cloning) | Invitrogen | Cat#11789-020 |
| Commercial assay or kit | LR Clonase II Enzyme Mix (Gateway cloning) | Invitrogen | Cat#11791-020 |
| Recombinant protein | Human Cyclin B-Cdk1 kinase | New England Biolabs | Cat#P6020L |
| Recombinant DNA reagent | L4440 (RNAi Feeding vector) | (87) | N/A |
| Recombinant DNA reagent | *plk-1* cloned into L4440 | (87) | Arhinger Library |
| Recombinant DNA reagent | *npp-1* cloned into L4440 | (87) | Arhinger Library |
| Recombinant DNA reagent | *npp-10* cloned into L4440 | (87) | Arhinger Library |
| Recombinant DNA reagent | *npp-14* cloned into L4440 | (87) | Arhinger Library |
| Recombinant DNA reagent | *npp-19* cloned into L4440 | (87) | Arhinger Library |
| Recombinant DNA reagent | Gal4 pDEST DB-PLK-1 PBD H542A, K544M | (42) | pMG538 |
| Recombinant DNA reagent | pGEX-4T (GST) | GE Healthcare | Cat#GE28-9545-49 |
| Recombinant DNA reagent | pFasTBAC Hta PLK-1 *C. elegans* | (40) | pLP871 |
| Recombinant DNA reagent | pGEX-6p1 GST-PLK-1 PBD *H. s* | Gift I. Sumara | N/A |
| Recombinant DNA reagent | pGEX-6p1 GST-PLK-1 PBD H538A/K540M *H. s* | Gift I. Sumara | N/A |
| Recombinant DNA reagent | pDONR201 | ThermoFisher | N/A |
| Recombinant DNA reagent | pDONR201 NPP-19 (aa1-378) | This study | pLP1675 |
| Recombinant DNA reagent | pDONR201 NPP-10N (aa513-821) |  | pLP2630 |
| Recombinant DNA reagent | pDEST17 His-NPP-19 (aa1-301) | This study | pLP2352 |
| Recombinant DNA reagent | pDEST17 His-NPP-19 3A (aa1-301) | This study | pLP2361 |
| Recombinant DNA reagent | pDEST17 His-NPP-19 8A (aa1-301) | This study | pLP2353 |
| Recombinant DNA reagent | pDEST15 | ThermoFisher | Cat#11802014 |
| Recombinant DNA reagent | pDEST 15 GST-NPP-19 (aa1-378) | This study | pLP1858 |
| Recombinant DNA reagent | pDEST 15 GST-NPP-19 3A (aa1-378) | This study | pLP2035 |
| Recombinant DNA reagent | pDEST 15 GST-NPP-19 10A (aa1-378) | This study | pLP2143 |
| Recombinant DNA reagent | pMAL Gateway | Thermofisher | pLP354 |
| Recombinant DNA reagent | pDEST MBP-NPP-10N (aa513-821) | This study | pLP2633 |
| Recombinant DNA reagent | pDEST MBP-NPP-10N 3A (aa513-821) | This study | pLP2682 |
| Recombinant DNA reagent | pDEST MBP-NPP-10N 8A (aa513-821) | This study | pLP2637 |
| Sequence-based reagent | Primers for cloning and site-directed mutagenesis (see oligonucleotide sequences table source) | This study | N/A |
| Software, algorithm | Clustal Omega | EMBL-EBI | https://www.ebi.ac.uk/Tools/msa/clustalo/ |
| Software, algorithm | The eukaryotic linear resource for functional sites in proteins | (88) | http://elm.eu.org/ |
| Software, algorithm | Jalview | (89) | https://www.jalview.org/ |
| Software, algorithm | Adobe Illustrator CS6 | Adobe | https://www.adobe.com/products/illustrator.html |
| Software, algorithm | Adobe Photoshop CS4 | Adobe | https://www.adobe.com/products/photoshop.html |
| Software, algorithm | Affinity Designer | Affinity | https://affinity.serif.com/en-gb/ |
| Software, algorithm | Image J | NIH; (90) | https://imagej.nih.gov/ij/ |
| Software, algorithm | ZEN | Zeiss | https://www.zeiss.com/microscopy/int/products/microscope-software/zen.html |
| Software, algorithm | PRISM | Graphpad | https://www.graphpad.com/ |
| Software, algorithm | Metamorph | Molecular Devices | https://www.metamorph.com/ |
| Software, algorithm | Proteome Discoverer 2.2 | Thermo Scientific | https://www.thermofisher.com/store/products/OPTON-30945#/OPTON-30945 |

### List of primers used in this study

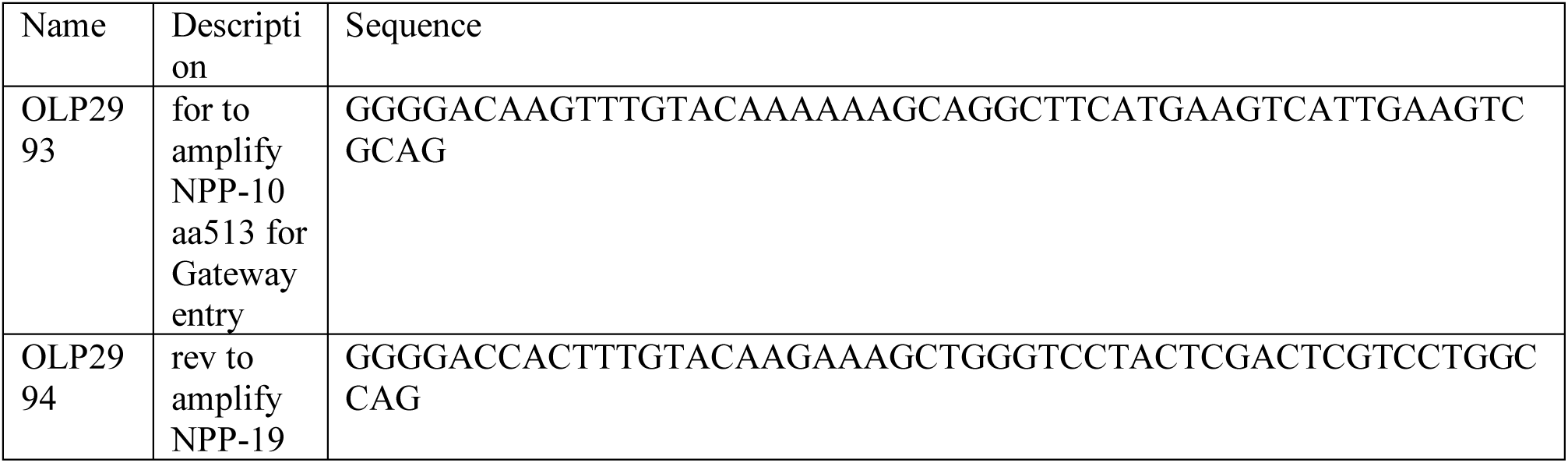
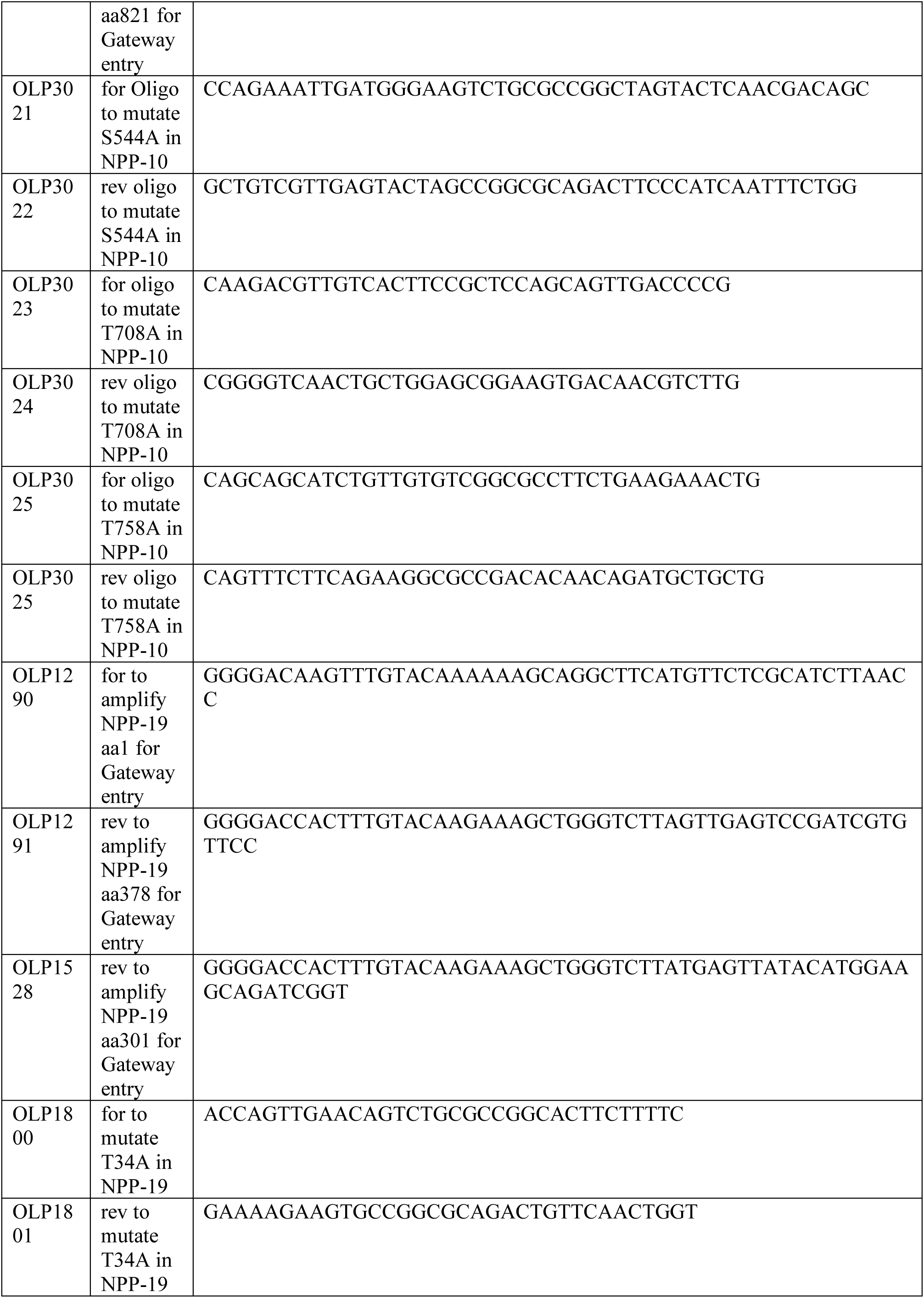
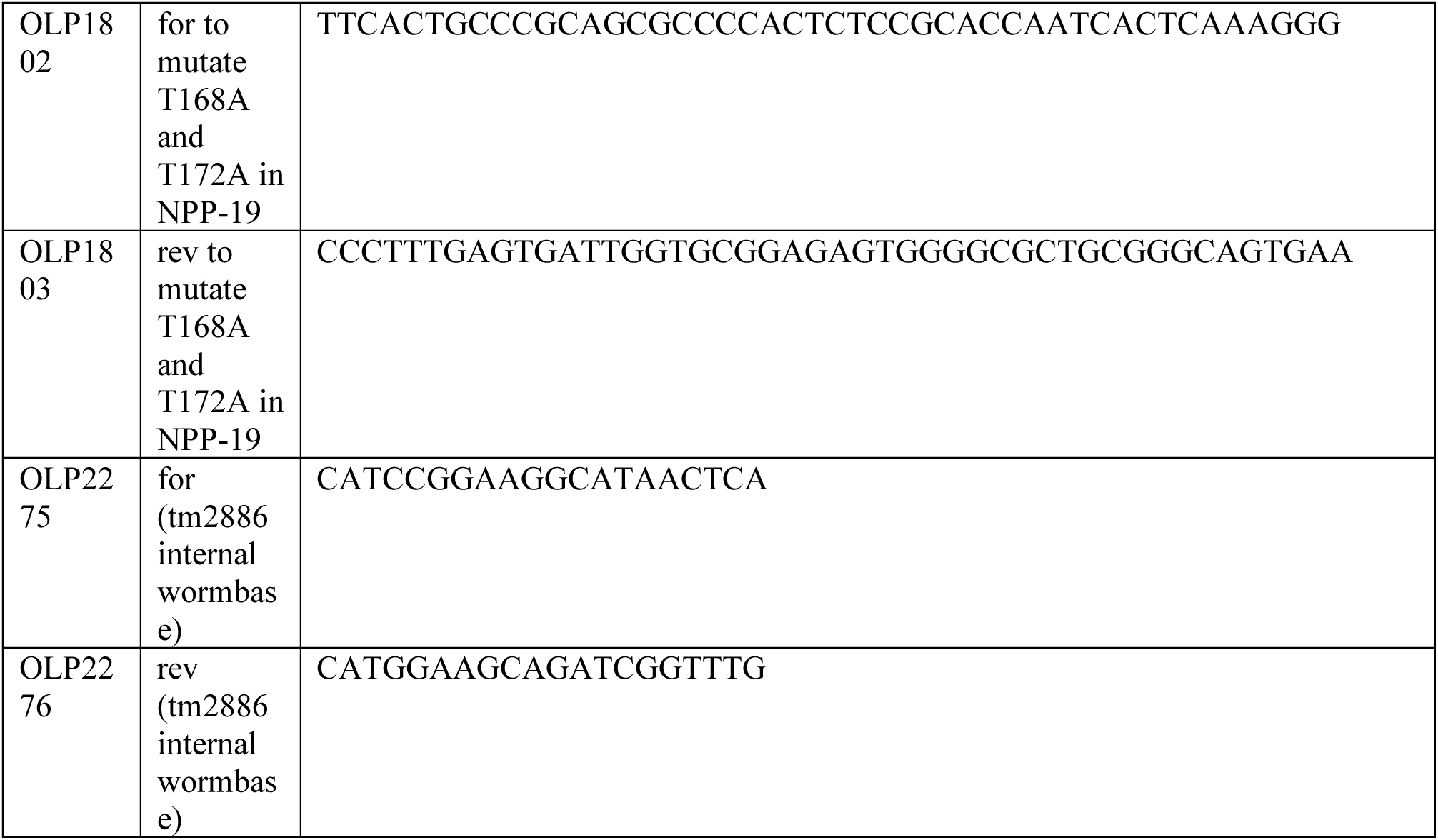

## Acknowledgments

We thank P. Moussounda for media preparation. We thank X. Baudin for microscopy data acquisition and V. Contremoulins for image analysis. We thank R. Karess, N. Joly and V. Archambault for critical reading of the manuscript. We thank V. Doye and all members of the laboratory for stimulating discussions. We acknowledge the ImagoSeine core facility of the Institut Jacques Monod, member of IBiSA and France-BioImaging (ANR-10-INBS-04) infrastructures and the Institut Jacques Monod ‘Structural and Functional proteomic platform”.

## Funding

PhD fellowship from the French Ministry of Higher Education and Research (SMN). 4^th^ year PHD fellowship from the Foundation ARC (SMN).

CONACYT grant CVU 364106 and the SECTEI/162/2021 fellowships (GVA). Laura Thomas is a Life Science Research Foundation Fellow.

Research in the Seydoux lab is supported by the National Institutes of Health (grant number R37 HD37047) and by the Howard Hughes Medical Institute. GS is an investigator of the Howard Hughes Medical Institute.

Agence Nationale pour la Recherche” ANR, France - ANR-17-CE13-0011 (LP). Ligue Nationale Contre le Cancer” Equipe labéllisée, France (LP).

Spanish State Research Agency, the European Union and the European Regional Development Fund (CEX2020-001088-M and PID2019-105069GB-I00; doi: 10.13039/501100011033), (PA).

## Author contributions

Conceptualization: SNN, GVA, BO, LP

Methodology: SNN, GVA, BO, VL, GC, LP

Investigation: SNN, GVA, BO, VL, GC, LP

Resources: LHV, SNN, GVA, CA, PA, LT, GS

Visualization: SMN, GVA, BO, VL, GC, LP

Funding acquisition: GS, PA, LP

Project administration: LP

Supervision: LP Writing—original draft: LP

Writing—review & editing: SNN, GVA, BO, LP

## Competing interests

Authors declare that they have no competing interests.

## Data and materials availability

All data are available in the main text or the supplementary materials.

## Supplementary Materials

Figs. S1 to S6 Tables S1, S2, S3 Data S1, S2

## Notes

### Competing Interest Statement

The authors have declared no competing interest.

