## Supplementary material for "Mechanisms of Nuclear Pore Complex disassembly by the mitotic Polo-Like Kinase 1 (PLK-1) in *C. elegans* embryos": Suppl. Materials

**This PDF file includes:**

Figs. S1 to S7

Data S1 to S2

**Other Supplementary Materials for this manuscript include the following:**

Tables S1 to S3 in excel format

**Supplementary Figures:**

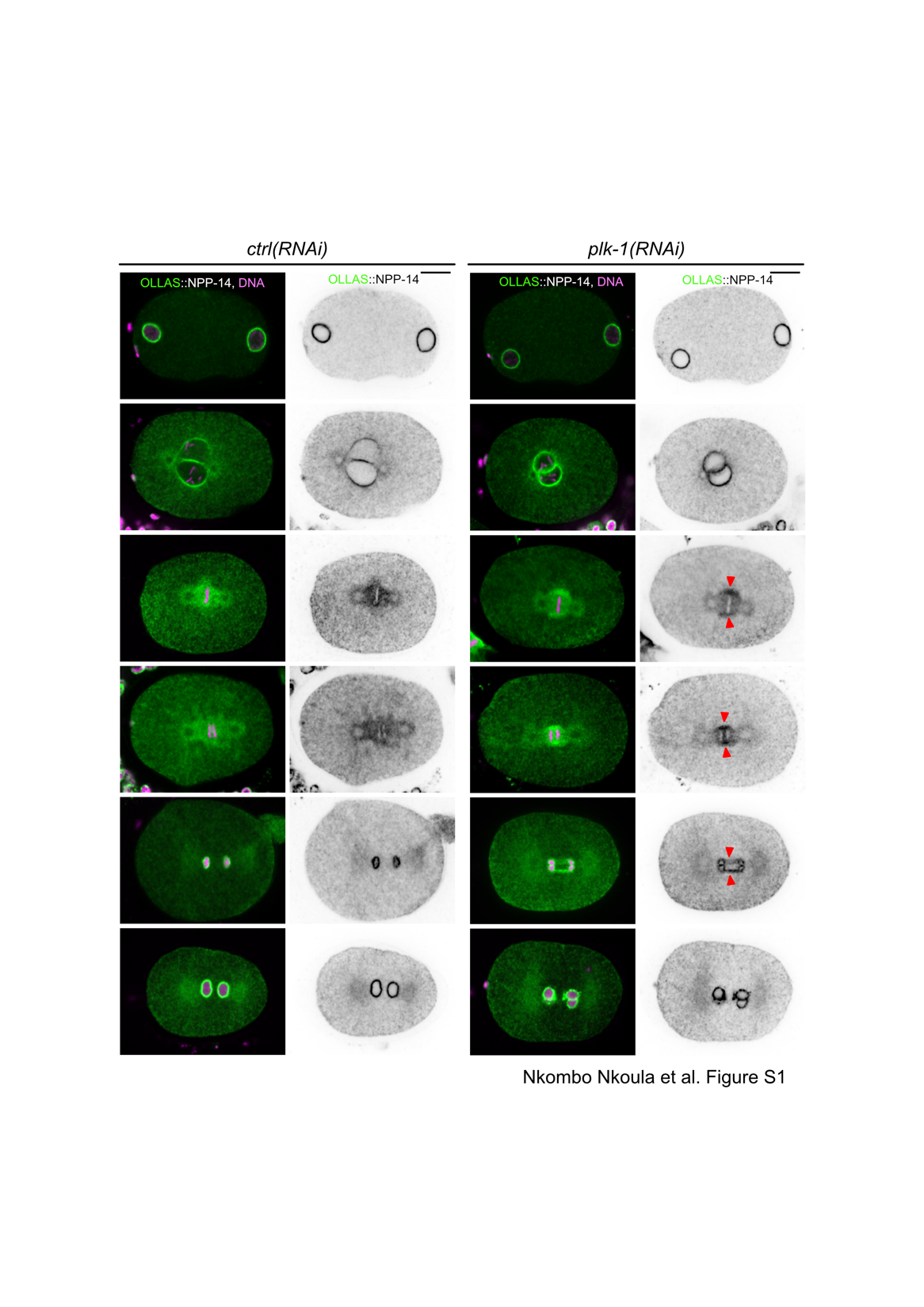

**Fig. S1: A fraction of OLLAS::NPP-14^NUP214^ persists at the NE during mitosis upon *plk-1* inactivation**

Fixed embryos expressing NPP-14^NUP214^ endogenously tagged with the OLLAS epitope exposed to mock (Ctrl: control) or *plk-1(RNAi)* stained with OLLAS antibody (green) and counterstained with DAPI (Magenta). The red arrowheads show persisting OLLAS::NPP-14^NUP214^ at the nuclear envelope during mitosis.

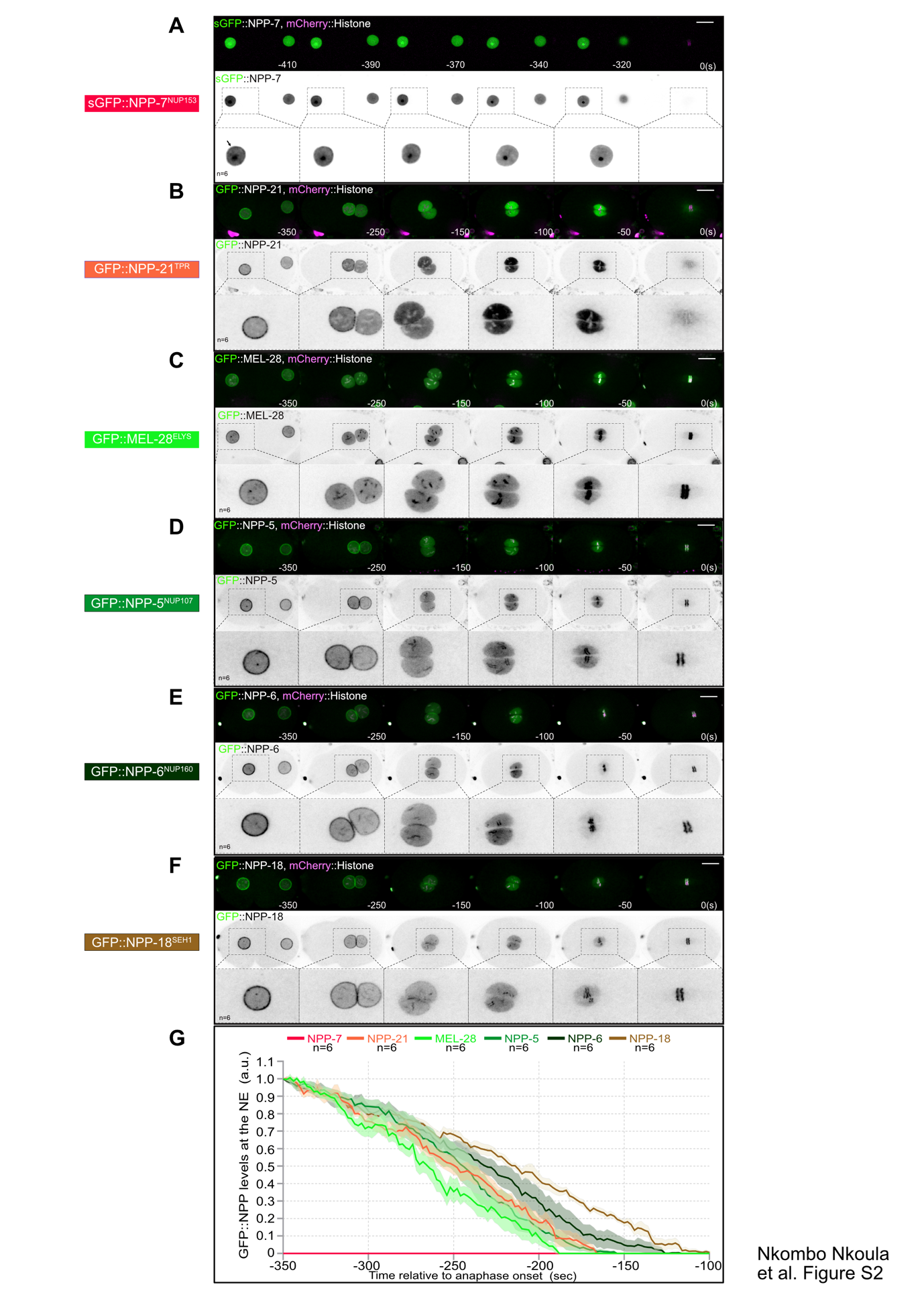

**Fig. S2: Y-complex and nuclear basket nucleoporins dynamics during pronuclear migration**

**(A-F)** Spinning disk confocal micrographs of embryos expressing the indicated tagged nucleoporins and mcherry::histone in one-cell embryos during pronuclear migration. The boxed regions, encompassing representative female pronuclei, are shown at higher magnification beneath. Time in second is relative to anaphase onset (time 0). Scale Bar, 10μm.

**F-** Quantification of GFP::NPP signal intensity above background at the NE in embryos of the indicated genotype during pronuclear migration. The mean +/- SEM is presented for n=6 embryos. Data were collected from three independent experiments. Time in second is relative to anaphase onset (time 0).

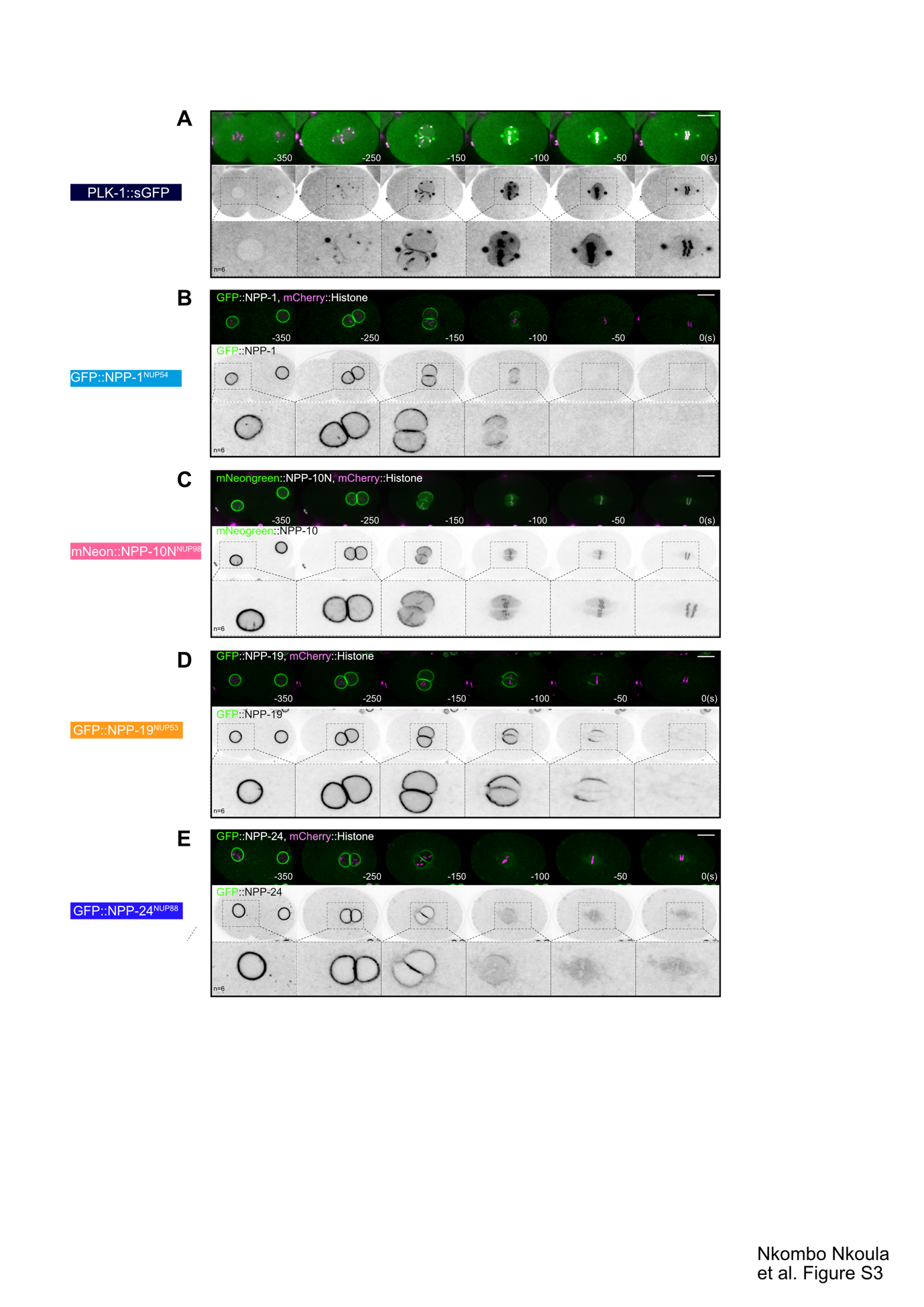

**Fig. S3: PLK-1 and nucleoporins dynamics during the first embryonic division**

**(A-E)** Spinning disk confocal micrographs of embryos expressing PLK-1::sGFP or the indicated tagged nucleoporins and mcherry::histone in one-cell embryos during mitosis. The boxed regions, encompassing representative female pronuclei, are shown at higher magnification beneath. Time in second is relative to anaphase onset (time 0). Scale Bar, 10μm.

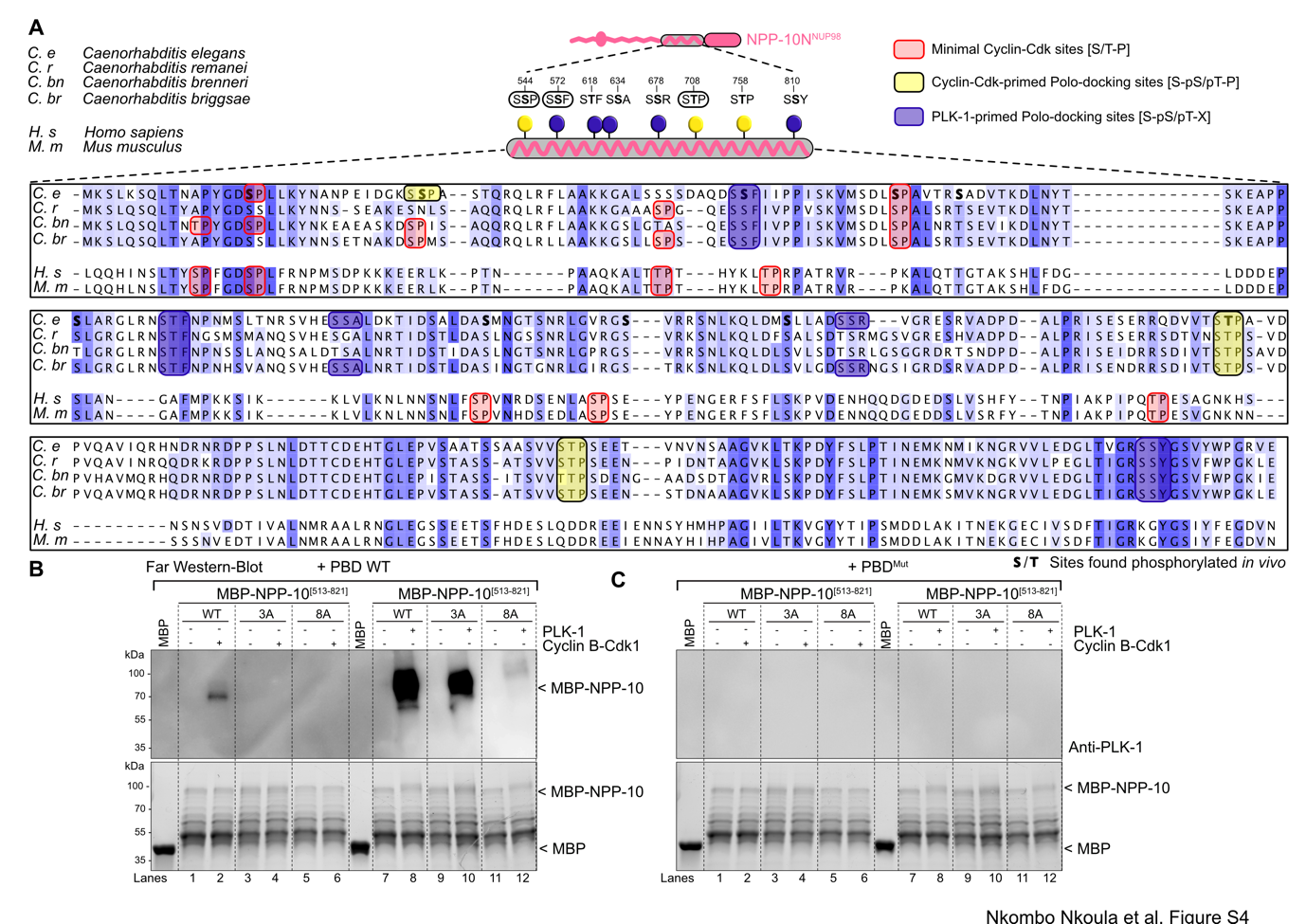

**Fig. S4: NPP-10N^NUP98^ [513-821]** **fragment binds the Plk1 PBD via multiple Polo-docking sites**

**A-** Mutliple sequence alignments of the C-terminal disordered NPP-10N^NUP98^ [513-821] fragment from *C. elegans*, *C. remanei*, *C. breneri*, *C. brigsae*, *H. sapiens* and *M. musculus*.

Sequences were aligned using T-coffee and visualized in Jalview. Sequence features including the minimal Cyclin-Cdk consensus sites as well as the Polo-docking sites matching self or non-self priming are indicated.

**B-C** *In vitro* kinase assays were performed with CyclinB-Cdk1 or PLK-1 kinases and the NPP-10N^NUP98^ [513-821] fragments WT, 3A or 8A tagged with the maltose-binding protein (MBP) as substrates. The samples were subjected to SDS-PAGE, followed by a Far-Western ligand-binding assay using the Polo-box domain WT (PBD WT) (panel B) or the PBD harboring mutations in the phospho-pincers (PBD WT) fused to GST (panel C). The bottom panels show the Stain-Free Blot (Chemidoc, Bio-Rad) of the same membranes.

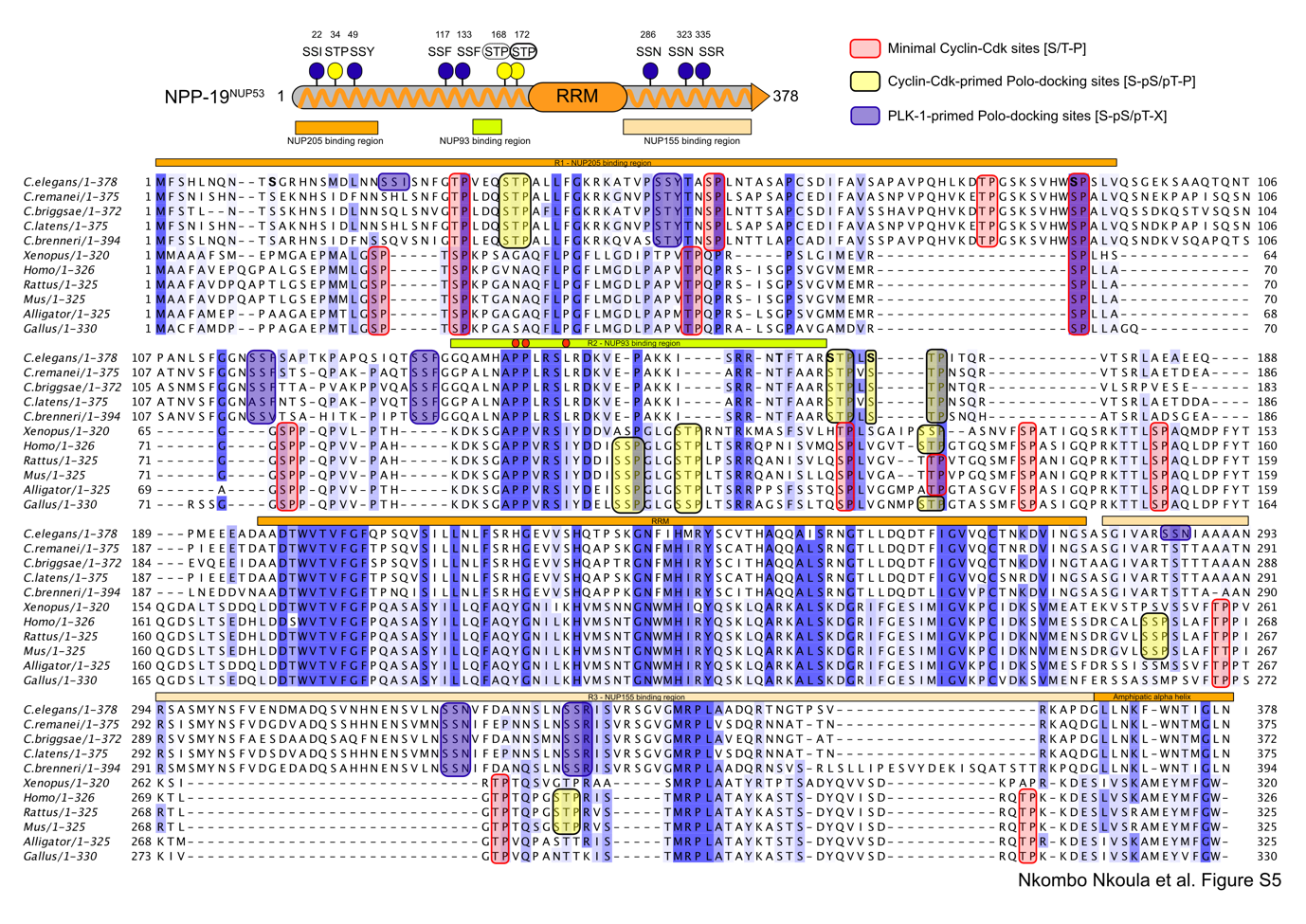

**Fig. S5: Multiple protein sequence alignment of NPP-19^NUP53^**

Multiple protein alignments of NPP-19^NUP53^ from different species. Sequences were aligned using T-coffee and visualized in Jalview. Sequence features including the minimal Cyclin-Cdk consensus sites as well as the Polo-docking sites matching self or non-self priming are indicated.

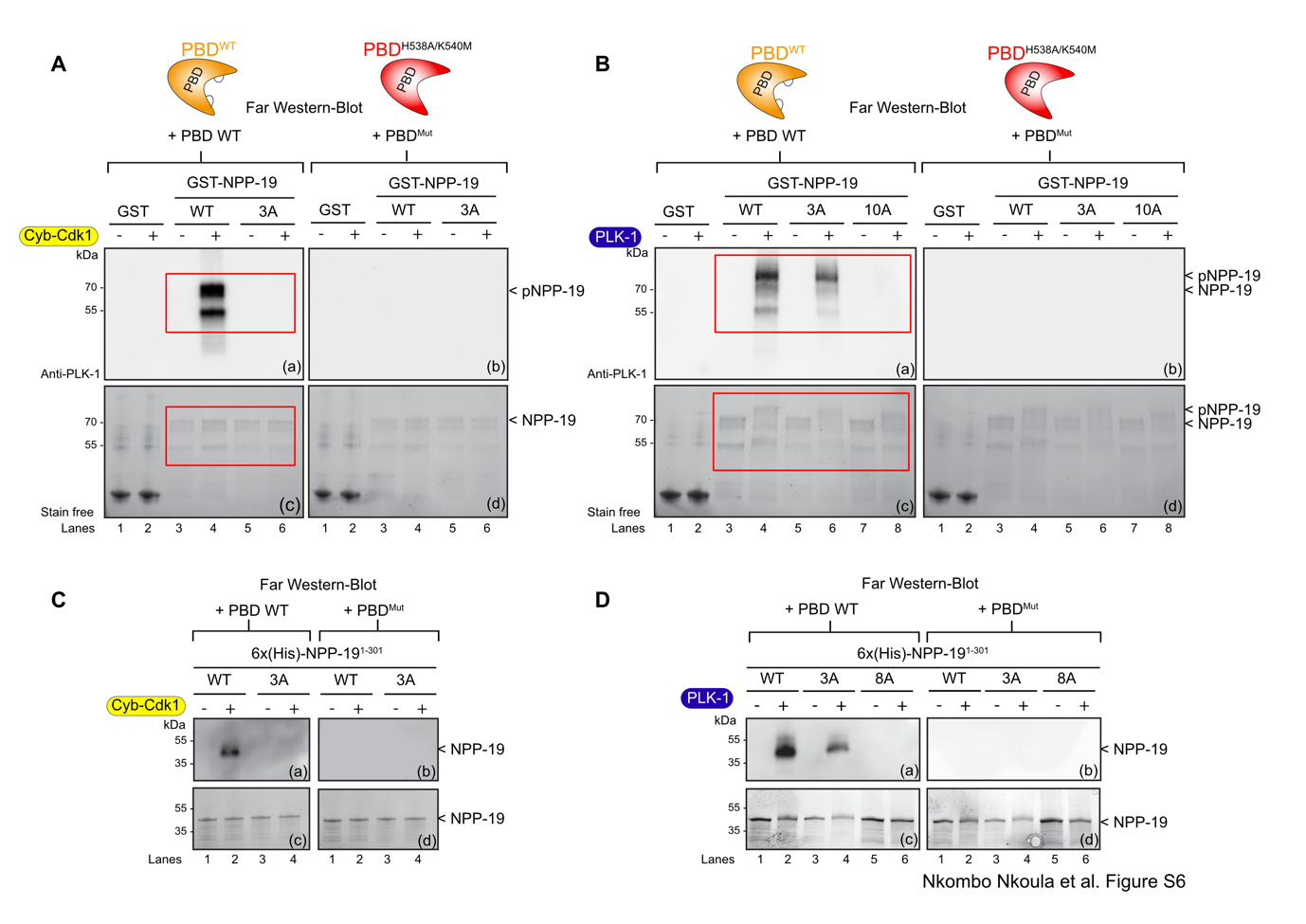

**Fig. S6: NPP19^NUP53^ interacts with the Plk1 PBD in a phospho-dependent manner via self and non-self priming and binding mechanism**

**A-** Full scans of Western blots corresponding to Figure 5B and including the Far-Western blot using the PBD mutated on the phosphate pincer as control (PBD^mut^). *In vitro* kinase assay was performed with Cyclin-Cdk1 and the GST-NPP-19^NUP53^ full-length wild-type or mutant (3A) as substrate. The samples were subjected to SDS-PAGE, followed by a Far-Western ligand-binding assay using GST-PBD wild-type **(a)** or the corresponding phosphate pincer (GST-PBD H538A/K540M) mutant **(b)**. The bottom panel shows the Stain-Free Blot (Chemidoc, Bio-Rad) of the same membrane **(c, d)**.

**B-** Full scans of Western blots corresponding to Figure 5B and including the Far-Western blot using the PBD mutated on the phosphate pincer as control (PBD^mut^). *In vitro* kinase assay was performed with PLK-1 and the GST-NPP-19^NUP53^ full-length wild-type or mutant (3A and 10A) as substrates. The samples were subjected to SDS-PAGE, followed by a Far-Western ligand-binding assay using GST-PBD wild-type **(a)** or the corresponding phosphate pincer (GST-PBD H538A/K540M) mutant **(b)**. The bottom panel shows the Stain-Free Blot (Chemidoc, Bio-Rad) of the same membrane **(c, d).**

**C-** *In vitro* kinase assay was performed with CyclinB-Cdk1 and the 6xHis-NPP-19^NUP53^ 1-301 fragment wild-type or mutant (3A) as substrates. The samples were subjected to SDS-PAGE, followed by a Far-Western ligand-binding assay using GST-PBD wild-type **(a)** or the corresponding phosphate pincer (GST-PBD H538A/K540M) mutant **(b)**. The bottom panel shows the Stain-Free Blot (Chemidoc, Bio-Rad) of the same membrane **(c, d).**

**D-** *In vitro* kinase assay was performed with PLK-1 and the 6xHis-NPP-19^NUP53^ 1-301 fragment wild-type or mutant (3A, 8A) as substrates. The samples were subjected to SDS-PAGE, followed by a Far-Western ligand-binding assay using GST-PBD wild-type **(a)** or the corresponding phosphate pincer (GST-PBD H538A/K540M) mutant **(b)**. The bottom panel shows the Stain-Free Blot (Chemidoc, Bio-Rad) of the same membrane **(c, d).**

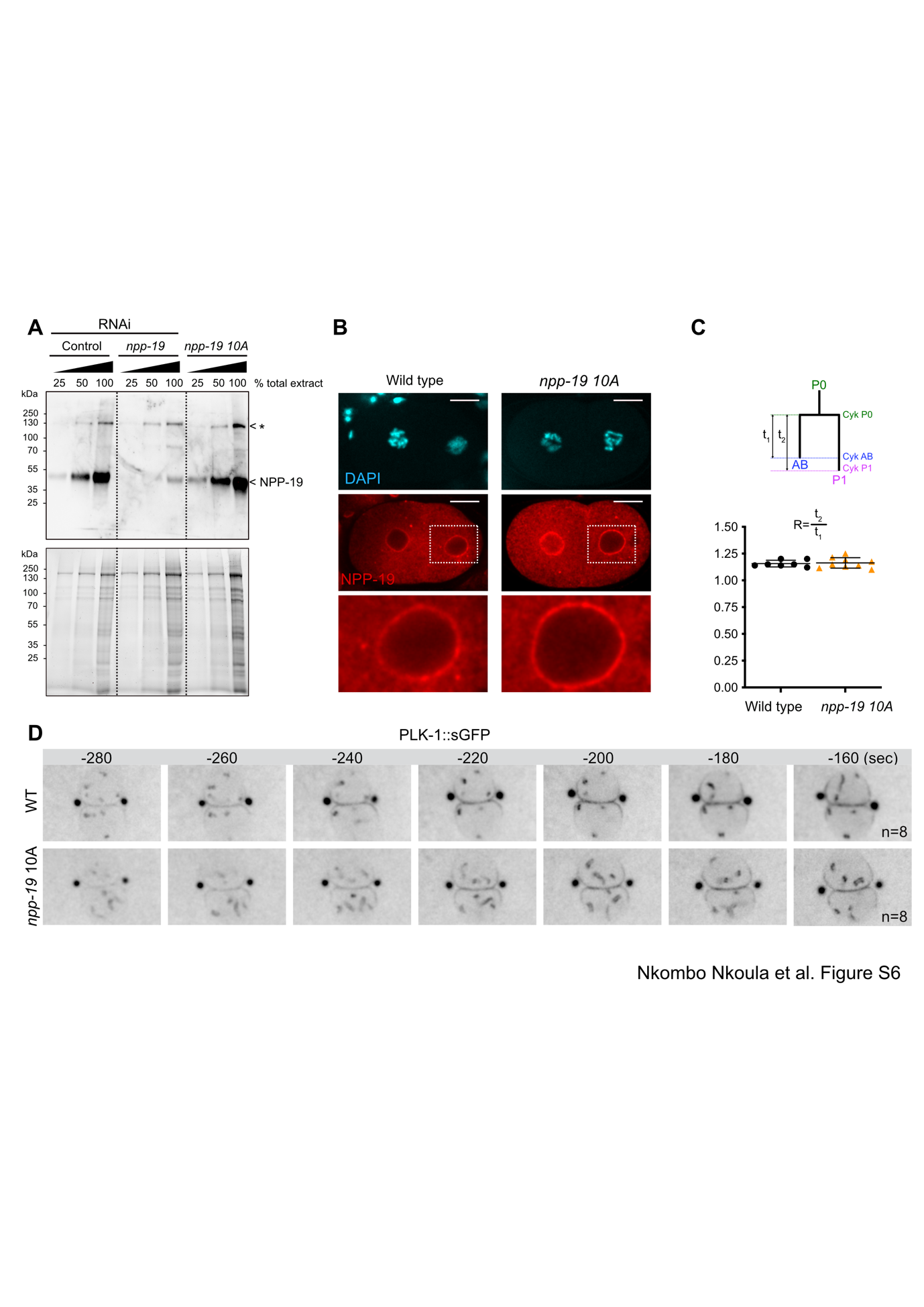

**Fig. S7: NPP-19^NUP53^ 10A is normally expressed in early embryos and localizes to the nuclear envelope**

**A-** Embryonic extracts from control (lanes 1-3), npp-19(RNAi) (lanes 4-6) and npp-1910A embryos were separated by 10% SDS-PAGE and immunoblotted with NPP-19^NUP53^ antibodies (top). Before transfer to nitrocellulose membrane, the gel was stained using tryptophan labeling (stain free, Bio-Rad) (bottom).

**B-** Confocal images of fixed wild-type and NPP-19^NUP53^ 10A embryos stained with NPP-19^NUP53^ antibodies (red) and counterstained with DAPI (bleu). Anterior is to the left in this and other figures. Scale bar: 5 μm. The boxed regions are shown at higher magnification beneath.

**C-** Cell cycle analysis of wild-type and *npp-19^NUP53^* 10A mutant embryos. The mean elapsed time in seconds between ingression of the cytokinesis furrow in P0 (Cyk P0) and AB (Cyk AB) [t1] or between P0 and P1 (Cyk P1) [t2] was determined and the [t2]/[t1] ratio was plotted for wild-type or *npp-19*^NUP53^ 10A mutant embryos.

**D-** Spinning confocal micrographs of wild-type and *npp-19^NUP53^* 10A mutant embryos expressing sGFP::PLK-1 at the one-cell stage. The time indicated in second (sec) is relative to anaphase onset. n corresponds to the number of embryos analyzed.

**Data S1: Phosphopeptides identified on nucleoporins in the total embryonic extracts.**

Unambiguously assigned phosphosites are highlighted in yellow and in bold.

Phosphorylated peptides with unassigned phosphosites are highlighted in yellow.

Phosphosites previously identified by Gnad et al. 2011 (Phosida) are marked by an asterisk highlighted in light blue [*].

Phosphorylated potential Polo-docking sites (S-pS/pT-X or S-pS/pT-P) are underlined.

Full Polo consensus sites (D/E/N/Q-X-pS/pT) are highlighted in dark blue.

[DNE]-[disfavoured PG]-[pS/T]-[FYILMVW] or ([disfavoured PEDGKN]-[FWYLIVM])

> **NPP-1^NUP54^**, isoform c

MSLFGS**ST^*^**PQKQAFTFPTPNAPTTSSGTLFGSTTPSKPLFGSTAQASSTPSLFGTTNTSTPSGGLFGKTGTSTTTTSTAGTLFGAAPTTSTATPSLFGASTTGLFGTSSTTSGGLGGIGSTQQTAKSPVATRLAAFKSGTLGAGSLSGNTPTSFATPSALPGTNAPQSSFLSPANNLNVAPAAYRPPYSTFGGSTPFGAASTGTAAGSTLFGSSTAKPATGGFFGSSSGSTLGGLGATQQQQQPVVQQQQVIQQYHPFVKAVGDPKLFGNDNDGVVAKLNQVAAGLGVGKAPYKDGNQLLSFSMEGNLFERFVGIGYNRISERTDDEGFVTLVLRHPITNLNTEERRDKILEIIKAILGGGPNVEVRYAPGTSMRTLSDGCTEICIIAKEGGFVAGAIKLAQILNDAPKMTQLESQLQVDKTRVLPKVGMSKAQRDRYLETVPDGIDERIWRQAIKENPAPNKLLPVPVRGWEALRDRQKAQVGESKLFHEAINALGNRVEEANHEHADAVVKMEIIRNRHKTLSYRIVRVMLAQWIVSRYSRQIDTDEDVIEAKADTLLAQMNRHNQVKFYVDKFYEILESKPDKLQESMWKMFDMTIEDEHYARRVLTKFVNICSGLYESTHQQIESLEACRRALEG

> **NPP-4^NUP58^**

MSLFGTSTTAPAASTTPLFGSTAAAAPKPGGLFGAPAAAMSSGLFGASQTAPAPASAGLFGSTTAPTPASTSSVPLFGSTTTTASPSGGLFGAKTATTAAPAPTGGLFGASTAAPATGSLFGSTAPATGGLFGAKSTTSAAPTLGGGLFGSSTAPAAAATTSFGAPPATAAAPLGLGGIQQQNSSGGGLPNASGTATSSLTGTGDSKDASKDGEWTQAATLIRDLLPLIEDKFKKNREFMEETENMSIDNVAIEEQIDKTRGWIEEVRRDVLTATQS**^*^S**ERIAYLVSTDKTLCDTTKRVQEHSGSANQTTYMNAIKEHLAELCNFYGNDMDALQERVNFLRNRFEKLLTGEKSITMEELDAYFLRCDANVSNAHHHVTELGKEIEEIRDFLIEQGYTQLRKWSTTAASAAAPIASNSEMIVREGAEFFPSQSSLAIIGSSLRAPAAPAPAVGGLGLGTGTSLFGNTGTTSLFGSTATKPAFSGGSLFGATTSTATSSAAATTTTPTTSLFGSTKPATTTTPFASTVANNSSGLLFSSKK

>**NPP-5^NUP107^**
MTDLFASGDNSSN**S**FDDS**^*^**NDEIARKNETAFKYTTIFHTNLSEKLYHELCALAFGEFDIPQGQISPLMWTRLHEIYSEVEMQTRKLPAGSNYSASSQKYANEAVQIASVLSIYTALAHEYDETVEVSLLSKLVVEDVEFRRIYALLLWSEKAISEQQYEKGLGDKVRKLEGIKSSRINSMMALKRPTFGAANAPKDASLDPDAPGTKEDQALEQMAMNVFFQLIRSGETSKAANLAIDLGMGAIGAQLQLHSMLRNPLDIPLEASKQNFGEYKRSRRAKYYQMTQKLIEQSQGSEDDAYWMLISAIRGNIQPMLKAGKSVIEKVWAYANSAVLARILAAEGAMTQETISTLFNVPLTSKSILDELRSEADRTKEVYILLRVIDDMLNDDIEDLYKFANETVGEFVPNDKNCQVNMLALDIFFHLVAVSYASGFEPNDDGNAVIILGFDDLRARSGTSSHKKMAAFYSRFLPEDMKLPEIVETMKAVDSDEEREILAESLKQSDIDFGRCACTLIEQIRKDDKTKVVTLEEQIDHWHWLLIGGEETALAALEECNRLVRKVMLSTPIDESVIRQIIRKALHFEVPKLLSQAVENEATVLSLITDGTLFEQKPGSQLAINKIEHAALEFYGLCSFVDVNNFMITIALKLGLMFKYTPITDDELSMIGGVKRLDNTTAADWEASLRVRARAEQTLREEVLKKRAAEHNTRVGMVQQHLDTVLPMLRGLVNNIGVRPEYFLSPRANGDPMRVHRKEIQEIRNLFLPQFFILLAQAAVRLDDTTNFNDFFTSFNNDLGLDQEWMVFIKAFYAELNLKVE

**>NPP-6^NUP160^** isoform a

MELVYGSEISFTDGFAIKPARTFVVNTNAPLHSNGFDVQPSAGVATFPHCSNYGDRIFLWRAIGQKLFIEERSLLYSITDGSLCIDFTRTPIIPGTSITIFEEGVLCIVVPTASAIHRFYARLHSKGHDTFSILARIKEDDDFRRFRTSHSLSTPGRPIRASVTHHPSRNTISYCTAEGQLVIVTLGSYTESSDKHEEFTMGEVGLLGKLLGGTDKRVSDACVMKNLRP**T**SGVTRERKISVQSDIVFAVTRDGWVQAWNVETKKQLPSTIDLNNYFASDGKSLNRDLESADFEEEKSPMEPTETVYSIRAYSFDVDILLIVGCDLAVSGKSVGMRVHILKVSNDQIQHLQMFETSMSVDERLVDLELIQNYFPSKESESLDETQDGELYNPLTAALFSLSALFKSSSAKKTYSMKRISFAIQWKKGRVFTDFDWHSVRQFTSPATTKEEPIGNTDEVEPAIERPYNLSADSSIETLIDVVFDTDIYPFDIVFRAVQIVSDNFRGTLGQVRNNNWPELSKLVDTYLTSVEFNRKFQQKTDRSIRLRLNAPQESQKTALKDFWWALLRACEELDFAARGAISLAPIQFCGDLRIMAVVHRDRMTFLGDNNTEFLEIISSENIPKQGNVEEKFRKLPNKHFIDLIEEASKFADRRVFLMNRERARLTNAKNGVAVVEDDNVENAYDYDSDGRFINIKTALEGPIVKLAAAFVEASTFDNSKPFEDPHPQIFGGSFTQSVVSANIRVSVESRVHFALTLQTLLNAITEKKSRAGAPSFGDVEALSCELREIIRVYRELNEQLDIKIVQNGAKMSIGTWLTSDAEGLSMMKKEGGYGPHGYDEIRENDFNWFVGVTTEAAIRALLSSSEILVLPRRLVLQKQYKVLLTILNSYISETRALKPVITFYRGIAYSGTDHPVKALNSFQSCLDAFSEGNNALRKAVYFLLPKRFDVAQGKDPVEELTASEYFLTVVRFLQEHNHAEEVCSVAVKAIENLPIENESVQLISNTLFNHLTNRREWFQSLKLILRTTLRSETRRASIRELLSLMLACGEWEAIATMKFGEHEQVVEDFLRDAACRQSPSEKKHYFELLFAFYVARKDFRKGACAMYEYARHIESTTCMTPELLRRKRDCLAVVLNLQSVLGMEPDDDRAIYDDATDTNLVFPEPSDDEDELITESDSSGENGKLPSGNKDSGSGTGNNSS**^*^**PDNSNGTSSSGTGVDLVALKARQMAYALGSDDMDTDDTEDDMKGGSIPLFKRRKLLVLSEKDIRDELVLCSARVGLLSSDQFKGVPPTDLNELFSLLVDNQHFDEAFDIARQFNLDSHRLFFTMTREAIMIDALRDDLNIEMAATHQAGWVRLNRRHCAAVATAEEHWSVVRGLVDAAQAEWPGDSRPLRGSTEAFLSFKLNVPVWLHSVFENNDANDYLRCLVDYESYSIALQVLSDIVEQETLQASQSNARTWLPYGIIDELMIRSADYIRKMSLKSPEDQVVAAEVANLRKSADQKMLIYFRKVADFEQAQKMSSRFFDK

**>NPP-7^NUP153^**
MSDKSGGFFSSVGRFFSVGA**S^*^**ATKSKDGDKSNDEGTSNSSSKSSPSASPALKNILSVDNEAKIASDETSVNT**S**QNHRIGSRLRYL**S^*^**PTALDRRT**^*^**S**^*^**ASNGLEIDVDAVPDIFFSTPAVSTNRKRQLTELDKQILPQSFRSESVKRSRFLNR**S**LLEPTLNSSMNNERKMDMTWCGELTANHSPASSITNFSLLSRR**S**GATTNGSLSTRTQEIFKKLEGAN**T**PAKEVQRMSMLRAGIARPEKWG**S**FSESKVPNGSAT**^*^**G**T**PPPPLKKAGDAIPSRIQLISKTMGMSARRAPYWTDLTRKRTSSKNGDTGSSDSMKSLNGNFATAELSSLFSLDLPAPKKTASTASTTTINSSNSRKQHASIMKGPDGKPVSRNTFKLS**^*^**DDIEEVDEDSQKLPPIPTTTLQNPQPLKLAPEYAPKRGFLDDLAFSFTAPVDLVTAVGTAKTASTASSHKASESSSESNAESTEKQTSESSGNESDSEESEDSEVDVGEENGSEVKESPQTSADTISSAGGSNQSSKDSSPKVAKDAEPVVVAPAPVVDAGSKKWECQSCFCSWDSTLSECGACGEARPGGSGTGPKSQPKPSEKQLVSNLSSFASNTPSTVKFGFGSGASTTTTIASTTSNTIPFGSGSSVAPLFGAPKTTAPPPTTVPATIPVAPTTIASAPVAAVTSSSNGTRVDWECPDCMVSNKASDDKCPCCSHVKYA**S**AEASSNVFGNRAFKPLSSTGSTISFGVGGSTATTQPSAFAFGLSKTTEVAPTSTPAFGLSAKPAVSSAPVEKPSAPEVPKTAGSLFGNIAKPADSATTSFLAPGAASSTSSLAPTAGSSLFGGSTGSIFNLNKTNTTETAKPGLFGSILDKETHVTTTPVSVALPSSTDSTQSKSAIPTIAPTMSLFGNSSTGSFGGSSMFGNSKTELPKIS**^*^**S**^*^**T**^*^**LS**^*^**FGNPTTAAAPVSTATAEAVKTNVFGSNSASASTSLFGAGSSSTATNLFTTKPADSTTSSIFGKPISFGDDSGTTTTDGAPAKRGLFSSDSQKLQFGGQQKVEMPKFTGFGNPASTASTTSGSLFGGASTNPPMFGGPSSSSIPAFSTSNSSSTSGFPSSTATPFGNAGTTSTSGVFGAFGNKPSQPGLSSSSSTNSLFGQAPADSSNPFGGTSNNGFNFGASSSTTGATAGGGGVFQFGNAATSAPAPTAAPGGGAFQFGGNMSVPQAPAPGGMENAFSYQAPSGVGARKMAMARRRNMRK

**>NPP-8^NUP155^, isoform b**

MSV**S**FMELGSADSAEAAANKVAHHVEQMMETSDFFDRLTQHGTTPVSGLGEKFYVKGAPEFVSTRRIPIPGELQMQMSNIHSEFSMGFFTQISRVWVVIDNNLYMWNYETNDDLAFFDSSDAAILKVSLVNIKPGVFEPEIQYGLVVGTISDICLYPVFDFVENGASSISIDSKRCFKIALDGATVNDISYTSNGRVFYTADDQLFEFVYEKQNGWFGSTNHKCRGVNQTASILGTLISLPFFGSSKEPLDQITIDKSRNIMYLLGRAGTVSVWDLGADGAACAKFLSVPISKIAHEAHILTQFGHDETSFHSITSIKALEASQSAALNLVATTAKGVRLYFSVSTGPQSTMAMFNNSGTPNERNRPQTSVRPQCLRVAHVRFAPGVTPASIYGDGPNGVSVVYADESICAMATANRNTIFAFSNFFYPSSQFFVESTTECDISGHVWEIETVSRCKVKSRPPPRHLVDRVHPHSYFRSQLESGNQKLLVCSNEGVFEFTHVNAVDALREALFDGGVEGNATLQLWQKLGSTEMLCLAFRILTSDAPIDERIRGKAEQILYSLKEAPEIVENEQQRMLDQSTTTSWSPNDSSMAEWRHRMK**T**PLLSS**T**PRTDGRNGNAGGAPPQAHFSSPFSPMLDMSMASGHHNGNMRMSPSRRHDALFYYFSRLVAPVWNDTICEVLNGKQLTITFEPESIQSLKEEIQKLARLMDDYRLVPMMEFNGYSSNMTDRLNHEATSLERHSLIGLRKLIDATLETLSLWLLAYEYNLTAISSGMNPQLLPNFSSRKLAHLVSDGSNLNAELIRAMIKYFLGDEAGTKILSESLRQLCPNLYSEDDACVTFAMEQLEAARKQGPGAARRRLVQSAVEMFKQSIGKVVLASTCQQLAESVEDYEPIVELCLLRAAKDDPKQLALLAYKHGRSGSDAEMLGAEKKREDCYRVITDVLDKLEDEATSEVPRDTAVNRDLMINAVLNSDDQLAHAAVFRWLLTKNKTNVILQSKSPFIEFFLVQEINAGRGQKYFDLLWRFYEKSGNYDKAARLLSKLAENDNWKMGLTQRCAYLSHAILCAQSCKDSTVTTNIDELRDRLDVANVQMRIKDALGCSASASARNQEFVRKLDGPILSLQELLLQYVVPFKLHKIKLSLLHCAGMYVEKHIFETWEDIIQDEFTTAQDEGTLCEQLSNTIGELFSVYRDTKYFPREFVIRRILEIGSGGVIGESVQQQRHILPPSFYPLLCKKINLSNCEFLRTASDEFRAGGDAWWTHNSRGQEYITKVVLKMARTVVRELENMPTAHSRRSTARDCLTHILPFIRRSCDVSASLSLQNLGTELTALQNRLSEFSN

**> NPP-9^NUP358^, isoform a**

MSDQKPNMGRIVASVVDVQQLMTMQFDSLIKSMDSLKIEHQTGVTQLRDDIRRSEDRFQKQLSDLSTNHGKELERLHQIIHTLLARDANPLGSMIPPQQLQQQQQMLILQRQMEMAHVQAAQAQAHAHAQAQAQAQAQSQVMANLLNAAKPAIPVTQPLVATTAQAKSTVPASGVIAPKT**^*^**SPPEVVIPPAKPTFSTPTPAVVPKPATTGFSFGGTNPATSIFGKKPET**^*^**A**S^*^**PVVVPAAKDDEDEEHDEDYEPEGEFKPVIPLPDLVEVKTGEEGEQTMFCNRSKLYIYANETKEWKERGTGELKVLYNKDKKSWRVVMRRDQVLKVCANFPILGSMTIQQMKSNEKAYTWFCEDFSEDQPAHVKLSARFANVDIAGEFKTLFEKAVAEAKSSSNAGKTIDKEIKPAAEVKKEVKQEVVIPSNNKPEETGFGDQFKPKPGSWECPGCYVTCKADEIECACCGTSKDGSVKEKNIFS**^*^**KPS**^*^**ILQPAPG**T^*^**PKVTFGFGASAPAKEPLAQTSQFGGSLSG**S^*^**PSTSSSIFGGG**T**PKGTSVFGGGAANTPTFSFNKPAAAVNATTPSFNFNNPAASTASPATSTTPGNSLFGGGLSKTESTA**S**STT**T**PSFMFAKNSESAFPKPTF**S**FGKQQ**T**PSTTAPAKQEENKQSETPKSVFGSGFTSGGATFAALSAN**S**AK**S**GSIFDAANVKKAQEELAAQKKA**S^*^**IFG**S^*^**KNTTLNTTSATSHDGDETNEDGDGEYEPEVEFKPVIPLPDLVEVKTGEEDEEVMFSARCKLYKYYSDLKENKERGLGDIKLLKSNDNKYRIVMRREQVHKLCANFRIEKSMKLSPKPNLPNVLTFMCQDFSEDASNADPAIFTAKFKDEATAGAFKTAVQDAQSKM

**>NPP-10N^NUP98^, isoform b**

MFGQNK**S**FGSSSFGGGSSGSGLFGQNNQNNQNKGLFGQPANNSGTTGLFGAAQNKPAGSIFGAASNTSSIFGSPQQPQNNQSSLFGGGQNNANRSIFGSTSSAAPASSSLFGNNANNTGTSSIFGSNNNAPSGGGLFGASTVSGTTVKFEPPISSDTMMRNGTTQTISTKHMCISAMSKYDGKSIEELRVEDYIANRKAPGTGTTSTGGGLFGASNTTNQAGSSGLFGSSNAQQKTSLFGGASTSSPFGGNTSTANTGSSLFGNNNANTSAASGSLFGAKPAGSSLFGSTATTGASTFGQTTGSSLFGNQQPQTNTGGSLFGNTQNQNQSGSLFGNTGTTGTGLFGQAQQQPQQQSSGFSFGGAPAATNAFGQPAAANTGGSLFGNTSTANTGSSLFGAKPATSTGFTFGATQPTTTNAFGSTNTGGGLFGNNAAKPGGLFGNTTNTGTGGGLFGSQPQASSGGLFGSNTQATQPLNTGFGNLAQPQIVMQQQVAPVPVIGVTADVLQMQANMKSLKSQLTNAPYGD**S**PLLKYNANPEIDGKS**S^*^**PASTQRQLRFLAAKKGALSSSSDAQDS**S**FIIPPISKVM**S**DL**S^*^**PAVTRSADVTKDLNYTSKEAPPSLARGLRNSTFNPNM**S**LTNR**S**VHESSALDKTIDSALDA**S**MNGTSNRLGVRGSVRRSNLKQLDM**S**LLADSSRVGRESRVADPDALPRI**S^*^**ESERRQDVVTS**T**PAVDPVQAVIQRHNDRNRDPPSLNLDTTCDEHTGLEPVSAATSSAASVVSTPSEETVNVNSAAGVKLTKPDYFSLPTINEMKNMIKNGRVVLEDGLTVGRSSYGSVYWPGRVELKDVALDEIVVFRHREVTVYPNEEEKAPEGQELNRPAEVTLERVWYTDKKTKKEVRDVVKLSEIGWREHLERQTIRMGAAFKDFRAETGSWVFRVDHFS

KYGLADDDEPMDG**S^*^**PPQQALQASSPLQVIDMNTSARDVNNQVQRKKVHKATDAHHQEIILERVPAPAALGDVVPIIRRVNRKGLGGGTLDD**S^*^**REESCIGNMTTEFNESGHDSIIEEGQQPEKKPKLELLADLEYESSRFIRNLQELKVMPKANDPAHRFHGGGHSAKMIGYGKSKLIDIGIVKGRSSHVGWSETGCLVWSAQPRHNQVLFGTIDRTSDVNENTLISMLDVNVHVSETSRKGPSSQSNSVKSSLTSNFVTYSDSYSSMFAKYIDVAQAGGYDGHVSVWKLISALFPYERREGWSFERGEEIGEWLRTEAVKSVPDDRSADTSSNGVWNQLCLGDIDKAFQIAIDNNQPQLATMLQTSAVCPEATVHCFKAQLDNWKKCETLHLIPKETLKCYVLMSGLSHYEWDQDGKNHSINCLDGLNWIQALGLHVWYLRAWTGLEESYDAYQKDVNAGRAASNRGDLPGELIKLACESQHSVEVVLDCAAGENPNDYFLQWHVWSLLYSVGYRTMSKTSETRLHRNYSSQLEASSLSKYALFVLQHIDDDEERSTAVRSLLDRIARFTDNDMFDSISEQFDIPSEWIADAQFSIAKSVDDSTQLFELAVAAKNYLEICRLFVDDIAPTAVVAGDHDALKAACAMVRPFENQIPEWGATGMVYTDYCRLINLIENDAEEELLQDVLESLETRLHAPTISKNSLQKLSLQTIGRVLFEYRADKNTLPEWTKLLGHRQMFKIFRDRSSWGIERFTIEFD

>**NPP-11^NUP62^**
MFGGSAPKPSIFGGTAATTTASSGFSFGNSSTSTANTGGNTNTTGGFSFGSAQPSTGSTGLFGNSTATGSMFGGSSAAAPAPASAGIFGNSGAAAPAPASTSIFGSSANSAAPATVTFGASAPSAGAGMFGANKPAAPTGGLFGSSTSTATTAPTGGLFGSSTAAPSSGLFGSTAAPAAPGGLFGSTSTSTAAPSGGLFGSSAAPTSTAPAPSGGLFGAAPATSNAAPTSGLFGNSAPAATASSGGLFGAAPKPAAPSGGLFGSTAPATTAATTTATSGLFGAPTSAPSSAPATGGLFGASTAPAAATGGLFSIGAASASTPSVGLFGNSSASTTAAAAPATAPAAASTGGLFGATTAAAAAPTSSTTGGLFGSTAAAPAASLPTGGLFGSSTPAKTPAAPTAGLFGASSTTTTSAPATGSLFGTAPATTTATSAAPAASTGGLFGASSTTTPASTAPTGGLFGAATTTAPAAAAPTGGLFGAATTTAPATVGPTGGLFGAATTTTPATAATLGQTPSTVSATPSLPTTTSTTSTLPKPAEATPTLGLGLGLTST**^*^**PLAKGTGWTTSGLKGAATTSAGLKIGA**S**GSD**T**LSEEEIKAGLGGNDTKAFFTALQEVVNSYHSEIAKQERVFHNKMLELNAYDRELITLEPKVLGLYNEMDDLSGSCKKLHFNVASMTSVLNDIEQNVVELENKLSLPEWHTLDYKFPLDSRFASRHDVQRVQIAQMMLNVDSQMKCADFDLDQITKSLNTMQSTVLKTKTETPLEKTELIMKKQLQKLMDLSTQHDATRDKLNKLKDDHNLKTNNSSKA

>**NPP-12^gp210^**
MIILRSLLVLALVQITISYRLNVPRVLLPYHPTVPVSFVLEVTHPTGGCFTWRSTRPDIVSVKRIETNEAGCSDKAEIRSVAKPGTVGSSELSAVIFAEDKGSGTTLSCGVTVDEIATISIETTTKVLFVDAAPARMTVDAFNADGDRFSTLSEIALEWELASTSSNKAKPLRIVPFEQSTYEAPSEIVKLEKNRKKGYLILIEGVGTGTATLTTKFSDAYLQKVAAHNVELAVVANLLLVPSQDVYLPVHSVLPFQVLIVKQRGTEIVNMPNPSYELQIDGGDVASLDKKSSSVRALTIGNTAVHLLSSHVDVRAKAGLRPPSTVIHVVDAESVQWHVSGDNWMLETGKQYTINVELLDEHGNVMFVADNSRFDTHIDEQFLHVDFKSENGTWFLVTPLKPSKTTLRTKFVAIIDAKGNRIAQSGKIGGEQRVTIVDPVRIVPPVIYLPFVSEKRSQIDLTATGGSGLFEWTSEDGHVATVDLLTGRMTANSLGSTKVKATDKRNDQLRDIASVHILEVSGIGFGETVRETFVGDTLTLNIKATGLTSDGLLVEMSDCRNIRAHVQITDNALLRHESSADSSLPMMGTGCGTITFKGLSSGDARVSISYLGHKASIDVAVYEKLSISEESSSIALGSTHPLTVSGGPRPWILDPANFYKTQETKQSQLQVTFENEKVLFKCGSSEVTEAVRLRIGNLKSSTLPLPIHSEITVSICCAKPTRLEIFDKKQRPSKCPLNVHSMLINTNVELVLRGSGVCNGAATPLASINGLSPKWTTSDSGLLTVNRHGIEADATSGKKEGQVTIQAQAGSLSTKYEITVKKGLNVEPARLVLWNEAVSKGTFTITGGSGHFHVDNLPTSDSPVAIALRARSLTVTPKNNGQVNLRISDACLVGQHADASVRIADIHSLAIDAPQFVEIGQEVEVEILAQDETGASFEKEHRPLADAQLDASNNHVILTKVDGLRYTLRANSIGTVSLSASSKSSSGRVLSSRPHTVQIFSPIFLQPKRLTLIPDSKFQLEVVGGPQPTPPLDFSLNNSMIASIEPNALITSSELGYTAITGTVRVGDGHVTLDTVVLRVASLGGIILSASSRKVETGGRVNLRLRGVIAGAEDEEPFAFGGAIYPFKVTWSVSDPSVLFTTHPLGGDVVEPTDNQFAIWFNAIRGGSVTVKAVVELNEKARKHFTGRTSTFTAETTITVEDGLSLVQPEMDINTVRVAPNSQLKMVTAWSQASFSVPSDFSSRIVISADGHLITNGKEGSAAITVRNVNSPDNETVLIPVTVSRVASLDVHPTIELKSAFENSPLIHLPVGAQIQLNVVPRDARGRRLAAASNSINFRPHRFDLTDIVATNSNQTLTITLKTAGDTVLRIGDASNTHIATFIRLSASESILPRAAHKYANDLVVSDVICLQSNIFTVDGSRWSSQSSSEGRISWLDENLGVAQLTKAGNTFIRLHADKQTIHSKISVVLPSSLRFPDGQKPEFVSNDEHSVFVIPVIAATNNTSGSKVSSIYGECTADQIRSFDAIGAPFECQVAFTRRSKIISAVNWLTVSAVFSPVFGYGCEIRRFDSSVSTSSIVVPEELLKDQFDARITAKWISDGTVQVNDATIDVPFHFAFIVEEKELVFSNMNQIEAALSIWAPTYDSKHIVVSGCEGDIVSVEKTSRSSDKHSAKANVFYNIRLNIKSAALFTEHAKKCQISVENTLTGQVIRVPVTVQLLDETAKQVYNALESRGVVDVLLILAHKYSHAIPTLLWTCLVGIIILVIGIYVKMNVFDKTGSFGDNTLNNTTHQTSMASSLSSTNVSLREPVFR**ST**PIAG**S**PQVSLPTARDRLRNQMGSGGDNRLWSY

>**NPP-13^NUP93^**, isoform a

MLQFTEILGRVDHQGFAQLLANPIFQTNRVGQENSQQFTVRNGVEQELCGILGGSEGDWTRQTHSVQKQMMFGERGIELPSRQPKSHTAEDTLTDEGVVDVPEAVMDDIDEEELDEDISRVKVETDRAFFNHMLLSRPAAVPMNPAQAMERDNLPFGGGKFVNKNISDRRELIFGDKLHKFLKNQGKKSLVDLMKEAIDESGTDGALGDVWNDVTSVLNRKTSASRDDLTTTANLVEDACKYLQAVFTEHMQTVVERNLEVAERGGIPGTRGLVNAFLKVGTEESFQPEDDSIDGMPTWQVTYHCVRAGDMKSASETLNRLKSFPQCATLVAALNHVAKHGKLDSELKKKLKVEWRHNLAHTKDKYKRALYAALLGGLDSAALADTLENWIWFKLYPLHVDPQLTDVLFKEVQKAVSVDYGEQYFMSNGPSEFQYFFTALWLSGQFERAIYLLHECGQRVDSVHVAVLAHKLGYLRMSKKSTDEMLVVDQNDSTKCHLNLARLIVAYTKSFELVDVPRSLDYWFLLKGITTPTGSDVFEMAVSRSVYLTGQTDEILGKLTPDGRREKGLIDEYLDDPSEVICRVASDTEITGEWDQAVGLYLLASKPTNAAILLSSEISETLRTENKEKIADLVHVAEQFKKVQRGCQASEYATLSLLVDLAVLFDHCRNEEAEIAYGISTHLRLIPTEPDQVTVIVNEFHMVPQKVREVLPDMCLHLMKCLVDHCIRQSTTQANRGANSATTSMFSSSNRYVKQIKAIVLYSATVPYKFPTHVTSRLLQLQASLGI

>**NPP-14^NUP214^**
MSNEDVAEDVSQVTDFHFHTCRKFRLFSSKSDGYSQNEINIRNRVSQLGVTFVTVNSNQLSCFHTKSLLGYKITRENMNVEVTDLPIKTIRLHGVVLINDMGVNSDGTVLGVLHTKNNDVSVDVFDIKKICTSSSIEPFKPLCTTRVGTEQINQGSCLEWNPAFPDTFAASSTDRSILVAKINVQSPANQKLVGIGKFGAVTTAISW**S**PKGKQLTIGDSLGKIVQLKPELEVVRSQHGPENKPNYGRITGLCWLATTEWLVSLENGTDQDAYLMRCKKDKPTEWIQFHELSYSSSKWPLPPQLFPATQLLVDWNVVIVGNSKTSEISTVGKRDDWQTWVPVEGESIYLPTTSSGKDTVPIGVAVDRSMTDEVLLNPDGSQRHRPSPLVLCLTNDGILTAHHIISTFAAHIPCQMSSQNLAINDLKKLQFDSQKPISAPPSDQTPVTKPSTVFGQKPEAETLKSSLVG**S**PSSVQTPKPSSSLFNPKSIASNIETSQLTE**S**KPS**T**PAAPSSQPKIAS**T**PK**S**EAIPKISDKTLEHKKAELIATKKQVLIERMDKINDSMAGAKDATMKLSFAVGKVKTTIMECADVVRASLGDSKEVMDELKNLILSIERMSDRTQHTVKEMDFEIDEKMELVAGVEDGNQVLEKLRNMSETEKLMRFNKLETAADLLNGKYEECSDLIKKLRMSLSEKESLRKQAIL**S^*^**PLRLSSNLNQLRSGSETELALKVMRNVSKIIMDTREQIQRTELEFVRFQRDVKFQNFKKGKENLNFTQPLEMSSLDGDAPQGKSLTDAESIKVRQALVNQIQKRGIVKTRNVIVESYKKSENSAAMKNDLLDT**S**NL**S**NAILKLSMTPRRVMPSSSLFSA**S**P**ST**PSTKSDAATQADEPPIVKTVVVTVESPAKPIASAPAVSSPLIKLNTTTATTTMT**T**PKVTVPKEEANKTQDQKPIIS**T**PASSSIFSSGSLFGTKTQ**T**PLVSKEESTLTTGVPSLINSSLSI**S**PQEIEKASSKVETLNK**T**EEVKDEKSENEV**T**PDLKSEEPKSLETKVKEEPKPAVQ**T**PVKEEETGSNIQK**T**P**S^*^**FSFNS**T**T**T**PKST**SS**TSSIFGGGLKTQ**T**PSSSNSTNIFGARTTTTA**T**PTPASNTSSIFGGGSKAAS**S**PFGSFGQAGCQPAKTSNPATSTASVTFSFNTGATSASAKPAGFGSFGAGASAKPSSVFGGSVTAPTVPNVDDGMEDDSMANGGGSGGFMSGLGNARTSN**T**SGGNNPFAPKTSTGTSASSSSWLFGGGGNQQQQQQQKPSFSFNTAGSSAQQASAPATGTSSVFGGAPKFGSQPAFGAKPFGGGANAGLSKNASIFGGATSSTTNNPATGGFAQFASGQKTSSLFGGGATPQTNTSIFGGGANTTPAPTSSVFGGGASANANKPTSFTSWR

**>NPP-15^NUP133^**
MSGRDLELTLDRVSSIEYPALVKEAFLNNWHASAHRSEVTSNCASLNDRYCWVLSRNQIFIWERAKSSHRAIIPTQLPLPTSGLPRSVKCVVVYDGVHRGANKTPCPGILVVSPEGVLRHWTSIESQTYIEEVLDINNEVALRVELTDEPIDGKSASFLLTTTSGTVYFLNGKGQDSAKTGALECNKVAGREAHGFRRRL**S^*^**SIMFGGESKESTSLITNSFQHQSKDLLVVTVSPDVLTVYNMYTPCELWSLKTKEFFQPKIASFFEADLKRTPLKVRARLIDAAVFRDGLMILIGGTHEESQSVHMFMVWMSANWQTEQPTGVVWSARVPMNEHRALFSKIDDSIYSNLTLCIPKNTAESKKADRTDGIIIINPYFAVSLYLPFDLAKPKKPESLYRHVSIPPRDQLLGYAICSQYVYIMMLESGVSTIRLLPRGFADSSIYTHEQVVVPSLSVGTDDWPILSELLSEMVASGLPKTPLYQSLHRAFELFAEKHMAESEEELKAIIKMPDQEIARIVSQFLYAIIDYSDAANKTDTELHAKRVLTSRIMLFLKHMGVYERIISSPLGISRGGILSLRVGGTMLGEVSERVAASTAIWTWKTSNETNSAVFDAIIEKVLRIPEVQDLGLKDKDALFGRCGLVHHIPVVAAQQLEKNVIGKTKSHRFEVFHAVCELLSGIKETIISWRNCRTKVAIPKFPIWWTLETFASCYRDVAEKIIEELKNGSSTDSERARLLMYILSIYDFYLSESDSQPDNDKVLQEMIALGKPADAMELAEKHKDFGTLVKNYLTTDVGTRQKTFERYKKMFEKDDFEMYLCDYLKEHGRNDVLLQQGGSRVDAYLDNFKELRYSREIANKQFGKAALTLMSLADAETKSFSKFVEFLTRAYYCACSSIDGTDVSEVLDFYKRRYPEMKHRKRIPTEILKICFGNDLDAMMSVEDMLEWNMAVQPNDEASVEGFARAFHLLADLLAVHPDSDELKKKIDKTWKALVDYDEWNRVRSKEDVEKKTIFGKFCNYLINSYPADKGDSFPIWMPISRRLIFPTDIDTVLDECIANTTGNHLSWIKGHLKWIGEQLCKQALLPKSAFFRPDMKQVGSISQAALEAFGPILQRREQRFIDQLNRDSMMET

**>NPP-16^NUP50^**
MNSLIPPPTSEQQMNMFRLRDKMSLLNAEFLKVINGYFTEKNHYDFSGTMKSYMDHVAQLKQIYKVDDDVAADMTVPRRTENSSESSGETVAPRKIAKAVRKNGTPKNPLNS**^*^**TVFAAS**^*^**S**^*^**PAATVASVPKFGDI**S**VITKETPAPLAKTAEPLVAPAAPA**T**ARKRAIRGGGPLGGAE**S**VVFKSGEDGQAATSSVKIPATTIKFPEPTKDFWTKKSDAPAAPSNSGSLFAFLGKDGDKPKETPKFSGF**S**FGKKPAEPSEEPKAADS**T**PKLTFG**S^*^**PKEADLPKPASSLFGASPSKPLVFGGSSADSTTSAPKPFSALSTAASLFGSSSASTTTTATQPLSFGSSSTGGSSLFSGFAGLAQKAMENQNQAKPEGSGEDDEGEYVPPKVETVENQEPDAVLSSKVSVFKFTGKEYTKLGVGMLHIKDNDGKFSVLIRAATATGTVWLNSLCNKAMKATVVDAKGDRIRLTCPSSSTEMATMMIRFGTADGAKKFTDKILEVAV

**>NPP-19^NUP53^** isoform b

MFSHLNQNT**S**GRHNS**^*^**MDLNNSSISNFGTPVEQSTPALLFGKRKATVPSSYTASPLNTASAPCSDIFAVSAPAVPQHLKDTPGSKSVHW**S**PSLVQSGEKSAAQTQNTPANLSFGGNSSFSAPTKPAPQSIQTSSFGGQAMHAPPLRSLRDKVEPAKKISRRN**T**FTARSTPLS**T**PITQRVTSRLAEAEEQPMEEEADAADTWVTVFGFQPSQVSILLNLFSRHGEVVSHQTPSKGNFIHMRYSCVTHAQQAISRNGTLLDQDTFIGVVQCTNKDVINGSASGIVARSSNIAAAANRSASMYNSFVENDMADQSVNHNENSVLNSSNVFDANNSLNSSRISVRSGVGMRPLAADQRTNILQG**T**PSVRKAPDGLLNKFWNTIGLN

**>NPP-21^TPR^** isoform b

MDVDAPLQAPEQPVADADDEEANWEMEKAEMKRIEFNQNRELTDMRERVEDVSRSNSRMLEELTRHAAEIKEHINRQRVLESTRNELTDKNLELETVVSRLKIEKEERDAAVVNATKLAQTAQVETFALKDEIKKLTNEQASLRHSKEALEKEIQGIQFERQKYATERSLHAESKTWLMQEVSERDNKVSSLRLELSNKDIQGANERLQYVQQINLLNSQVENLNEKLDMLKFTNADLIKRMENTELSKVSEIANLEEEIRCQTELQRVMKSSMEESKNAADLFKDQLEAQENVLVEVRKVLQEHQDEMERENLAHADAIKHRDEELAQTRAELVKVTEMMKSMSDVKLNV**S^*^**EEELSELAPAAAETVRYLRGGQ**S**LSSLVLEHARVRGKLTEVEEDNVNLRNTLEELLETIDQNKPQMISQKMVTDELFDKNNRFEKQLDLAESERRQLLSQRDTAQRDLAYVRAELEKYQRDYEFVSKRNAELLYAVERQSRMQDPNWSEQADEQLFQNIVQLQRRNVELESDIENAKASAAQAAINAQSEEMAQLRADLAVTKKSEAELKTKVEQTKAAFDSLKERTEHFKELVRDSVTAAEARTARLRAEEAIAAKVVADATIERLRTQAEDYKADHLRREQDLEQRIRNTEANIASVTETNIKLNAMLDAQKTNTASMDQEFKSALKEKENIFEELKKVTAVNAENEQRLVDLGRQTLEAVEQAGSLRVRVRSLEDELQSARTEINSLQFTANGQRNILEKEEQVRMSVVEMANFLSRVEAERLTHANTQLDVLRLERDSLKASTTRLSDQLTHTKNESKLVQQRLEKELEIARQRLSEKETQVTRDEMELADLRSKLASMHSQYTGSDASGM**T**PDRLKREYMQLKTRTQFLESELDDAKRKLLESETTQKRMDAEHAISASHNTVLEENLKQSEQMGVMEKERLVAKAKCFEDRSKQLAESLEQNQKKLDELRSKNDEQLFAHERETNELRRQLQVASLNLDGVRRELEVVNNNLISMQNEATRNSSALEQHTTIVRQFEDRITEIESANLRLQTELNNKCAALVAESTAKREADQMIEHAERLLQKKTEELNSIEEENRQKQAEYDEKLAQLSLQYESLSANLTNQNTTMEVKVNTDGSSSTVENLQSLLQFVRQSKDEATSRAMTAEVEMRRLRAETAEYERGRNELLRKIRDLETEKIATTAALVEKASLMEKIQALTDVHNINAKLTEEKTKLQAQLHQIQKEKADLENQRSRLSASNEEQKLKIASSDQEANQRKREIEQLKQRVQTNARGAASQPQLDQLKAQLATARQESAAATAKAKAAEDKFNQTRQLAIKYRNENTELKKLAEAPPGEGDPCAARLKLQFDDFTAKINDYKTEIENLNMKVLRMGILEKSLKNTNDQINQLKQENLKLTENIRMAQLQSVSATDVESKPGPSSASKSVSSIRQ**T^*^**PTKVLDPLS**S**AAKQPNEPDQTTGLKTSQQPPSFAAKRPS**^*^**FAPTPTSQQKVSPVKRPIPPSIPNEPLDIIPPVPSDNIPDPTPPTNSFGTVLPVPHTFQTSVRVPTQSLFSSSSTTTVQPQPEKKNVLPSIDSAPSTPGGNSSMVTTTSSMAPGQSIFGNIGNVPVPTTAPTDNLALPEESVIEGSAGQSSLVSGSIDQRKVQDIDLVANDGE**S^*^**RDS**^*^**T**^*^**NVGGV**SSS**DVRRKRT**^*^**ANDFEL**S**EAKRLRE**S^*^**PNET**^*^**VTS**^*^**SETRQQSNVADIPELDDDDGVLGMEHEVSDEDPNDNTIQEQRPDVIDLENDEEVLEDEMDEEEDDDSFGNDEEFEDEEEIPEDDDDDDVVVLSDGDDEPANDNDEESLNDIDDDDGIEEIEMVEESNNRDIEEVLGGED**S^*^**QP**S^*^**LDDQDREAASAVEEAEDEGRDPLGTIDEPSAPADPTGAAGIGSSGRMGQDVQRVRLPTGLRDAEREDQCSSRFFSNETNDERPAERLTARNLARMQRPTRGAKPTRGVYTPARGNRGGRGGGTA

>**NPP-22^NDC1^**
MMGDSHSSFTTTTDEHLYNQFSPGRRKNDFPAAS**S**S**S**S**S**PNLRRSPNRTVSSPRVQQKPITIFDQIVDWFQAEISVRKRLAGAACGYLSTIFFIVTVSILKLTIWAPFSSVQDSLAWWIYPNAWASIIFVGIASVAMSLFSIIKFCKVDQLPRLAATDTFALAGVALEFVTRLTFVYTAFCVADFSFSREFAFVAISLAIAISSALVVFRSDYQLNFSHIQVNSVKTLIDFGTSLPYANISEICGIDAAISYTAAVALILVVGPMVSGFSAWWLLLNIPFHVVLFGLCFTQQFYSKISMKIVNQIVMKPISFPFPPPYTVHSPTPEQTRTLPNVIETDDSLLKFFALHDLRTIAWNDEKRRVDVFSLSQPGKHPRNWKAVSLPCVRMLDELCSRMTVSAARLVGYSWDDHDIENEDVPRDALLMPRKMREMAYRGTGQSRQQKSMAPIRSHNTQTVGLLSKISNFLGFGVTEKLVISRFDAHMNAYAAEALYMLVVDSMGEDRFGVVQKDLKDLITLLCKLIAAIDTYERAKASVADKSDVTFLRIVDASLKSSLQRVVTTFGSHLSSLNLPEEHSRTIRMICLTDEL

>**NPP-23^NUP43^**, isoform a

MAVVTIT**T**DTED**S**EIPQRTEVGSSTPLVDEIMQHADVKLSKILFTGETSSQIISLGKGRGRCISLWERDDGIDPFKVLATKNSEIDPNDACTMTDNRVCIGYADGSLAVFSTDKDDLALMSRIPSIHSGSASRKICRHGNSILSASSNGSLVAVDVETGMPRTIFTGQAGIRSVCTTFGTNVVMAGDANGQITMWDLRENNEHSNTLDPIKTLIPSKKALDAVTALCSHPAQSNLVCCGTDDGIVGLIDARNVRGANITSTYLVAKKAISQVMFHPKCGDNLLVSSNDGSLIRIDASGAPIAVGPRSTKDTIWLEGDLVNSLRLDPIRNESVHSISSFDIRSDTVVTASSIGLISLYQQLPFFPNSNFGRF

**>NPP-24^NUP88^**
MLITHLIDSLSDRAVITPLTPTSVIIYDKNELKVYVGFTNNRNSLEYTKEVVLLLSTEIPKKAEIREIIVSKNGDYVILEGPRSLFVVRIGAEILVAKPDRLPSECFCECYPLHDSLLLQNISLSVVKVRLLPEKCDEKTFVTAVLFSDNCIRFYNLQKKFDSLLLAVDFRNHLHQVHDENVANN**T**FGLQKALVSFDLIPPKPNTSHFSIISIDSDCDFYTSFVHFSCFKEGYAPRIHRIEPVDGLPCDPLDLRYIQTTNPRICSVFVLVSGGGVLSHLVVFPNEFGEFRFLVKDQLRLPSSNGDPRIVQNQIRSLKVSRYEIATSSSLFSVNIFPWFEALTSISPTSTLEKETRVSELVDAVIPSDELSNTTKWTGARALRAVSVQLTQSLATEEEELLPESENIMHLVILENKDGQPAHLFNISTFDNIWSTENKTSFGRDSVSQPPMKSTGSLEQQLAALKPLAACVISEKVSCEEAIDAAMKFFDAVDERLKKHCEISKLFVERCLAVSSSAQALDEKQQSVDQRLIEETNTVEELKIRMHETKERMEGARKGINVLFHRVDENVPLSDNEIRIFERLKEHQKMLSDMTKLVPKMTLDSNEIHRMANIVLKKRGTGEEQNRFAAVEKNATEIESLEARENKLNTGISELSI

>**NPP-26^GLE1^**
MDLVKVDDDVVREMEIRRQQHSKKYDRKFEELVKSMSEAHVEPEKMKIPFSRKPTEQETNNLREEWEKEYAALCTSGRTLARPNPNSSIFSKILAQG**S**PSSAKAGTTRSTPRTSKPPSADENIIFGQKLPETPAPKLLCPPRASPGSSRRLSIFVENLPENHAEIVRGTIFAGREAKIMQNDPENVPSTSTYGEPIRPYLLFARYISLEKGLLDDKNRYKSATDPRFREFLKENIKYKVKQSIKHRTSSEACAQILNYFKTLLQKQEVQVLGGYAPVLQLKTNKDIEYASLCIVHNYVILAERDESLIPLISSHITRLSVFYGYIEQVFCIILLSRSALLRQHPEECMVKFLATSMNPSSPDAIKKWSRQKAHIILFCSVFTKNAYCFSKGFKLKLTDEILWKYVDASTAHILEIPRGSFILWQLITTCKPRLQVDKERWIQLLLNIKFKIIPELDLDSFDGDEAMLMQLKQTIDELLHP

>**MEL-28^ELYS^**
MDNENSSIFKSYQGYECWRGEKQIILKDSIGRQLPYIVNFKKNTCQIFDIEWERVTHSFVFPEGCALIDADYFPTEEGKLGILVGVEDPRQSCGAEHFVLALAVDPDSPAMTITHSLEVPSKITVVKTLFSSADMADETQRTVLKLYHRLMTWQHIVAIGCKETQCYLARLVAVETPSSPVITVHSEKKYLINLMNAYVSGSVLQYTLDDGAYREYPTAAVYISALSLMPRSRTLLVGLSMGGILAASLNPSNQMMLLELRHERLVRKIAPLEPEDDPDKFEYFIATVDCSPRHPIMIQLWRGSFKTLEDVDGEEKYDRPSFSVCLEHKILFGERWLAVNPIVTERDHMMLTRKRGTEDSMHNV**S**QTFGSTSNRNSVLLAYERKKMVIGTEDPNAEPEYIVEAAIFDIDSWYYKRVPGRVSTDGTVLKQCAFLSTIKSNIRSEDVNDIGILTNEATDVSSFSSMVSDADQLFYPSALSFERVFVAKNTRIDWMKIQNIQDTILNKCAVKLPALIRNPEMISSVVMAAGLVRKNILSGSPNSSAAEINELQLSSDQKVLLNVIVYYGKIEEFCQLASRPDISDTLKRELAEWALHEAVDYKRTISDKMVSLFQGRSLALSPLAEESIAQGIKLFRVVYEYLKACSKALKDDRLRNLAHSVICMRNHTKLTSQFINFAIIPVDPIRQQRMKDLHSKRKNMARKNSSSLPVQSVVRKMNRQAPNAQFWNDIPHDEWYPPTPLDLLECLLNVSISESIKRELVVQYVIDWISTSPEDSEHSEKQLALETIKIMTNQMLNVNLEKIYYILDQGKKALTSSKTSDDMRALGEKVFSMKDDEISYEKLWGKDAPMTVTIGKHDLQRFEQRMKMQMEGGKVRLPVLDPESEILYQMFLFENEKFEAMSSEAISSNKLLSAFLPGMIKKDGRGRQKTAKEQEIEISVKKMFERKVQNDDEDMPEVFASVNDKTERKRKSSQFGEDDESSVSSSQYVPPTAKRIQQWKSAVESVANNSSINSITSPDSHQNAEINMMIATPARYYKRHNEEENVQDGFL**S^*^**PAGNRPPPVSAHNSILKTAKGGQSASRGRIRFRADVPRGADE**S**IEDNGRKGLALNFAILEDEEEETMTIRKSRSMGKHDEEKDSEKNVVDEMEEVKDQEQENDECIESEKTFENQDDFEVLEDTSAPEAANTENGSETPPMEDTFEVRDDDVMPPTDETYLSHLQTDKTGILEEEGEDEDIWDGVQR**S**FEVQMDEDCEAVPTIDVADDLESKSEEVNEEEVVESEEVQQDAKEPEKTEKRQEEPEPEVMQPVIPEEPQNESLESSIKLQEELQEEPDIVPT**^*^**GDEDTADKVQEQAVEEDRPP**S^*^**RNTRSSSVQKSTSQVEDRDPKELVEEERPP**S^*^**RN**T^*^**RSASVQKSSNQEKTSESGEVTEEDRPP**S**RNTRSASVQKSSSKVKDQKPEELIEEDRPP**S^*^**RN**T**RSASAQKTVAANK**S**VLESEIP**S**RSASRRTRSTSLRNDTVAEPDETSVAM**T**TRRRTRAT**S^*^**EVVSKQSSEDDGRSTPKTGRTPTKKAAASTSSSRAGSVTRGKKSIIQKMPSPLEVTMEVQEEEEEEAEEERPASRSTRSASVKNTTVDPSSSALA**S**TKRTTSRKRGN**S^*^**ETIDFNQDDK**S**APT**T**PKRGRPAKKDAGSPKVGSKARGTKPKSIFENQEDEEDRSS**^*^S^*^**PDIEQPA**T**PTRSSKRTARSRANSESIDDDSKQK**T**PKKKNAAVNEAGTSKQSRSVTRSRAS**^*^**S**^*^**IDVQQEVEEPT**T**PKRGRGRPPKTVLENIEEGEEERKE**T**AA**T**PLLRSARRAKQ

**Data S2: Phosphosites identified in GST-Plk1 PBD Pull-downs**

Unambiguously assigned phosphosites are highlighted in yellow and in bold

Phosphorylated peptides with unassigned phosphosites are highlighted in yellow

Phosphorylated potential Polo-docking sites are underlined

> **NPP-1^NUP54^**, isoform c

MSLFGSSTPQKQAFTFPTPNAPTTSSGTLFGSTTPSKPLFGSTAQASSTPSLFGTTNTSTPSGGLFGKTGTSTTTTSTAGTLFGAAPTTSTATPSLFGASTTGLFGTSSTTSGGLGGIGSTQQTAKSPVATRLAAFKSGTLGAGSLSGNTPTSFATPSALPGTNAPQSSFLSPANNLNVAPAAYRPPYSTFGGSTPFGAASTGTAAGSTLFGSSTAKPATGGFFGSSSGSTLGGLGATQQQQQPVVQQQQVIQQYHPFVKAVGDPKLFGNDNDGVVAKLNQVAAGLGVGKAPYKDGNQLLSFSMEGNLFERFVGIGYNRISERTDDEGFVTLVLRHPITNLNTEERRDKILEIIKAILGGGPNVEVRYAPGTSMRTLSDGCTEICIIAKEGGFVAGAIKLAQILNDAPKMTQLESQLQVDKTRVLPKVGMSKAQRDRYLETVPDGIDERIWRQAIKENPAPNKLLPVPVRGWEALRDRQKAQVGESKLFHEAINALGNRVEEANHEHADAVVKMEIIRNRHKTLSYRIVRVMLAQWIVSRYSRQIDTDEDVIEAKADTLLAQMNRHNQVKFYVDKFYEILESKPDKLQESMWKMFDMTIEDEHYARRVLTKFVNICSGLYESTHQQIESLEACRRALEG

**>NPP-7^NUP153^**
MSDKSGGFFSSVGRFFSVGASATKSKDGDKSNDEGTSNSSSKSSPSASPALKNILSVDNEAKIASDETSVNT**S**QNHRIGSRLRYL**S**PTALDRRTSASNGLEIDVDAVPDIFFSTPAVSTNRKRQLTELDKQILPQSFRSESVKRSRFLNRSLLEPTLNSSMNNERKMDMTWCGELTANHSPASSITNFSLLSRRSGATTNGSLSTRTQEIFKKLEGAN**T**PAKEVQRMSMLRAGIARPEKWGSFSESKVPNGSATGTPPPPLKKAGDAIPSRIQLISKTMGMSARRAPYWTDLTRKRTSSKNGDTGSSDSMKSLNGNFATAELSSLFSLDLPAPKKTASTASTTTINSSNSRKQHASIMKGPDGKPVSRNTFKLSDDIEEVDEDSQKLPPIPTTTLQNPQPLKLAPEYAPKRGFLDDLAFSFTAPVDLVTAVGTAKTASTASSHKASESSSESNAESTEKQTSESSGNESDSEESEDSEVDVGEENGSEVKESPQTSADTISSAGGSNQSSKDSSPKVAKDAEPVVVAPAPVVDAGSKKWECQSCFCSWDSTLSECGACGEARPGGSGTGPKSQPKPSEKQLVSNLSSFASN**T**PSTVKFGFGSGASTTTTIASTTSNTIPFGSGSSVAPLFGAPKTTAPPPTTVPATIPVAPTTIASAPVAAVTSSSNGTRVDWECPDCMVSNKASDDKCPCCSHVKYASAEASSNVFGNRAFKPLSSTGSTISFGVGGSTATTQPSAFAFGLSKTTEVAPTSTPAFGLSAKPAVSSAPVEKPSAPEVPKTAGSLFGNIAKPADSATTSFLAPGAASSTSSLAPTAGSSLFGGSTGSIFNLNKTNTTETAKPGLFGSILDKETHVTTTPVSVALPSSTDSTQSKSAIPTIAPTMSLFGNSSTGSFGGSSMFGNSKTELPKISSTLSFGNPTTAAAPVSTATAEAVKTNVFGSNSASASTSLFGAGSSSTATNLFTTKPADSTTSSIFGKPISFGDDSGTTTTDGAPAKRGLFSSDSQKLQFGGQQKVEMPKFTGFGNPASTASTTSGSLFGGASTNPPMFGGPSSSSIPAFSTSNSSSTSGFPSSTATPFGNAGTTSTSGVFGAFGNKPSQPGLSSSSSTNSLFGQAPADSSNPFGGTSNNGFNFGASSSTTGATAGGGGVFQFGNAATSAPAPTAAPGGGAFQFGGNMSVPQAPAPGGMENAFSYQAPSGVGARKMAMARRRNMRK

**>NPP-10N^NUP98^, isoform** b

MFGQNKSFGSSSFGGGSSGSGLFGQNNQNNQNKGLFGQPANNSGTTGLFGAAQNKPAGSIFGAASNTSSIFGSPQQPQNNQSSLFGGGQNNANRSIFGSTSSAAPASSSLFGNNANNTGTSSIFGSNNNAPSGGGLFGASTVSGTTVKFEPPISSDTMMRNGTTQTISTKHMCISAMSKYDGKSIEELRVEDYIANRKAPGTGTTSTGGGLFGASNTTNQAGSSGLFGSSNAQQKTSLFGGASTSSPFGGNTSTANTGSSLFGNNNANTSAASGSLFGAKPAGSSLFGSTATTGASTFGQTTGSSLFGNQQPQTNTGGSLFGNTQNQNQSGSLFGNTGTTGTGLFGQAQQQPQQQSSGFSFGGAPAATNAFGQPAAANTGGSLFGNTSTANTGSSLFGAKPATSTGFTFGATQPTTTNAFGSTNTGGGLFGNNAAKPGGLFGNTTNTGTGGGLFGSQPQASSGGLFGSNTQATQPLNTGFGNLAQPQIVMQQQVAPVPVIGVTADVLQMQANMKSLKSQLTNAPYGD**S**PLLKYNANPEIDGKS**S**PASTQRQLRFLAAKKGALSSSSDAQDS**S**FIIPPISKVMSDL**S**PAVTR**S**ADVTKDLNYTSKEAPP**S**LARGLRNSTFNPNMSLTNRSVHESSALDKTIDSALDA**S**MNGTSNRLGVRG**S**VRRSNLKQLDM**S**LLADSSRVGRESRVADPDALPRI**S**ESERRQDVVTS**T**PAVDPVQAVIQRHNDRNRDPPSLNLDTTCDEHTGLEPVSAATSSAASVVSTPSEETVNVNSAAGVKLTKPDYFSLPTINEMKNMIKNGRVVLEDGLTVGRSSYGSVYWPGRVELKDVALDEIVVFRHREVTVYPNEEEKAPEGQELNRPAEVTLERVWYTDKKTKKEVRDVVKLSEIGWREHLERQTIRMGAAFKDFRAETGSWVFRVDHFS

>**NPP-12^gp210^**
MIILRSLLVLALVQITISYRLNVPRVLLPYHPTVPVSFVLEVTHPTGGCFTWRSTRPDIVSVKRIETNEAGCSDKAEIRSVAKPGTVGSSELSAVIFAEDKGSGTTLSCGVTVDEIATISIETTTKVLFVDAAPARMTVDAFNADGDRFSTLSEIALEWELASTSSNKAKPLRIVPFEQSTYEAPSEIVKLEKNRKKGYLILIEGVGTGTATLTTKFSDAYLQKVAAHNVELAVVANLLLVPSQDVYLPVHSVLPFQVLIVKQRGTEIVNMPNPSYELQIDGGDVASLDKKSSSVRALTIGNTAVHLLSSHVDVRAKAGLRPPSTVIHVVDAESVQWHVSGDNWMLETGKQYTINVELLDEHGNVMFVADNSRFDTHIDEQFLHVDFKSENGTWFLVTPLKPSKTTLRTKFVAIIDAKGNRIAQSGKIGGEQRVTIVDPVRIVPPVIYLPFVSEKRSQIDLTATGGSGLFEWTSEDGHVATVDLLTGRMTANSLGSTKVKATDKRNDQLRDIASVHILEVSGIGFGETVRETFVGDTLTLNIKATGLTSDGLLVEMSDCRNIRAHVQITDNALLRHESSADSSLPMMGTGCGTITFKGLSSGDARVSISYLGHKASIDVAVYEKLSISEESSSIALGSTHPLTVSGGPRPWILDPANFYKTQETKQSQLQVTFENEKVLFKCGSSEVTEAVRLRIGNLKSSTLPLPIHSEITVSICCAKPTRLEIFDKKQRPSKCPLNVHSMLINTNVELVLRGSGVCNGAATPLASINGLSPKWTTSDSGLLTVNRHGIEADATSGKKEGQVTIQAQAGSLSTKYEITVKKGLNVEPARLVLWNEAVSKGTFTITGGSGHFHVDNLPTSDSPVAIALRARSLTVTPKNNGQVNLRISDACLVGQHADASVRIADIHSLAIDAPQFVEIGQEVEVEILAQDETGASFEKEHRPLADAQLDASNNHVILTKVDGLRYTLRANSIGTVSLSASSKSSSGRVLSSRPHTVQIFSPIFLQPKRLTLIPDSKFQLEVVGGPQPTPPLDFSLNNSMIASIEPNALITSSELGYTAITGTVRVGDGHVTLDTVVLRVASLGGIILSASSRKVETGGRVNLRLRGVIAGAEDEEPFAFGGAIYPFKVTWSVSDPSVLFTTHPLGGDVVEPTDNQFAIWFNAIRGGSVTVKAVVELNEKARKHFTGRTSTFTAETTITVEDGLSLVQPEMDINTVRVAPNSQLKMVTAWSQASFSVPSDFSSRIVISADGHLITNGKEGSAAITVRNVNSPDNETVLIPVTVSRVASLDVHPTIELKSAFENSPLIHLPVGAQIQLNVVPRDARGRRLAAASNSINFRPHRFDLTDIVATNSNQTLTITLKTAGDTVLRIGDASNTHIATFIRLSASESILPRAAHKYANDLVVSDVICLQSNIFTVDGSRWSSQSSSEGRISWLDENLGVAQLTKAGNTFIRLHADKQTIHSKISVVLPSSLRFPDGQKPEFVSNDEHSVFVIPVIAATNNTSGSKVSSIYGECTADQIRSFDAIGAPFECQVAFTRRSKIISAVNWLTVSAVFSPVFGYGCEIRRFDSSVSTSSIVVPEELLKDQFDARITAKWISDGTVQVNDATIDVPFHFAFIVEEKELVFSNMNQIEAALSIWAPTYDSKHIVVSGCEGDIVSVEKTSRSSDKHSAKANVFYNIRLNIKSAALFTEHAKKCQISVENTLTGQVIRVPVTVQLLDETAKQVYNALESRGVVDVLLILAHKYSHAIPTLLWTCLVGIIILVIGIYVKMNVFDKTGSFGDNTLNNTTHQTSMASSLSSTNVSLREPVFRSTPIAG**S**PQVSLPTARDRLRNQMGSGGDNRLWSY

**>NPP-14^NUP214^**
MSNEDVAEDVSQVTDFHFHTCRKFRLFSSKSDGYSQNEINIRNRVSQLGVTFVTVNSNQLSCFHTKSLLGYKITRENMNVEVTDLPIKTIRLHGVVLINDMGPSSSLFSASVNSDGTVLGVLHTKNNDVSVDVFDIKKICTSSSIEPFKPLCTTRVGTEQINQGSCLEWNPAFPDTFAASSTDRSILVAKINVQSPANQKLVGIGKFGAVTTAISWSPKGKQLTIGDSLGKIVQLKPELEVVRSQHGPENKPNYGRITGLCWLATTEWLVSLENGTDQDAYLMRCKKDKPTEWIQFHELSYSSSKWPLPPQLFPATQLLVDWNVVIVGNSKTSEISTVGKRDDWQTWVPVEGESIYLPTTSSGKDTVPIGVAVDRSMTDEVLLNPDGSQRHRPSPLVLCLTNDGILTAHHIISTFAAHIPCQMSSQNLAINDLKKLQFDSQKPISAPPSDQTPVTKPSTVFGQKPEAETLKSSLVGSPSSVQTPKPSSSLFNPKSIASNIETSQL**T**ESKPSTPAAPSSQPKIAS**T**PK**S**EAIPKISDKTLEHKKAELIATKKQVLIERMDKINDSMAGAKDATMKLSFAVGKVKTTIMECADVVRASLGDSKEVMDELKNLILSIERMSDRTQHTVKEMDFEIDEKMELVAGVEDGNQVLEKLRNMSETEKLMRFNKLETAADLLNGKYEECSDLIKKLRMSLSEKESLRKQAIL**S**PLRLSSNLNQLRSGSETELALKVMRNVSKIIMDTREQIQRTELEFVRFQRDVKFQNFKKGKENLNFTQPLEMSSLDGDAPQGKSLTDAESIKVRQALVNQIQKRGIVKTRNVIVESYKKSENSAAMKNDLLDTSNLSNAILKLSM**T**PRRVMPSSSLFSA**S**PS**T**PSTKSDAATQADEPPIVKTVVVTVESPAKPIASAPAVSSPLIKLNTTTATTTMT**T**PKV**T**VPKEEANKTQDQKPIISTPASSSIFSSGSLFGTKTQTPLVSKEESTLTTGVPSLINSSLSISPQEIEKASSKVETLNKTEEVKDEKSENEVTPDLKSEEPKSLETKVKEEPKPAVQTPVKEEETGSNIQKTPSFSFNSTT**T**PKSTSST**SS**IFGGGLKTQ**T**PSSSNSTNIFGARTTTTA**T**PTPASNTSSIFGGGSKAASSPFGSFGQAGCQPAKTSNPATSTASVTFSFNTGATSASAKPAGFGSFGAGASAKPSSVFGGSVTAPTVPNVDDGMEDDSMANGGGSGGFMSGLGNARTSNTSGGNNPFAPKTSTGTSASSSSWLFGGGGNQQQQQQQKPSFSFNTAGSSAQQASAPATGTSSVFGGAPKFGSQPAFGAKPFGGGANAGLSKNASIFGGATSSTTNNPATGGFAQFASGQKTSSLFGGGATPQTNTSIFGGGANTTPAPTSSVFGGGASANANKPTSFTSWR

**>NPP-19^NUP53^ isoform b**

MFSHLNQN**T**SGRHNSMDLNNSSISNFGTPVEQSTPALLFGKRKATVPSSYTASPLNTASAPCSDIFAVSAPAVPQHLKDTPGSKSVHWSPSLVQSGEKSAAQTQNTPANLSFGGNSSFSAPTKPAPQSIQTSSFGGQAMHAPPLRSLRDKVEPAKKISRRNTFTARS**T**PLS**T**PITQRVTSRLAEAEEQPMEEEADAADTWVTVFGFQPSQVSILLNLFSRHGEVVSHQTPSKGNFIHMRYSCVTHAQQAISRNGTLLDQDTFIGVVQCTNKDVINGSASGIVARSSNIAAAANRSASMYNSFVENDMADQSVNHNENSVLNSSNVFDANNSLNSSRISVR**S**GVGMRPLAADQRTNILQG**T**PSVRKAPDGLLNKFWNTIGLN

**>NPP-24^NUP88^**
MLITHLIDSLSDRAVITPLTPTSVIIYDKNELKVYVGFTNNRNSLEYTKEVVLLLSTEIPKKAEIREIIVSKNGDYVILEGPRSLFVVRIGAEILVAKPDRLPSECFCECYPLHDSLLLQNISLSVVKVRLLPEKCDEKTFVTAVLFSDNCIRFYNLQKKFDSLLLAVDFRNHLHQVHDENVANN**T**FGLQKALVSFDLIPPKPNTSHFSIISIDSDCDFYTSFVHFSCFKEGYAPRIHRIEPVDGLPCDPLDLRYIQTTNPRICSVFVLVSGGGVLSHLVVFPNEFGEFRFLVKDQLRLPSSNGDPRIVQNQIRSLKVSRYEIATSSSLFSVNIFPWFEALTSISPTSTLEKETRVSELVDAVIPSDELSNTTKWTGARALRAVSVQLTQSLATEEEELLPESENIMHLVILENKDGQPAHLFNISTFDNIWSTENKTSFGRDSVSQPPMKSTGSLEQQLAALKPLAACVISEKVSCEEAIDAAMKFFDAVDERLKKHCEISKLFVERCLAVSSSAQALDEKQQSVDQRLIEETNTVEELKIRMHETKERMEGARKGINVLFHRVDENVPLSDNEIRIFERLKEHQKMLSDMTKLVPKMTLDSNEIHRMANIVLKKRGTGEEQNRFAAVEKNATEIESLEARENKLNTGISELSI

**>MEL-28^ELYS^**

MDNENSSIFKSYQGYECWRGEKQIILKDSIGRQLPYIVNFKKNTCQIFDIEWERVTHSFVFPEGCALIDADYFPTEEGKLGILVGVEDPRQSCGAEHFVLALAVDPDSPAMTITHSLEVPSKITVVKTLFSSADMADETQRTVLKLYHRLMTWQHIVAIGCKETQCYLARLVAVETPSSPVITVHSEKKYLINLMNAYVSGSVLQYTLDDGAYREYPTAAVYISALSLMPRSRTLLVGLSMGGILAASLNPSNQMMLLELRHERLVRKIAPLEPEDDPDKFEYFIATVDCSPRHPIMIQLWRGSFKTLEDVDGEEKYDRPSFSVCLEHKILFGERWLAVNPIVTERDHMMLTRKRGTEDSMHNVSQTFGSTSNRNSVLLAYERKKMVIGTEDPNAEPEYIVEAAIFDIDSWYYKRVPGRVSTDGTVLKQCAFLSTIKSNIRSEDVNDIGILTNEATDVSSFSSMVSDADQLFYPSALSFERVFVAKNTRIDWMKIQNIQDTILNKCAVKLPALIRNPEMISSVVMAAGLVRKNILSGSPNSSAAEINELQLSSDQKVLLNVIVYYGKIEEFCQLASRPDISDTLKRELAEWALHEAVDYKRTISDKMVSLFQGRSLALSPLAEESIAQGIKLFRVVYEYLKACSKALKDDRLRNLAHSVICMRNHTKLTSQFINFAIIPVDPIRQQRMKDLHSKRKNMARKNSSSLPVQSVVRKMNRQAPNAQFWNDIPHDEWYPPTPLDLLECLLNVSISESIKRELVVQYVIDWISTSPEDSEHSEKQLALETIKIMTNQMLNVNLEKIYYILDQGKKALTSSKTSDDMRALGEKVFSMKDDEISYEKLWGKDAPMTVTIGKHDLQRFEQRMKMQMEGGKVRLPVLDPESEILYQMFLFENEKFEAMSSEAISSNKLLSAFLPGMIKKDGRGRQKTAKEQEIEISVKKMFERKVQNDDEDMPEVFASVNDKTERKRKSSQFGEDDESSVSSSQYVPPTAKRIQQWKSAVESVANNSSINSITSPDSHQNAEINMMIATPARYYKRHNEEENVQDGFLSPAGNRPPPVSAHNSILKTAKGGQSASRGRIRFRADVPRGADE**S**IEDNGRKGLALNFAILEDEEEETMTIRKSRSMGKHDEEKDSEKNVVDEMEEVKDQEQENDECIESEKTFENQDDFEVLEDTSAPEAANTENGSETPPMEDTFEVRDDDVMPPTDETYLSHLQTDKTGILEEEGEDEDIWDGVQRSFEVQMDEDCEAVPTIDVADDLESKSEEVNEEEVVESEEVQQDAKEPEKTEKRQEEPEPEVMQPVIPEEPQNESLESSIKLQEELQEEPDIVPTGDEDTADKVQEQAVEEDRPPSRNTRSSSVQKSTSQVEDRDPKELVEEERPPSRNTRSASVQKSSNQEKTSESGEVTEEDRPPSRNTRSASVQKSSSKVKDQKPEELIEEDRPPSRNTRSASAQKTVAANKSVLESEIPSRSASRRTRSTSLRNDTVAEPDETSVAMTTRRRTRAT**S**EVVSKQSSEDDGRSTPKTGRTPTKKAAASTSSSRAGSVTRGKKSIIQKMPSPLEVTMEVQEEEEEEAEEERPASRSTRSASVKNTTVDPSSSALA**S**TKRTTSRKRGNSETIDFNQDDKSAPTTPKRGRPAKKDAGSPKVGSKARGTKPKSIFENQEDEEDRSSSPDIEQPATPTRSSKRTARSRANSESIDDDSKQK**T**PKKKNAAVNEAGTSKQSRSVTRSRASSIDVQQEVEEPT**T**PKRGRGRPPKTVLENIEEGEEERKETAATPLLRSARRAKQ

*Macro generated in Fiji to convert .tif movie to an image sequence of channel 1 compatible with Ilastik and quantify the integral density value at the nuclear envelope.*

roiManager("reset");

run("Options...", "iterations=1 count=1 black do=Nothing");

run("Set Measurements...", "area mean integrated stack display redirect=None decimal=3");

run("Colors...", "foreground=white background=black selection=yellow");

close("*");

run("Image Sequence...");

imagesList = getList("image.titles");

level = getNumber("Threshold between O and 1: ",0.20);

for (i=0; i<imagesList.length-1; i++){Berg et al., 2019, #88830}agesList[i], "Probabilities.tif") == true)

{print("	"+imagesList[i]);

selectImage(imagesList[i]);

run("Duplicate...", "duplicate channels=1");

setThreshold(level, 1000000000000000000000000000000.0000);

setOption("BlackBackground", true);

run("Convert to Mask");

selectImage(imagesList[i]);close();

}

}

run("Images to Stack", "name=Stack title=[Probabilities] use");

//run("Invert LUT");

for (i=0; i<imagesList.length-1; i++){Berg et al., 2019, #88830}af0 setSlice(i+1);

run("Create Selection");

//	run("Make Inverse");

RoiExist = getValue("selection.size");

if (RoiExist > 0)

{

roiManager("Add");

roiManager("Select", i);

//	roiManager("Rename", i);

}
